# There are cores, indicators, and differences – a high-throughput and quantitative PCR study of the *Ambrosia artemisiifolia* L. leaf, root, and seed endophytic bacterial microbiomes within the European invasive range

**DOI:** 10.64898/2026.10.23.761399

**Authors:** Tomasz Grzyb, Michal Hrabovský, Barbara Tokarska-Guzik, Justyna Szulc

**Affiliations:** Department of Environmental Biotechnology, Faculty of Biotechnology and Food Sciences, Lodz University of Technology, Wolczanska 171/173, 90-530 Lodz, Poland; Department of Botany, Faculty of Natural Sciences, Comenius University, Révová 39, 811 02 Bratislava, Slovakia; Institute of Biology, Biotechnology and Environmental Protection, Faculty of Natural Sciences, University of Silesia in Katowice, Jagiellońska 28, 40-032 Katowice, Poland

**Keywords:** Ambrosia artemisiifolia, endophytes, endophytic bacteria, core microbiome, seed microbiome, invasive plants

## Abstract

**Background and aims:** *Ambrosia artemisiifolia* L. is among the most harmful invasive plants in Europe, a major agricultural weed and a source of severe allergenic pollen. Endophytes may affect invasion, but this species’ endophytic bacteria were profiled by high-throughput sequencing only in a single study from China. We aimed to characterize the leaf, root, and seed endobacteriome and test whether tissue type, habitat, or region shape its composition within the European invasive range.

**Methods:** Plants were sampled from six sites in Poland and Slovakia across three habitat types. 16S rRNA gene amplicons from surface-sterilized tissues and soil were sequenced and bacterial load quantified by qPCR. Analyses included alpha and beta diversity, core microbiome, indicator analysis, differential abundance, and functional prediction.

**Results:** Tissue type, not region or habitat, structured communities. *Gammaproteobacteria* exceeded *Alphaproteobacteria* in most plant tissue samples, but in soil the reverse was true. Mean bacterial load in plant tissues ranged from 1.73-3.49 × 10^8^ 16S rRNA gene copies g dry weight^-1^. The pan-tissue core (80% prevalence) contained 13 genera, 9 of them absent from the soil core. Seeds had significantly lower alpha diversity, the smallest core and no seed-exclusive predicted functional indicators. Leaves and roots differed significantly in taxonomic composition but not in predicted functions.

**Conclusion:** Tissues shared a core that overlapped only partly with the soil core. Seeds carried a reduced subset of the endobacteriome that, in this annual invader, may pass across generations. Further studies should compare these communities with native-range populations and test the enhanced mutualism and pathogen spillback hypotheses.

## 1. Introduction

As anthropogenic pressures intensify and the world becomes increasingly interconnected, the number of invasive alien species is growing rapidly, with no signs that the rate of new species introductions or the emergence of new invasive species is slowing down (Pyšek et al. 2020; Guo et al. 2024). Biological invasions have substantial ecological impacts, further raising the risk of biodiversity loss and shifting biogeographical boundaries (Guo et al. 2024). Invasive species also negatively affect human well-being and lead to significant economic losses worldwide (IPBES 2023).

*Ambrosia artemisiifolia* L. (common ragweed) is one of the most harmful invasive plants in Europe, with severe economic and public health consequences. It is a major weed in crop fields, especially in spring-sown wide-row crops (Knolmajer et al. 2024), highly present beside fields as well. Ragweed pollen allergy represents a serious health issue, reaching high sensitization rates; de novo sensitization in adults is frequent, and ragweed pollen induces asthma more frequently than other pollens (Chen et al. 2018). European ragweed epicentre is located in the Pannonian Basin, the Rhône Valley in France, and Northern Italy (Sikoparija et al. 2017; Skjøth et al. 2019), yet it is present in various other countries. It is projected to further expand with climate warming (Knolmajer et al. 2024).

Among different hypotheses in invasion biology, plant-microbe interactions are regarded as a priority in discovering roles of various traits in the invasion process (Gioria et al. 2023). Under the holobiont concept, plants and their associated microbiota function as an integrated eco-evolutionary unit whose fitness depends on interactions between host genotype, microbiome composition and the environment (Vandenkoornhuyse et al. 2015). The net effects of soil microbial communities may determine an invasive plant’s failure or success, depending on the strength of pathogenic (negative) effects as opposed to mutualistic (positive) interactions (as in the enhanced mutualist hypothesis) with microorganisms in the introduced range (Gioria et al. 2023).

In the case of *A. artemisiifolia*, it has been demonstrated that the root-associated strain of *Sphingomonas sanxanigenens* facilitates colonization by arbuscular mycorrhizal fungi (AMF) (Li et al. 2025). A rhizosphere-isolated strain of *Priestia megaterium* also stimulated common ragweed’s growth (Zhang et al. 2023a), and endophytic *Pseudomonas* and *Bacillus* strains improved its drought tolerance (Wu et al. 2026) There are very few studies using high-throughput sequencing to investigate ragweed’s microbiome. Bieker et al. (2020) surveyed the leaf microbiome of selected herbarium, and modern North American specimens using shotgun metagenomics, and in a subsequent, broader-scale study, Bieker et al. (2022) used shotgun metagenomics to detect bacterial pathogens across modern European, modern North American, historical European, and historical North American leaf collections, without distinguishing between epiphytes and endophytes. Zhang et al. (2023b) and Wang et al. (2023) have studied its rhizosphere microbiome, but not the tissue microbiome; Li et al. (2025) have studied its root microbiome, but without distinguishing between epiphytes and endophytes, and Zhou et al. (2025) have studied the phyllosphere fungal microbiome; they did it without distinguishing between epiphytes and endophytes, too. Only Wu et al. (2026) focused on endophytes and studied bacterial seed endophytes from Chinese populations, traced their vertical transmission to seedlings, and described root and leaf endophytes from a common-garden experiment. To our knowledge, there are no more published articles concerning these topics, and besides one study from 2020, all of this research has been published in 2022-2026. Leaf and root endophytes of natural population plants have still not been studied via sequencing. Concerning seeds, there is no sequencing microbiome-related data for the European invasive range.

Despite growing interest in ragweed-associated microbiomes, endophytic bacterial communities remain unexplored to underexplored. Endophytes, defined by their habitat rather than host function (Hardoim et al. 2015), comprise mutualistic, commensal, and antagonistic microorganisms that colonize internal plant tissues and interact more closely with their hosts than epiphytes (Compant et al. 2021; Grzyb and Szulc 2024). Through phytohormone production, nutrient acquisition, modulation of plant immunity, and enhancement of tolerance to biotic and abiotic stresses, they can affect plant fitness (Hardoim et al. 2015; Ameen et al. 2024). Conversely, some endophytes may behave as latent pathogens, pointing out the dynamic continuum between beneficial and detrimental host interactions (Brader et al. 2017; Grzyb and Szulc 2024). Because of their intimate association with plant tissues, endophytes are gradually recognized as potential contributors to invasion success by enhancing host performance, stress tolerance, and establishment in novel environments. However, evidence remains scarce, and only limited studies, primarily on fungal endophytes, have suggested their involvement in plant invasions (Bard et al. 2024).

Defining the core microbiome is recognized as an important challenge in microbial ecology. Identification of a core microbiome is essential for understanding which microbial components remain stable and consistent across complex assemblages and environmental conditions (Shade and Handelsman 2012). The core plant microbiome is considered to comprise keystone taxa that are critical for plant fitness, assembled through evolutionary mechanisms of selection and enrichment (Compant et al. 2019). There is also a growing consensus that the core microbiome should be characterized both taxonomically and functionally (Lemanceau et al. 2017). Prevalence-based core analysis identifies taxa that persist across a defined proportion of samples, but does not assess whether their abundance differs among contexts.

Another approach is indicator species analysis, which identifies taxa significantly and specifically associated with particular habitats or conditions. The method, originally developed for macroecological surveys (De Cáceres and Legendre 2009; De Cáceres et al. 2012) is now being increasingly adopted in plant microbiome research (Carrascosa et al. 2023; Noble et al. 2025; Pawłowicz et al. 2025). Indicator analysis distinguishes taxa that are statistically associated with specific ecological contexts by jointly evaluating fidelity and specificity – a distinction particularly relevant when multiple host compartments are compared simultaneously. Differential abundance analysis (Lin and Peddada 2020) complements both approaches by detecting quantitative shifts in individual taxa between compartments, regardless of their prevalence or indicator status. The three methods combined, applied to the same dataset, provide a more reliable characterization of the microbiome. This framework extends to predicted functional profiling.

This study aimed to characterize the bacterial endophytic microbiomes of leaves, roots and seeds of *Ambrosia artemisiifolia* and assess whether tissue type, habitat and geographic region shape their composition. By integrating metataxonomics with absolute abundance estimates, performing core microbiome, indicator taxa, and differential abundance analyses, and applying functional prediction, we sought to identify bacterial taxa that consistently characterize the endosphere of this globally invasive plant and improve understanding of microbiome assembly across plant compartments.

## 2. Materials and Methods

### 2.1. Site location and sampling

*Ambrosia artemisiifolia* L. plants were sampled from six locations across Poland and Slovakia, representing three habitat types: agricultural maize fields, adjacent meadows, and high-traffic roadsides (**Figure 1a-b**). Two paired sites were established in distinct biogeographical regions, allowing habitat type and geographic region to be examined within a comparable framework. A third set of roadside sites in a separate region was included as a contrasting reference, which differed from the paired sites in both habitat type and geographic context. The first pair was located in Starzawa, southeastern Poland (Podkarpackie Voivodeship), comprising a conventionally managed maize field) and a meadow situated approximately 600 m to the southwest. The second pair was established near Kaplna in southwestern Slovakia (Bratislavský kraj), with a maize field and an adjacent meadow approximately 600 m to the southeast. Two additional populations were sampled from roadside habitats in the Silesian Voivodeship in Poland. The first roadside population was in Jaworzno along the S1 expressway (European route E75). The second was sampled in Kuźnica Warężyńska along national road DK 86, approximately 2 km from its junction with the S1 expressway. Both roadside sites experience high vehicular traffic. Coordinates of individual sampling sites are presented in **Table 1**.

**Table 1.** Coding system for sample types and sampling sites; na – not applicable.

| Component | Code | Description | Coordinates | Sample number |
| --- | --- | --- | --- | --- |
| Plant compartment | L | Leaf | na | 6 |
|  | K | Root |  | 6 |
|  | N | Seed |  | 5 |
|  | G | Bulk soil |  | 6 |
| Sampling site | 1 | Starzawa maize field | 49°52'36.45"N,<br>23°01'18.81"E | 4 |
|  | 2 | Starzawa meadow | 49°52'35.00"N,<br>23°00'48.90"E | 4 |
|  | 3 | Kaplina maize field | 48°17'54.80"N,<br>17°26'50.60"E | 4 |
|  | 4 | Kaplina meadow | 48°17'38.60"N,<br>17°27'06.40"E | 4 |
|  | 5 | Jaworzno roadside | 50°13'02.30"N,<br>19°10'17.60"E | 4 |
|  | 6 | Kuźnica Warężyńska roadside | 50°24'10.20"N,<br>19°11'36.00"E | 3 |

Sampling was conducted in two phases. The Phase 1 sampling (early flowering) was conducted in early August 2024; at each site, 7-10 mature *A. artemisiifolia* individuals (without disease symptoms) were randomly selected to capture within-site heterogeneity, and bulk soil (0-10 cm depth, plant-free spaces), middle-section taproot, and the second-to-fourth leaves from the apex were collected. The Phase 2 sampling (seed maturity) was conducted in October – early November 2024, depending on seed ripeness. Mature seeds (dry, brown achenes) were collected from reproductively mature plants at five sites; seed collection was not possible at Kuźnica Warężyńska, where plants had been mowed by road maintenance authorities. Surface sterilization of plant tissues followed Sahu et al. (2022) using sequential ethanol, NaClO, and sterile distilled water rinses (full protocol in **Methods S1**). Surface-sterilized tissues were preserved in DESS solution (Carvalhais et al. 2022); bulk soil served as an environmental reference and was stored similarly without sterilization. All samples were stored at −80 °C until DNA extraction.

### 2.2. DNA isolation and sample coding

DNA was extracted from DESS-preserved tissues (leaves, roots, seeds; 0.4-0.5 g fresh weight per replicate) using an SDS-based protocol modified from Maropola et al. (2015), and from soil (∼250 mg per replicate) using an adjusted Protocol III by Dwivedi and Yadav (2023). Dry mass was determined (105 °C to constant weight) on parallel sub-samples for absolute abundance normalization. For each biological sample, the three independent extraction replicates were pooled into a single tube to yield one composite DNA preparation per sample, and DNA concentration was measured using a Qubit fluorometer with the dsDNA HS assay. Detailed lysis buffer compositions, bead-milling parameters per tissue type, enzymatic and chemical lysis steps, organic extraction sequences, and precipitation protocols are provided in **Methods S2**. Samples were coded according to the scheme presented in **Table 1**.

**Figure 1a.**
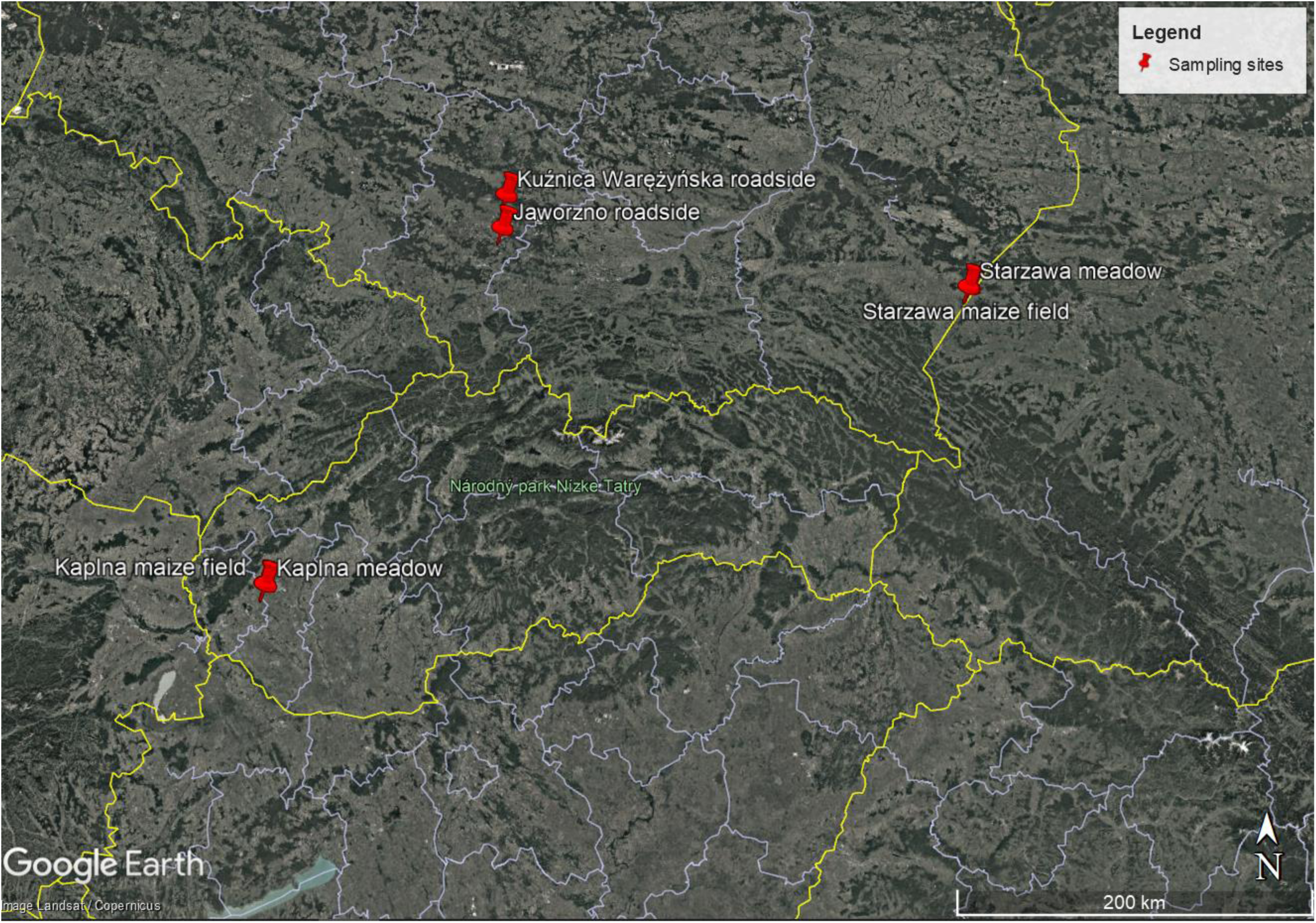
Geographic distribution of Ambrosia artemisiifolia L. sampling sites across Poland and Slovakia. Red pins indicate the six study locations representing three habitat types: paired agricultural (maize field) and semi-natural (meadow) sites in Starzawa (southeastern Poland) and Kaplna (southwestern Slovakia), and two roadside sites in the Silesian Voivodeship (Poland). Map generated in Google Earth Pro; scale bar: 200 km.

**Figure 1b.**
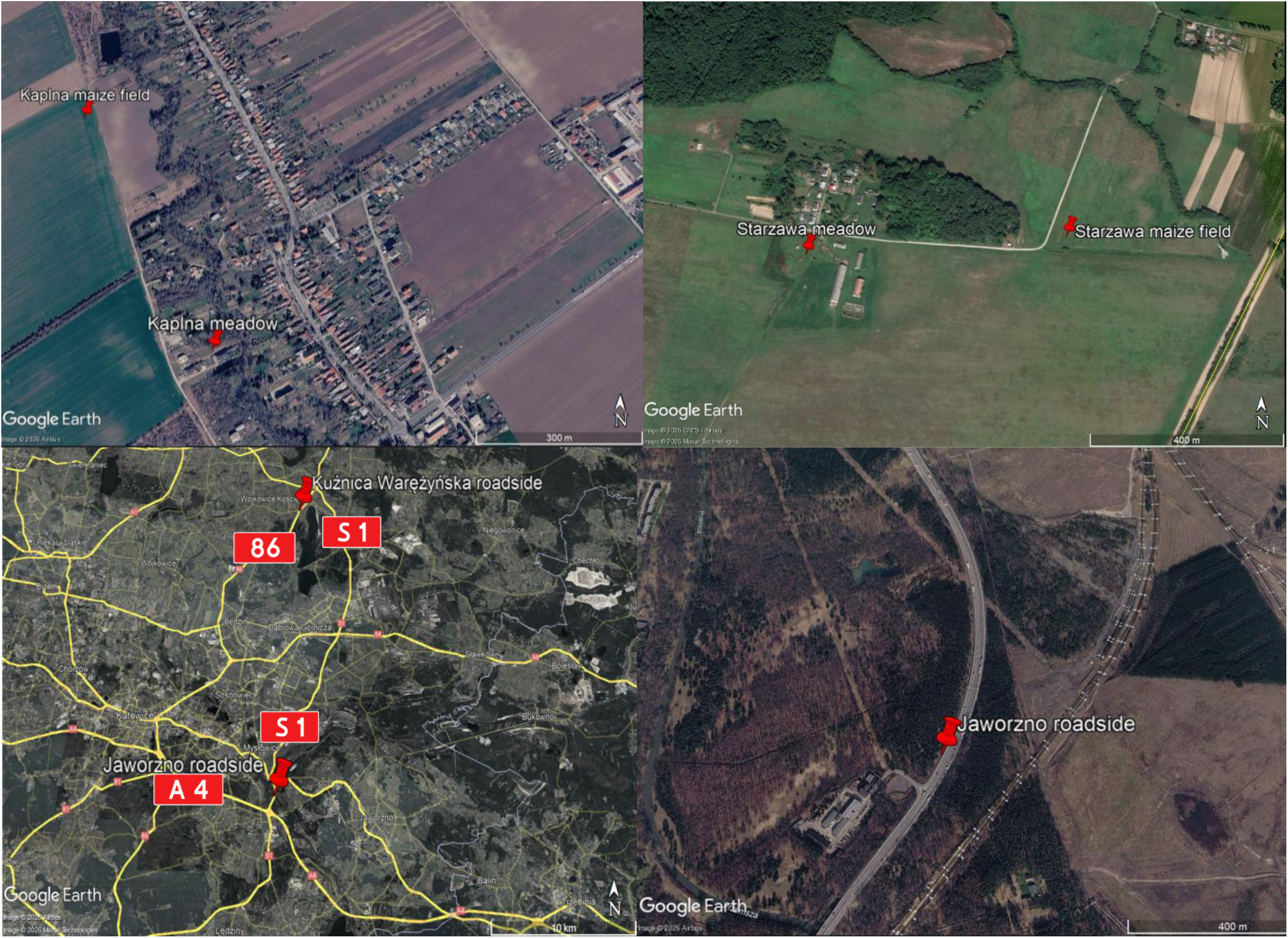
Detailed satellite imagery of individual sampling sites. Top left: Kaplna meadow and maize field sites (southwestern Slovakia); top right: Starzawa meadow and maize field sites (southeastern Poland); scale bars: 400 m. Bottom left: overview of the two Silesian Voivodeshipn roadside sites (Jaworzno and Kuźnica Warężyńska) along the S1 expressway, national road DK 86, and the A4 motorway; scale bar: 10 km. Bottom right: Jaworzno roadside site; scale bar: 400 m. Maps generated in Google Earth Pro.

### 2.3. Primer selection and in silico evaluation

Based on literature research, two candidate primer pairs targeting the bacterial 16S rRNA gene were considered for this study: 799F/1193R (V5-V7 region; 799F: 5’-AACMGGATTAGA-TACCCKG-3’; 1193R: 5’-ACGTCATCCCCACCTTCC-3’) (Bodenhausen et al. 2013), and 322F/796R (V3-V4 region; 322F: 5’-ACGGHCCARACTCCTACGGGA-3’; 796R: 5’-CTAC-CMGGGTATCTAATCCKG-3’) (Chen et al. 2022b). To compare the two pairs, an in silico evaluation was performed using SILVA TestPrime 1.0 (Klindworth et al. 2013) against the SSU RefNR database, with the 3′ mismatch zone disabled and allowing 0, 1, or 2 mismatches (**Table S1a**). The 322F/796R pair yielded higher bacterial coverage than 799F/1193R at 0 and 1 allowed mismatches (65.9% vs. 48.2% and 76.8% vs. 73.4%), with the relative advantage reversing only at 2 mismatches (87.7% vs 90.5%). 799F/1193R showed substantially greater in silico co-amplification of mitochondrial sequences across all thresholds (27.8–59.7% database coverage vs. 8.6–14.1% for 322F/796R). Chloroplast co-amplification coverage remained low for both pairs at 0 mismatches (<1%) but diverged at 2 mismatches (10.7% vs. 6.9%). The 322F/796R pair was selected for all downstream amplicon analyses. Phylum-level in silico coverage for the selected pair is reported in **Table S1b.**

### 2.4. Polymerase chain reaction (PCR) and metataxonomic sequencing

PCR amplification of the bacterial 16S rRNA gene (V3-V4 region) was performed using the abovementioned primers 322F and 796R (Chen et al. 2022b) (full reaction composition and thermocycler program in **Methods S3**). PCR products were verified by agarose gel electrophoresis (2% TBE gel). PCR products were purified using Agencourt AMPure XP magnetic beads and resuspended in Elution Buffer. Library construction, quality assessment (Agilent 2100 Bioanalyzer), and sequencing were performed by BGI Genomics using the DNBSEQ-G400 platform (paired-end, 300 bp reads). Per-sample sequencing yields ranged from 23.4-24.2 Mbp raw data, with >94% clean reads retained after quality filtering.

### 2.5. Quantitative PCR

Bacterial 16S rRNA gene copy number per gram of dry weight was quantified by qPCR using the same 322F/796R primers in triplicate reactions (full protocol in **Methods S4**). A standard curve was generated using serial dilutions of purified *Bacillus cereus* zielonkawy (NCBI Ref-Seq: GCF_043099295.1) genomic DNA. qPCR efficiency was 74-76% across runs, which is within the expected range for long amplicons (∼470 bp) and consistent with published values for amplicons >400 bp (68.7% for ∼440 bp amplicon considered satisfying by Zemb et al. (2020)). While amplicon length inversely correlates with PCR efficiency (Van Holm et al. 2021), this trade-off is acceptable for achieving sufficient amplification and quantification accuracy. R^2^ values were >0.97 for all standard curves. Standard curve equations were applied to calculate the number of 16S rRNA gene copies in all samples. Melting curve analysis was performed from 65 °C to 95 °C with 0.3 °C increments and 5 s hold time to confirm PCR product specificity.

### 2.6. Bioinformatic and statistical analysis

#### 2.6.1. Sequence Processing and Taxonomic Classification

Raw paired-end sequencing reads were processed using QIIME 2 (version 2025.10) (Bolyen et al. 2019). Demultiplexing, primer trimming, and quality filtering were performed by the sequencing provider. Denoising, chimera removal, and merging of paired-end reads were performed using DADA2 (Callahan et al. 2016). Representative sequences were aligned using MAFFT (Katoh and Standley 2013), and a rooted phylogenetic tree was constructed using FastTree (Price et al. 2010) within QIIME 2. Taxonomic classification was performed using a Naïve Bayes classifier (Bokulich et al. 2018) trained on the Genome Taxonomy Database (GTDB) release 226.0 (Parks et al. 2022). The reference sequences were extracted to match the V3-V4 region corresponding to the 322F/796R primer pair and dereplicated in ‘super’ mode. Following taxonomic assignment, ASVs were filtered to remove low-abundance and rare features: ASVs present in fewer than two samples or with fewer than ten total reads across all samples were excluded. This filtering retained 1,271 ASVs for downstream analysis.

#### 2.6.2. Identification and Removal of Contaminant and Organellar Sequences

Potential contaminants were identified based on taxonomic identity and distribution patterns across sample types. ASVs assigned to genera characteristic of human skin and oral microbiota (*Cutibacterium, Finegoldia, Anaerococcus, Lawsonella, Streptococcus, Staphylococcus, Corynebacterium, Kocuria, Micrococcus*) that were consistently detected across plant tissues but largely absent from soil samples were flagged as probable contaminants (Salter et al. 2014); the same procedure was applied to human oral and respiratory tract microbiota (*Gemella, Moraxella_A*), and lactic acid bacteria typically associated with human microbiome, food fermentation, and laboratory environments (order *Lactobacillales*, including families *Aerococcaceae, Lactobacillaceae, Streptococcaceae*, and *Listeriaceae*). Those were removed before downstream analyses. In total, 48 ASVs (2.48% of total reads) were removed, with per-sample loss from 0 to 6.35% (median 2.19%).

Although GTDB classified all retained ASVs as *Bacteria*, all 1,271 representative sequences were verified against the NCBI nucleotide database (nt) using BLASTn (Altschul et al. 1990) with default Megablast parameters, to detect organellar co-amplification. This screening identified eight ASVs of organellar origin, all matching *Asteraceae* organellar genomes at ≥99% identity: seven mitochondrial sequences, of which four were assigned by GTDB to genus CALKXV01 within *Rhodobacteraceae* (*Alphaproteobacteria*), and three remained unclassified beyond domain level – and one chloroplast sequence, assigned as *Cryptobacteroides* (*Bacteroidota*). Together, these ASVs accounted for 9.89% of total sequencing reads (9.87% mitochondrial, 0.02% chloroplast). Organellar contamination was unevenly distributed across sample types (**Table S1c**): no organellar reads were detected in soil samples. The disproportionately high organellar proportion in seed sample N3 (74.78%) was driven almost exclusively by a single mitochondrial ASV (matching CALKXV01). All organellar ASVs were removed together with human-associated contaminants prior to downstream analyses.

#### 2.6.3. Data Import and Integration

Filtered QIIME 2 artifacts (feature table, representative sequences, rooted phylogenetic tree, and taxonomy) were imported into R (version 4.5.2) using the qiime2R package v. 0.99.6 (Bisanz 2018) and converted into a phyloseq object (phyloseq package v. 1.54.0) (McMurdie and Holmes 2013) for subsequent analysis. Sample metadata, including sample type (leaf, root, seed, soil), site (meadow, maize field, roadside), and region (Podkarpackie Voivodeship, Bratislavský kraj, Silesian Voivodeship), were integrated into the phyloseq object. qPCR values were corrected for the contaminant fraction identified in sequencing data: for each sample, the total bacterial load was multiplied by (1 − c), where c is the fraction of sequencing reads assigned to contaminant and organellar ASVs in that sample, to contain only non-contaminant bacterial biomass.

#### 2.6.4. Bacterial load and alpha diversity

Bacterial load (16S rRNA gcn ‧ g of dry weight^-1^) obtained from qPCR (corrected for the contaminant fraction as described in section 2.6.3) was analyzed to assess differences across sample types, regions, and sites. Data were visualized on a log_10_ scale due to the wide range of bacterial loads across sample types. Three grouping factors were tested across analyses: tissue type (leaf, root, seed, soil), region, and site. For tissue type comparisons, all four sample types, including soil, were tested. For region and site effects, analyses were performed both on the full dataset and on a subset excluding soil samples to separately assess patterns within endophytic communities. Alpha diversity was assessed after rarefaction to an even sequencing depth of 5,268 reads per sample, set to the smallest post-filtering library to retain all samples in the analysis (rarefaction was applied to alpha diversity estimation only). Three diversity metrics were calculated: Shannon index, Chao1, and Faith’s PD (phylogenetic diversity) (Faith 1992). Rarefaction curves confirmed that all three metrics reached a plateau below 5,268 reads (**Figure S1**, computed from 50 rarefaction iterations per depth point). Shannon and Chao1 indices were computed using the phyloseq 1.54.0 package, while Faith’s PD was calculated using the picante package v. 1.8.2 (Kembel et al. 2010). For both bacterial load and alpha diversity, omnibus comparisons among groups were performed using Kruskal-Wallis rank-sum tests. Where omnibus tests were significant at p < 0.05, pairwise Wilcoxon rank-sum tests with Benjamini-Hochberg (BH) correction were applied. Root sample K1 was excluded from alpha diversity and all subsequent primary analyses; the rationale and treatment of K1 across analyses are detailed in **Methods S5**.

#### 2.6.5. Quantitative Microbiome Profiling

Absolute taxa abundances were calculated by multiplying relative abundances from contaminant-filtered sequencing data by the corresponding corrected qPCR-derived bacterial loads, following the Quantitative Microbiome Profiling (QMP) approach by Vandeputte et al. (2017), which is equivalent to the Estimated Absolute Abundance qPCR variant approach by Zhang et al. (2017).

#### 2.6.6. General Taxonomic Composition & Visualization

Taxonomic composition was visualized at five hierarchical levels (phylum, class, order, family, genus) using three approaches: relative abundance stacked barplots (taxa <0.5% collapsed into "Other"), and bubble plots showing the union of top-15 taxa by mean relative abundance and top-15 by total absolute abundance per level. K1 was included in composition visualizations to illustrate its position within the broader dataset (**Methods S5**). All visualizations were generated using ggplot2 version 3.5.1 (Wickham 2016) with colour palettes from RColorBrewer version 1.1-3 (Neuwirth 2022).

#### 2.6.7. Beta Diversity Analysis of Relative and Absolute Data

Beta diversity was assessed using both relative and absolute abundance data. For relative abundance metrics, Bray-Curtis dissimilarity (Bray and Curtis 1957) was calculated to assess compositional differences without phylogenetic weighting. Weighted UniFrac distance (Lozupone et al. 2007) was computed to include phylogenetic relationships. For absolute abundance metrics, Euclidean distance on log_10_-transformed absolute abundances was calculated to assess community differences including microbial load. Additionally, Absolute UniFrac (U^A^) distance was computed according to Pendleton & Schmidt (2026), which combines phylogenetic information and absolute abundances. Principal Coordinates Analysis (PCoA) ordinations were generated for each distance metric to visualize community structure. Statistical significance was assessed using PERMANOVA (Anderson 2001) implemented via adonis2 (in vegan v. 2.7-2; (Oksanen et al. 2025)) with 999 permutations, accompanied by PERMDISP (Anderson et al. 2006) tests of multivariate dispersion homogeneity via betadisper (in vegan), and BH-corrected pairwise PERMANOVA among factor levels. Three grouping factors were tested: tissue type, region, and site. For region and site effects, analyses were performed both on the full dataset and on a subset excluding soil samples.

#### 2.6.8. Core Microbiome and Indicator Species Analysis

Core microbiome taxa were identified using the microbiome package v. 1.32.0 (Lahti and Shetty 2017) based on prevalence thresholds. Taxa were considered core members if present in ≥50%, ≥80% or =100% of samples within a defined group, using a detection threshold of 0.01% abundance. Taxa unclassified at the genus level were retained in relative abundance calculations but excluded from core membership lists and downstream outputs. Core analyses were performed separately for each sample type (leaf, root, seed, soil), and globally across all samples. Pantissue core was defined as the intersection of leaf, root, and seed cores at a given prevalence threshold. To distinguish true endophytic core taxa from ubiquitous environmental bacteria, a plant-specific core was defined as taxa present in the plant core but absent from the soil core. Shared and unique taxa among compartments were visualized using Venn diagrams generated with the ggVennDiagram package v. 1.5.7 (Gao and Dusa 2019), with presence in at least one sample of a given group.

Indicator species analysis was performed using the multipatt function from the indicspecies package v. 1.8.0 (De Cáceres and Legendre 2009), using the group-equalized indicator value index (IndVal.g) to identify taxa significantly associated with specific sample groups, while accounting for unequal sample sizes. Statistical significance was assessed using 999 permutations, with p-values adjusted using BH correction significance threshold: adjusted p < 0.05. Analyses were performed at all taxonomic levels from phylum to genus.

#### 2.6.9. Differential abundance

Taxonomic differential abundance was assessed using ANCOM-BC2 (Lin and Peddada 2020), with sample type as the grouping factor. Genera unclassified at the genus level were retained in the abundance matrix to preserve compositional structure during the model’s bias correction step but excluded from the reported output. Genera below the prevalence threshold of 10% of samples were excluded by the internal ANCOM-BC2 filter, and structural zeros were modelled explicitly. Omnibus tests for differential abundance across all groups were performed with p-values adjusted using the BH method (p < 0.05). Pairwise comparisons were performed only for genera identified as significant by the global test, with BH correction applied independently per genus. Significant genera were aggregated to the class level for the direction summary, and the enrichment direction (which compartment harboured higher abundance) was reported.

#### 2.6.10. Functional prediction

Functional potential of bacterial communities was predicted using PICRUSt2 (Douglas et al. 2020). Representative sequences were exported from the QIIME 2 artifacts to FASTA format, while the contaminant-filtered feature table was exported from the phyloseq object to BIOM format. Both inputs were processed through the PICRUSt2 pipeline (picrust2_pipeline.py) with default parameters. Two ASVs were excluded by the pipeline based on the default minimum alignment proportion threshold (min_align = 0.8); these ASVs were detected in 8 of 23 samples, with a combined mean relative abundance of 0.09%, indicating negligible contribution to predicted functional profiles. No ASVs exceeded the default weighted NSTI cut-off (2.0); per-sample weighted NSTI values ranged from 0.012 (N4) to 0.286 (G5) (mean 0.107; full values in **Table S2**). Predicted KO abundances were mapped to plant growth-promoting trait (PGPT) categories using the PLaBAse database (Patz et al. 2021) at hierarchical Levels 2 to 5, with abundances aggregated by summation within each level. All functional analyses were performed at PGPT Level 5 resolution. Beta diversity was assessed using Bray-Curtis dissimilarity with PERMANOVA, PERMDISP, and pairwise testing performed as in section 2.6.7. Functional core features were identified using the prevalence framework described in section 2.6.8; pan-tissue and plant-specific functional cores were defined analogously to their taxonomic counterparts. Indicator features were identified using the prevalence and IndVal.g approach described in section 2.6.8., with results aggregated to Level 4 for biological interpretation. Functional differential abundance was assessed using ANCOM-BC2 following the method described in section 2.6.9, with results mapped back to their parent Level 3 and Level 2 categories, and enrichment direction summarized per category.

#### 2.6.11. Concordance between taxonomic and functional community structure

Concordance between taxonomic and functional community structure was assessed on the K1-excluded primary dataset (n = 22) for taxonomic and PGPT Level 5 functional profiles. Mantel test (Mantel 1967) with Spearman rank correlation and 999 permutations was performed via mantel (in vegan) to compare the taxonomic distance matrix against each functional distance matrix. Procrustes superimposition with associated permutation-based significance testing (PROTEST) (Peres-Neto and Jackson 2001) was performed via protest (in vegan) with 999 permutations on paired k = 2 PCoA ordinations, assessing the geometric correspondence between taxonomic and functional ordinations.

Scientific names at all taxonomic ranks are set in italics, as postulated by Thines et al. (2020).

## 3. Results

### 3.1. Quantitative metataxonomics

#### 3.1.1. Total bacterial load and alpha diversity

**Figure 2.**
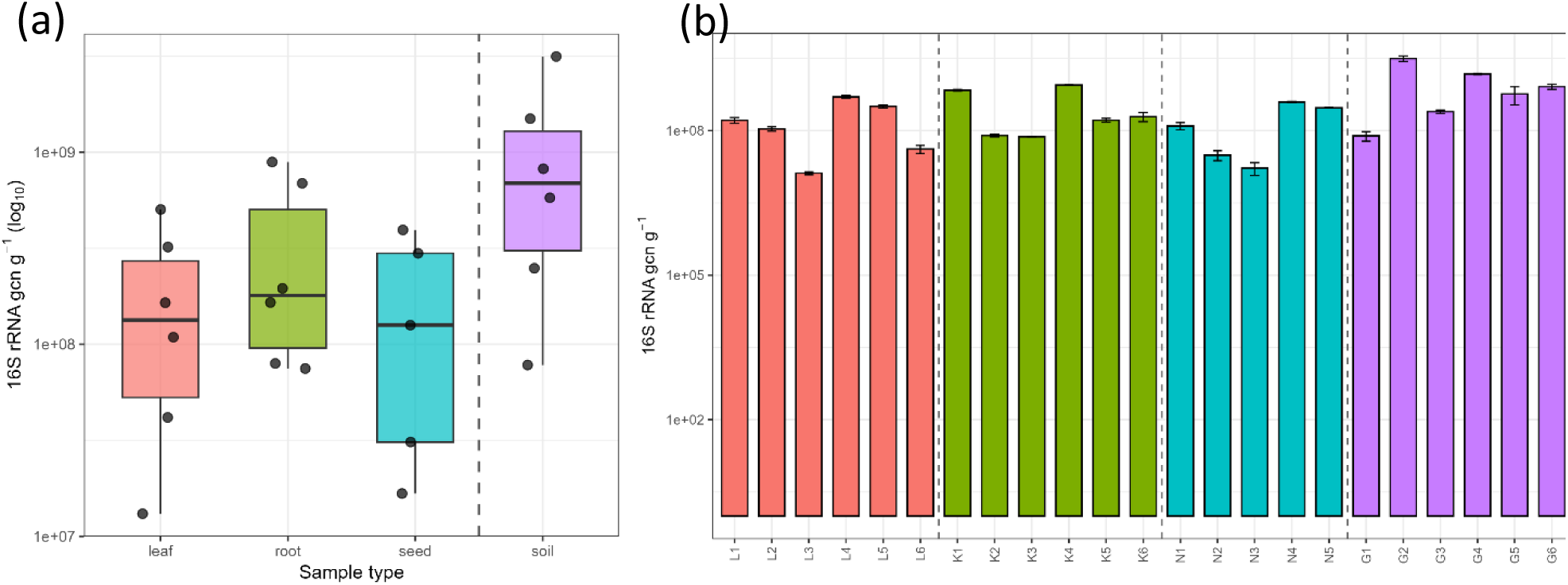
Total bacterial load (16S rRNA gene copies g^-1^ dry weight) across sample types and individual samples. (a) Boxplots with individual data points showing the distribution of qPCR-derived bacterial loads by sample type (n = 6 for leaf, root, and soil; n = 5 for seed). The soil compartment is separated by a dashed line to distinguish the environmental reference from endophytic tissues. (b) Per-sample bacterial loads (bars) with standard deviations from triplicates (error bars). Both panels are displayed on a log_10_ scale. Bacterial loads were corrected for the contaminant fraction identified during sequencing-based contaminant removal (see section 2.6.2). Samples are ordered by tissue type (leaf, root, seed, soil).

Mean absolute bacterial abundance followed the trend: soil (1.06 × 10^9^ 16S rRNA gcn ‧ g of dry weight ^-1^ > root (3.49 × 10^8^) > leaf (1.92 × 10^8^) > seed (1.73 × 10^8^), although differences among sample types were not statistically significant (Kruskal-Wallis, p = 0.141). Likewise, neither geographic region nor sampling site significantly affected bacterial load (all p > 0.12; **Table S3b**). The broad range of observed values combined with high within-group variability may have contributed to the lack of statistical significance. Because the primer pair used underrepresents several soil-associated bacterial phyla (**Table S1b**), bacterial abundance in soil is likely underestimated. qPCR-derived bacterial loads were used to calculate absolute taxon abundances for quantitative microbiome profiling (QMP).

Alpha diversity differed significantly among sample types according to all three metrics (Shannon: p = 0.017; Chao1: p = 0.021; Faith’s PD: p = 0.012) (**Table S4b**). Pairwise comparisons indicated that significant differences were exclusively related to **seed communities** (twice against leaves and once against roots and soil; significant p = 0.024 - p = 0.035), which had significantly lower richness and phylogenetic diversity than leaves, roots, or soil (**Table S4c**). No significant differences were detected among leaves, roots, and soil. Sensitivity analyses including all samples gave the same qualitative conclusions.

Seed endophytic communities showed the lowest mean diversity across all 3 metrics (**Figure S2, Table S4a**). High among-sample variance fell into two patterns: samples N1, N2, and N3 showed diversity values approaching those of other compartments, whereas N4 and N5 were near-monodominant communities. Leaf, root, and bulk soil communities were not significantly differentiated from one another by any metric. Because alpha diversity estimates for soil were probably underestimated by both the rarefaction depth (5,268 sequences) and primer (**Table S1b**) bias against several soil-associated phyla, comparisons involving soil should be interpreted cautiously. Neither region nor site had a significant effect on any alpha diversity metric, whether tested on the full dataset or on endophytic tissues only (all p ≥ 0.13; **Table S4b**).

#### 3.1.2. General Taxonomic Composition & Visualization

Taxonomic composition was visualized at the phylum, class, order and genus levels using relative-abundance stacked barplots and bubble plots, and absolute-abundance bubble plots (**Figure 3** and **Figures S3-S6)**. Complete per-sample visualizations data for all taxonomic ranks are provided in **File S1**. Compositional profiles grouped primarily by sample type rather than by geographic origin or collection site.

**Figure 3.**
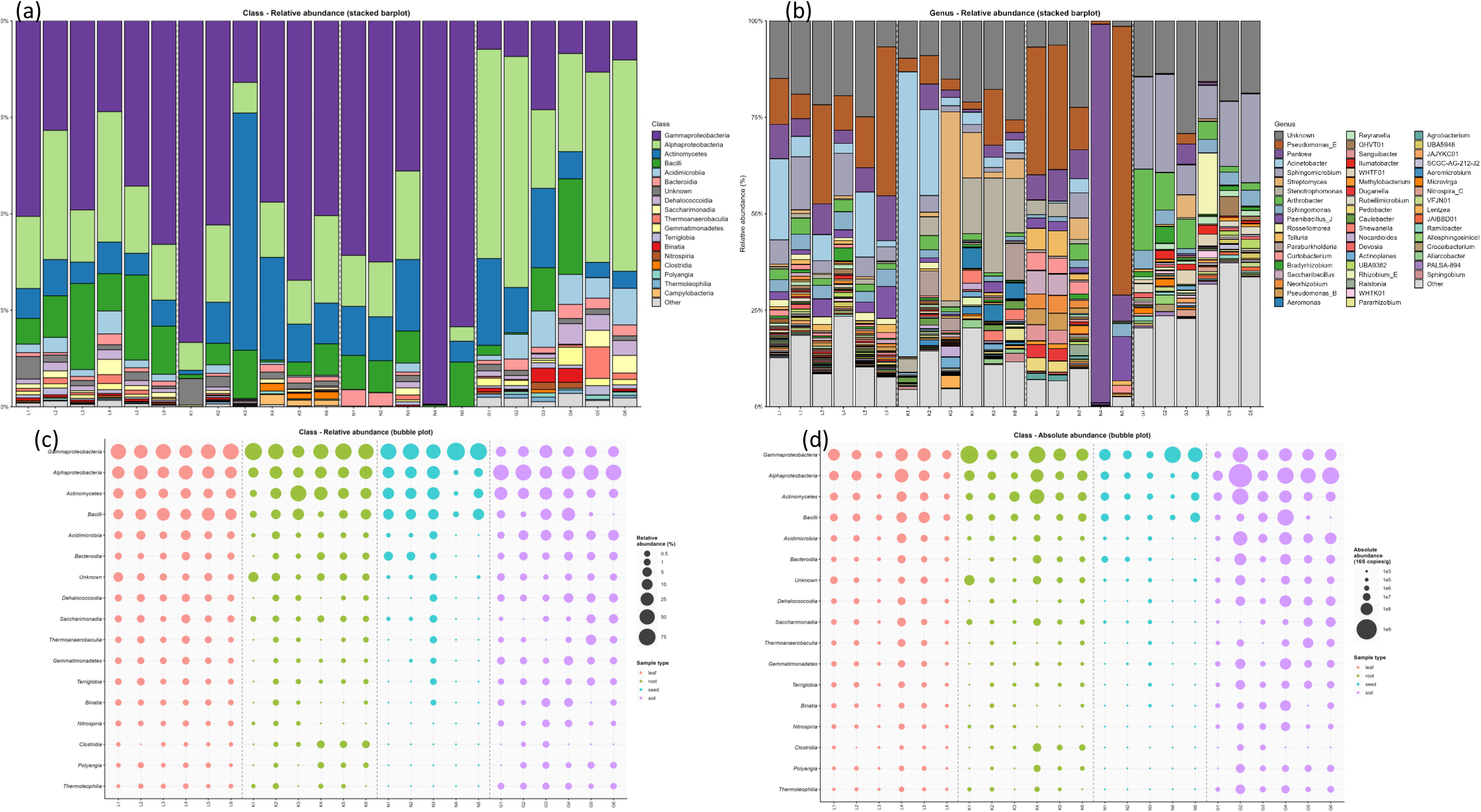
Bacterial community composition at the class and genus levels. (a) Relative abundance stacked barplot at the class level; (b) relative abundance stacked barplot at the genus level. Taxa with mean relative abundance ≥0.5% in at least one sample type are displayed; remaining taxa are collapsed into "Other". (c) Relative abundance bubble plot at the class level (%; square-root-scaled bubble size); (d) absolute abundance bubble plot at the class level (16S rRNA gene copies ‧ g of dry weight^-1^; square-root-scaled bubble size). Bubble plots show a union of the top 15 classes by mean relative abundance and the top 15 by total absolute abundance. Samples are ordered by tissue type and sample number.

At the phylum level, *Pseudomonadota* dominated all four sample types (**Figures S3–S4, File S1**). The second most abundant phylum differed among sample types, with *Bacillota* in leaves and *Actinomycetota* in roots, seeds, and soil. Soil contained several characteristic phyla, including *Acidobacteriota, Chloroflexota, Bacteroidota* and *Gemmatimonadota*. *Acidobacteriota, Chloroflexota and Patescibacteria* were potentially underrepresented, whereas *Verrucomicrobiota, Planctomycetota* and *Cyanobacteriota* were absent, in line with the limited coverage of the 322F/796R primer pair (**Table S1b**). *Campylobacterota* occurred almost exclusively in roots, particularly in samples K4, K5, and K6.

Plant tissues were dominated by ***Gammaproteobacteria***, whereas ***Alphaproteobacteria*** dominated in soil (**Figure 3a,c, File S1**). Leaves showed the most stable class-level profiles, while roots and seeds showed considerably greater between-sample variability. The most pronounced deviation occurred in root sample K3, where *Actinomycetes* became dominant, whereas two leaf samples (L2 and L4), both collected from meadows, showed a relative dominance of *Alphaproteobacteria* over *Gammaproteobacteria*. Beyond the dominant *Pseudomonadota* classes, *Bacilli* ranked third in leaves, *Actinomycetes* second in roots, while both *Actinomycetes* and *Bacilli* contributed to seed communities.

Seed communities were dominated by *Gammaproteobacteria* in all samples, although their relative abundance ranged from near-complete dominance in N4 (99.3%) to 38.9% in N3, where *Alphaproteobacteria* (22.9%) and *Actinomycetes* (18.5%) were also abundant. *Bacilli* were subordinate (0.4-11.6%), reaching second only in N5.

In leaves, *Gammaproteobacteria* dominated in four of six samples (42.8-57.9%), but in two samples *Alphaproteobacteria* (33.5% and 33.8%) exceeded *Gammaproteobacteria* (28.4% and 23.5%, respectively). Both L2 and L4 originated from meadow habitats in different regions (Podkarpackie Voivodeship and Bratislavský kraj). *Bacilli* showed considerable variability, reaching 22.0-22.3% in L3 and L5 but remaining at 6.6-12.5% in the remaining samples. *Actinomycetes* were repeatedly the third or fourth class (5.5-9.4%).

In roots (K1 excluded), *Gammaproteobacteria* dominated in four of five samples, but with the highest variability in subordinate classes. Three samples (K2, K5, K6) followed the pattern *Gammaproteobacteria* > *Alphaproteobacteria* > *Actinomycetes*, while in K4 *Actinomycetes* moved *Alphaproteobacteria* into third position. The most extreme deviation was K3, where *Actinomycetes* accounted for 61.5% of reads, above *Gammaproteobacteria* and *Alphaproteobacteria* entirely; within *Pseudomonadota*, however, the *Gammaproteobacteria* > *Alphaproteobacteria* ratio was preserved even in this sample. The high between-sample variability in roots and seeds contrasted with relatively more stable leaf profiles, while soil communities showed the lowest between-sample variance.

At the order level (**Figure S5, File S1**), each compartment showed a structure with variable internal hierarchies. Seed communities were generally dominated by *Pseudomonadales* (37.2-70.3%), except for N4, where *Enterobacterales* accounted for 98.1% of reads (driven by *Pantoea*), and N3, which exhibited a more even composition involving *Pseudomonadales, Entero-bacterales, Actinomycetales, Rhizobiales*, and *Sphingomonadales*. *Burkholderiales* and *Paenibacillales* were secondary taxa in several seed samples (7.9-12.5%). In leaves, *Pseudomonadales* were the first or second in order (12.0-41.2%), with *Sphingomonadales* as a notable secondary group, particularly in L2 (19.2%) and L4 (17.2%). *Enterobacterales* (3.9-12.1%), *Rhizobiales* (4.7-11.9%), and *Burkholderiales* (2.8-5.3%) were stable components. *Bacillales (B)* reached 17.2% in L5 but remained below 6% in all other leaf samples. In roots, the order-level profile was the most variable: *Pseudomonadales* ranged from 5.4% to 30.0%, *Enterobacterales* from 2.5% to 20.7%, *Xanthomonadales* from 1.6% to 24.9%, and *Streptomycetales* from 1.2 to 48.9%, with no two root samples sharing the same dominance hierarchy. *Sphingomonadales* (2.6-7.7%) and *Actinomycetales* (2.4-5.9%) were secondary taxa across root samples.

At the genus level (**Figure 3b, Figure S6, File S1**), unclassified genera constituted, on average, 17.8% in leaves, 16.4 % in roots, and 18.7 in soil, but were markedly lower in seeds on average (7.4 %), with N3 as an exception (22.4%) and N4 virtually devoid of unclassified reads (0.07%).

Seed samples presented a gradient of taxonomic evenness (**File S1**). N4 was near-monodominant, with ***Pantoea*** accounting for 98.1% of reads, whereas N3 had the highest taxonomic evenness and the largest number of genera above the 1% threshold. N1 and N2 showed highly similar profiles dominated by ***Pseudomonas (E)***. N5 was dominated by *Pseudomonas (E)* (69.6%), with *Paenibacillus (J)* (11.4%) and *Pantoea* (6.8%) as secondary elements. *Pseudomonas (E)* was present in all five seed samples, ranging from dominant (69.6% in N5) to co-dominant (11.0% in N3), while *Pantoea* was ubiquitous but spanned abundances from 98.1% to 6.5%.

Leaf samples showed more consistent genus-level profiles than roots or seeds (**Figure 3b, File S1**). ***Pseudomonas (E)*** was present above 1% in all six samples (6.3-38.6%), dominating or co-dominating in all cases. ***Sphingomicrobium*** (3.3-14.0%) and *Sphingomonas* (2.8-4.7%) were detected across all leaf samples. *Acinetobacter* was highly variable (1.6-21.0%). *Arthrobacter, Pantoea*, and *Paenibacillus (J)* were detected above 1% in multiple leaf samples. *Rossellomorea* was a repeated minor group (1.4-2.1% in five of six samples). *Stenotrophomonas* was present but at low abundance (max 1.3%), contrasting with its dominance in roots. Two samples (L2 and L4) harboured the highest number of genera exceeding 1% relative abundance among leaf samples.

Root samples showed the highest between-sample genus-level heterogeneity of any compartment (**File S1**). ***Stenotrophomonas*** dominated (14.1-24.6%) three samples (K4, K5, K6), followed by *Streptomyces* (11.8%) in K4, *Pseudomonas (E)* (14.5%) in K5, and *Paraburkholderia* (9.5%) in K6. K2 was dominated by *Acinetobacter* (22.2%), followed by *Pseudomonas (E)* (7.4%) and *Pantoea* (6.6%). K3 was dominated by *Streptomyces* (48.9%), followed by *Lentzea* (3.2%) and *Paraburkholderia* (3.2%). *Paraburkholderia* was stably present in roots (3.2-9.5%). *Aliarcobacter* (*Campylobacterota*) was detected above 1% in K4, K5, and K6 (1.1-2.1%), accompanied by *Aeromonas* (4.0-5.3%) and *Shewanella* (2.7-3.3%).

Quantitative microbiome profiling revealed that relative and absolute perspectives diverged substantially (**Figure 3c-d, File S1**). For instance, *Gammaproteobacteria* accounted for only 11.9% of soil reads but reached a mean absolute abundance of 1.07 × 10^8^ copies g of dry weight^-1^ – exceeding their absolute load in leaves (6.68 × 10^7^ copies ‧ g of dry weight^-1^), where they represented 42.1% of the community. Another pattern was observed for *Alphaproteobacteria*, which were proportionally less abundant in leaves than in soil (21.9% vs 44.0%) but exhibited an order-of-magnitude higher absolute abundance in soil (5.16 ± 6.93 × 10^8^ vs 5.02 ± 6.21 × 10^7^ copies ‧ g of dry weight^-1^). Compositional dominance did not directly relate to numerical dominance when total bacterial load differed substantially across compartments. However, within-compartment standard deviations at higher taxonomic ranks frequently exceeded the means, precluding precise between-compartment comparisons of absolute abundance.

#### 3.1.3. Beta diversity

Beta diversity was assessed using four distance metrics. PCoA ordinations were computed on the K1-included dataset to visualize all samples, whereas all statistical tests were performed on the K1-excluded dataset (n = 22). Omnibus PERMANOVA and PERMDISP results were concordant between the two datasets (**Table S5d-f**).

The Bray-Curtis ordination (**Figure 4a**) explained the clearest compartment-level structure despite the relatively low variance explained by the first two axes (PC1 18.1%, PC2 14.0%). Soil formed a distinct cluster, roots were almost the only one with negative PC2, whereas seeds were shifted the most towards the positive PC1 region. In contrast, the Weighted UniFrac ordination (**Figure 4b**) and Absolute UniFrac (**Figure 4d**) showed little clustering by sample type, site or region, although in the latter soil tended to separate along negative PC1. Euclidean log10 ordination (**Figure 4c**) separated the two Silesian soil samples (G5, G6) along PC1 and roots K4– K6 along PC2.

**Figure 4.**
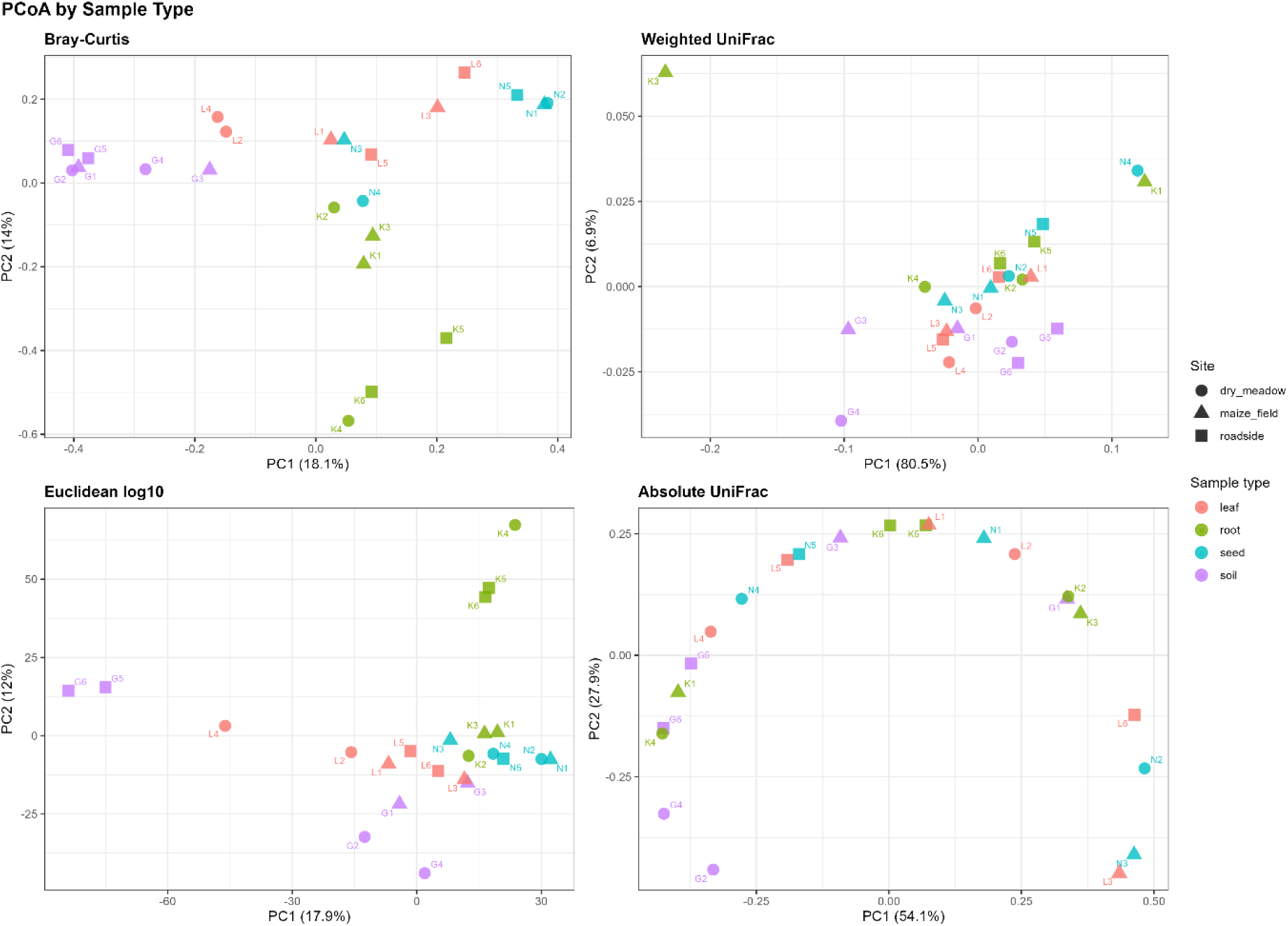
Principal coordinates analysis (PCoA) ordinations of bacterial community composition based on four distance metrics. (a) Bray-Curtis dissimilarity (relative abundance), (b) Weighted UniFrac distance (relative abundance, phylogeny-weighted), (c) Euclidean distance on log_10_-transformed absolute abundances, (d) Absolute UniFrac distance (absolute abundance, phylogeny-weighted). Points are coloured by sample type and shaped by site. Sample labels are shown for identification.

PERMANOVA revealed significant differences among sample types for Bray-Curtis (p = 0.001) and Euclidean log_10_ (p = 0.001), but not for Weighted UniFrac (p = 0.292) or Absolute UniFrac (p = 0.272) (**Table S5a**). A significant regional effect was detected only for Weighted UniFrac (p = 0.035), and disappeared when soil samples were excluded (p = 0.337), whereas sampling site had no significant effect for any metric (all p > 0.09). PERMDISP indicated homogeneous multivariate dispersions for all analyses except Euclidean log_10_ across sample types (p = 0.001), indicating that the corresponding PERMANOVA result should be interpreted with caution (**Table S5b**).

Pairwise PERMANOVA showed **significant differences among all sample-type pairs** for Bray-Curtis (adjusted p = 0.006-0.010, R^2^ = 0.228-0.304). Euclidean log_10_ produced the same pattern (adjusted p = 0.008-0.023, R^2^ = 0.149-0.272), although these results should be interpreted in light of the significant PERMDISP. Inclusion of K1 affected only four pairwise comparisons close to the significance threshold, but the conclusions remained unchanged (**Table S5c**).

#### 3.1.4. Core Microbiome and Indicator Species Analysis

Core microbiome analysis at the genus level identified taxa consistently present across samples within each compartment at three prevalence thresholds (50%, 80%, and 100%; detection threshold 0.01% relative abundance). The number of core genera visibly decreased with increasing stringency across all compartments (**Figure 5a**): leaves 132/74/47, roots 114/62/28, seeds 28/14/9, soil 111/37/16. Seeds harboured the smallest core at all thresholds, compatible with their low alpha diversity and high compositional variance. The pan-tissue core, defined as the intersection of leaf, root, and seed cores, comprised 24 genera at 50%, **13 at 80%**, and **8 at 100%** prevalence (**Figure 5a-b, Table S6**).

**Figure 5.**
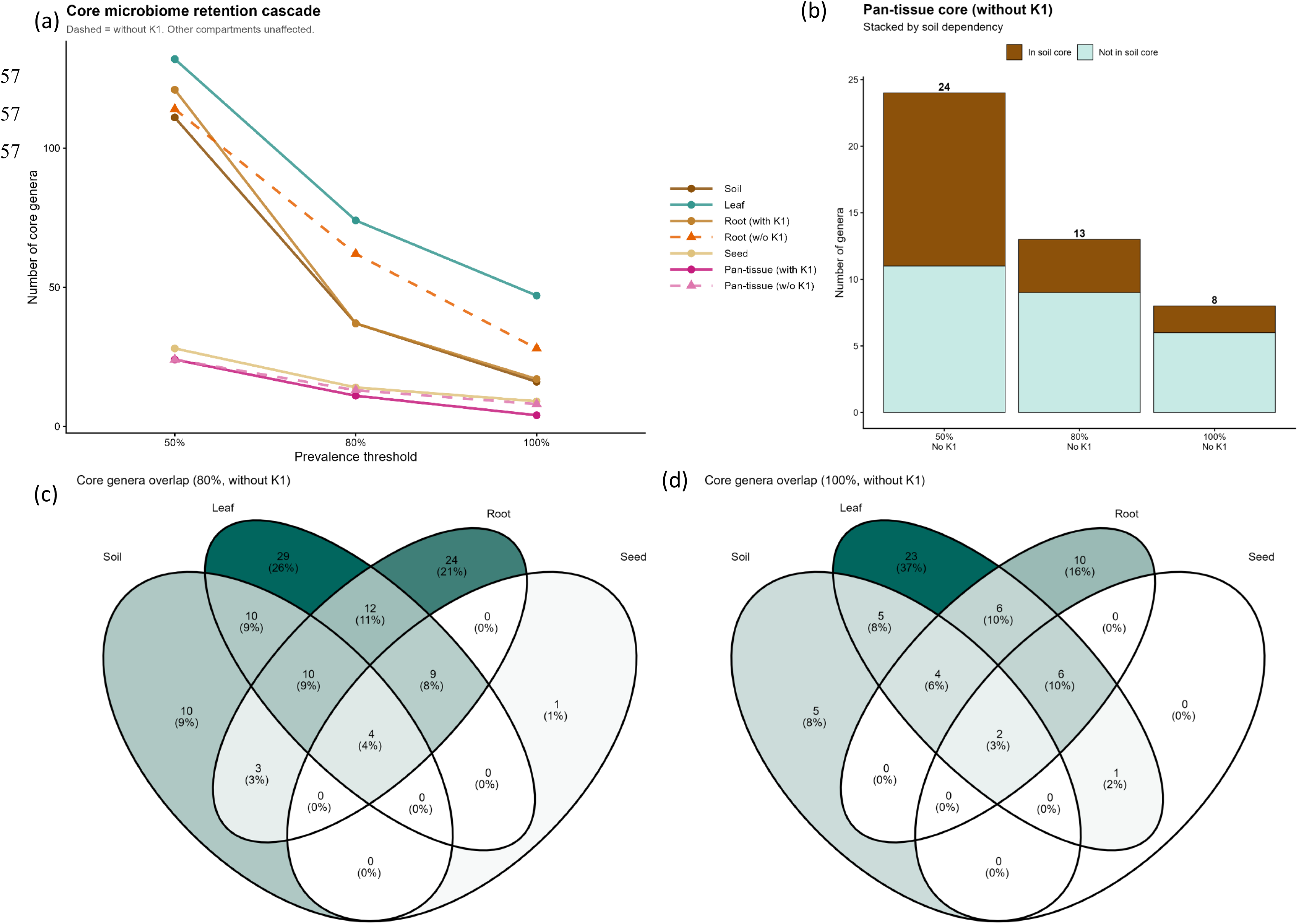
Core microbiome analysis (primary dataset). (a) Core microbiome retention cascade showing the number of core genera at 50%, 80%, and 100% prevalence thresholds for each compartment and the pan-tissue intersection. Dashed lines indicate comparisons with the K1-included dataset for root and pan-tissue; all other compartments were unaffected by K1 exclusion. (b) Pan-tissue core genera at 50%, 80%, and 100% prevalence, partitioned by soil core membership (brown = present in soil core at the same threshold; teal = absent from soil core). Numbers above bars indicate total pan-tissue core size. (c) Venn diagram showing overlap of 80% prevalence core genera among leaf, root, seed, and soil compartments. (d) Venn diagram at 100% prevalence.

To distinguish ubiquitous plant-associated taxa from those shared with soil, the pan-tissue core was further partitioned according to soil core membership (**Figure 5b**).The proportion of soil-core members decreased with increasing stringency: 13/24 were soil core members at 50% prevalence, 4/13 at 80% (*Pseudomonas (E)*, *Sphingomonas, Methylobacterium, Devosia)* and 2/8 at 100% (*Pseudomonas (E)*, *Sphingomonas*). The plant-specific complement – pan-tissue genera absent from the soil core – therefore comprised 11 genera at 50%, 9 at 80% (*Pantoea, Paenibacillus (J), Telluria, Neorhizobium, Pseudomonas (B), Acinetobacter, Curtobacterium, Ralstonia, and Kineococcus*), and 6 at 100% prevalence (*Pantoea, Paenibacillus (J), Telluria, Neorhizobium, Pseudomonas (B), Methylobacterium*) (**Table 2**; **Table S7**). The 13 pan-tissue core genera at 80% prevalence are detailed in **Table 2**, with their abundance and prevalence across all four compartments, soil core membership, and indicator species status.

**Table 2.** Pan-tissue core endophytic genera at 80% prevalence. Mean relative abundance (Abund. %) and prevalence (Prev. %) are shown for each compartment. Soil core: membership in the soil 80% prevalence core (Y = yes). Indicator: significant indicator species association pattern (L = leaf, R = root, Se = seed).

| Genus | Root |  | Leaf |  | Seed |  | Soil |  | Soil core | Indicator |
| --- | --- | --- | --- | --- | --- | --- | --- | --- | --- | --- |
|  | Abund. % | Prev. % | Abund. % | Prev. % | Abund. % | Prev. % | Abund. % | Prev. % |  |  |
| *Pseudomonas* (E) | 6.0 | 100 | 17.5 | 100 | 29.3 | 100 | 0.6 | 100 | Y |  |
| *Sphingomonas* | 1.1 | 100 | 3.8 | 100 | 1.5 | 100 | 3.5 | 100 | Y |  |
| *Methylobacterium* | 0.3 | 100 | 0.4 | 100 | 0.9 | 100 | 0.4 | 83.3 | Y |  |
| *Pantoea* | 2.9 | 100 | 7.1 | 100 | 25.4 | 100 | 1.0 | 33.3 |  | L+R+Se |
| *Paenibacillus* (J) | 0.9 | 100 | 3.5 | 100 | 3.7 | 100 | 0.1 | 16.7 |  | L+R+Se |
| *Telluria* | 0.5 | 100 | 1.2 | 100 | 3.0 | 100 | 0.3 | 66.7 |  |  |
| *Neorhizobium* | 0.3 | 100 | 0.7 | 100 | 2.2 | 100 | 0.1 | 50 |  | L+R+Se |
| *Pseudomonas* (B) | 0.3 | 100 | 0.6 | 100 | 2.2 | 100 | 0.03 | 33.3 |  | L+R+Se |
| *Devosia* | 0.2 | 100 | 0.4 | 83.3 | 0.2 | 80 | 0.6 | 83.3 | Y |  |
| *Acinetobacter* | 6.7 | 100 | 9.0 | 100 | 0.9 | 80 | 0.1 | 66.7 |  |  |
| *Curtobacterium* | 0.3 | 80 | 1.1 | 100 | 2.7 | 80 | 0.1 | 16.7 |  |  |
| *Ralstonia* | 0.3 | 80 | 0.2 | 100 | 0.6 | 80 | 0.3 | 66.7 |  |  |
| *Kineococcus* | 0.1 | 80 | 0.2 | 100 | 0.3 | 100 | 0 | 0 |  | L+R+Se |

At 80% prevalence, the largest shared subsets outside the pan-tissue core were leaf+root (12 genera), leaf+root+soil (10), leaf+soil (10), and root+soil (3) (**Figure 5c-d**). Single-compartment cores included 29 leaf-only, 24 root-only, 10 soil-only, and one seed-only genus, whereas all remaining intersections involving seeds were empty. At 100% prevalence, all shared subsets shrank accordingly. The only notable change was the appearance of *Kineococcus* in the leaf+seed intersection, reflecting its loss from the root core at this stricter threshold.

Indicator species analysis using the group-equalized IndVal.g index identified **28 genera** significantly associated with specific sample-type combinations in the primary dataset (**Table S8a-b, Table S9**). No significant indicator taxa were detected for region or site at any taxonomic level (**Table S8a**). The dominant association pattern was leaf+soil (10 genera, 35.7%; mean IndVal.g = 0.92), comprising predominantly uncultured or poorly characterized lineages (**Table S9**). The next two most frequent patterns each consisted of five genera: leaf+root+seed (*Pseudomonas (B)*, *Paenibacillus (J)*, *Pantoea*, *Neorhizobium*, and *Kineococcus*) and leaf+root+soil (*Reyranella*, *Sphingomicrobium*, *Arthrobacter*, *Bradyrhizobium*, and *Chryseolinea*). Four genera (*Ferrovibrio*, *Actinoplanes*, *Kribbella*, and *Agromyces*) were root-specific indicators, whereas the remaining indicators represented the leaf+root (*Brachybacterium*), root+soil (*Rhizobium (E)*), seed (*Sphingomonas (R)*), and soil (*Gaiellasilicea*) patterns (**Table S8c, Table S9**). Mapping indicator taxa onto the pan-tissue core showed that the leaf+root+seed group contained four members of the 100% plant-specific pan-tissue core and one member of the corresponding 80% core, whereas *Telluria*, despite belonging to the 100% plant-specific core, was not identified as an indicator. Inclusion of K1 resulted in 30 significant indicator genera with only minor redistribution among association patterns (**Table S8d**).

Core membership and indicator species status across prevalence thresholds are combined and summarised in Table 3. At 80% prevalence, the combined plant core (102 genera) contained 26 of the 28 significant indicators; only VFMC01 (leaf+soil, no core) and *Gaiellasilicea* (soil indicator, soil core) fell outside the plant core. The leaf core category contributed the largest number of indicator genera (9) followed by the leaf+root (6) and the root-only core (5). At 100% prevalence, the plant core retained only 18 of 28 indicators.

**Table 3.**
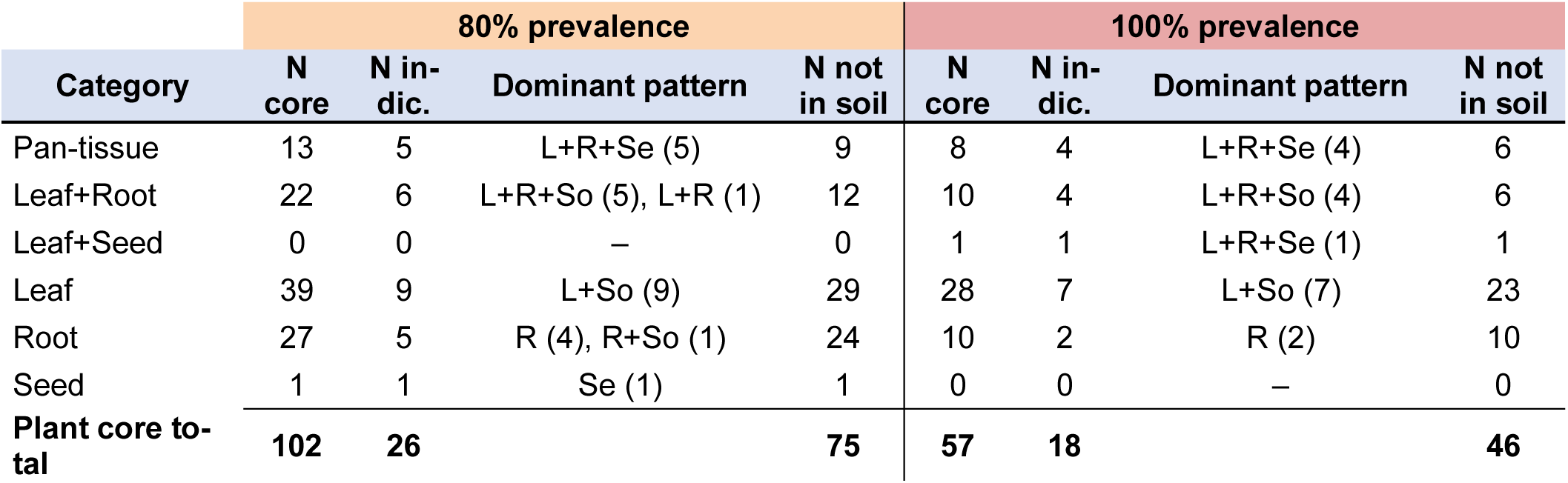
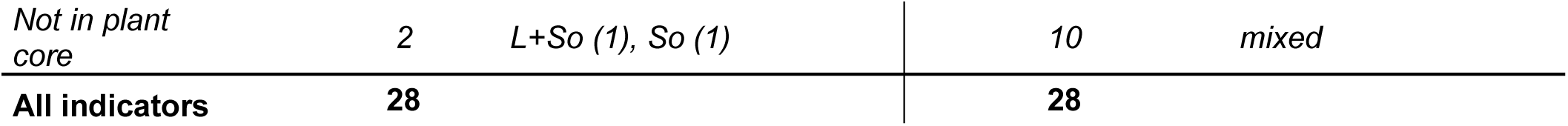
Cross-referencing of core microbiome membership and indicator species status at genus level. Plant core categories represent the intersection pattern of leaf, root, and seed 80% and 100% prevalence cores. N core = number of genera in that category; N indic. = number of those genera that are also significant indicators; Dominant pattern = most common indicator association pattern(s) with count in parentheses; N not in soil = number of genera not present in the soil core at the corresponding threshold.

#### 3.1.5. Differential abundance

Differential abundance analysis using ANCOM-BC2 identified 18 of tested genera (3.6%) as significantly differentially abundant in the omnibus test across sample types (BH-adjusted p < 0.05; **Table S10a-b**), of which 8 remained significant in at least one BH-corrected pairwise comparison (**Table S10**). Pairwise contrasts were strongly asymmetric: soil-versus-leaf and soil-versus-seed identified six differentially abundant genera, and soil-versus-root identified three, whereas among the plant tissue only the root-versus-leaf comparison yielded a single significant genus. No significant differences were detected between seeds and either leaves or roots.

Aggregated by parent class (**Table 4**, **Figure S7**), soil-versus-tissue contrasts were dominated by ***Gammaproteobacteria***. *Acinetobacter, Stenotrophomonas, Pseudomonas (E)*, and *Pseudomonas (B)* were enriched in plant tissues relative to soil in at least two comparisons, with ***Stenotrophomonas*** and ***Pseudomonas (E)*** differing significantly in all three. Additional tissue-specific enrichments included *Neorhizobium* (*Alphaproteobacteria*) and *Paenibacillus (J)* (*Bacilli*) in leaves; and *Neorhizobium, Agrobacterium, and Curtobacterium (Acinomycetes)* in seeds. Among plant tissues, only *Stenotrophomonas* differentiated root from leaf (higher in roots); whereas no genera differed significantly between seeds and either leaves or roots after BH correction.

**Table 4.**
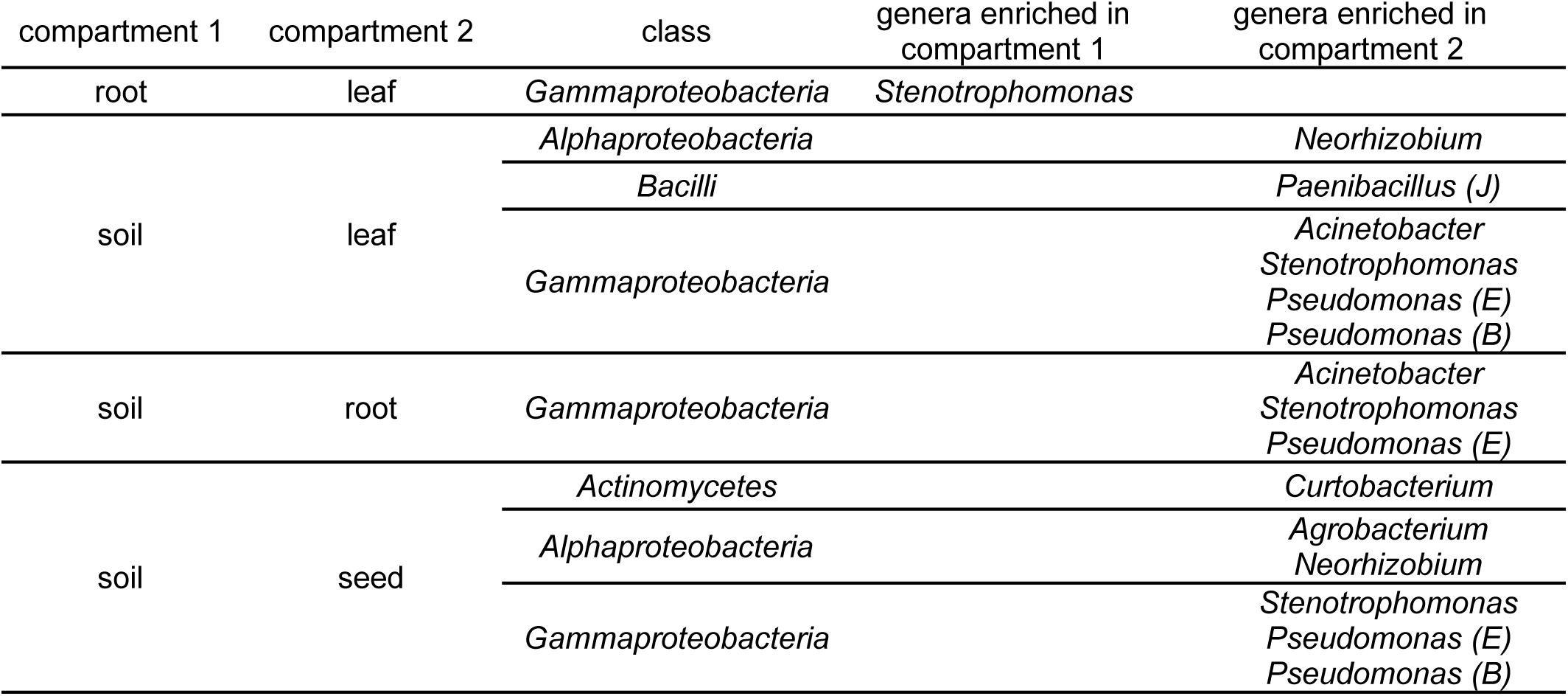
Differentially abundant genera between sample types, aggregated by taxonomic class (primary dataset). Significant pairwise contrasts from ANCOM-BC2 (BH-adjusted q < 0.05) at the genus level were grouped by their parent class. Detailed statistics in Table S10.s.

### 3.2. Functional prediction

#### 3.2.1. Beta diversity

Predicted functional structure was assessed using Bray-Curtis dissimilarity on relative abundances of the PGPT Lv5 profile. PCoA ordinations were computed on the K1-included dataset (**Figure 6**); statistical tests were performed on the K1-excluded dataset. In contrast to the taxonomic Bray-Curtis ordination (**Figure 4a**), the first two axes captured a substantially larger fraction of total variance (PC1: 45.2%, PC2: 19.7%). Samples separated primarily along PC1: all soil samples clustered in the negative-PC1 region, while seeds occupied the most positive-PC1 positions; leaf and root samples were intermixed in the central region. N4 characterized by the near-monodominance of *Pantoea*, was the most extreme positive-PC1 outlier among seeds. N3 deviated from the other seed samples, clustering closer to the leaf–root region. K1 was separated from all other samples along PC2, corresponding to its taxonomic anomaly extending to predicted function. Leaf and root samples showed no clear separation from each other in the ordination space.

**Figure 6.**
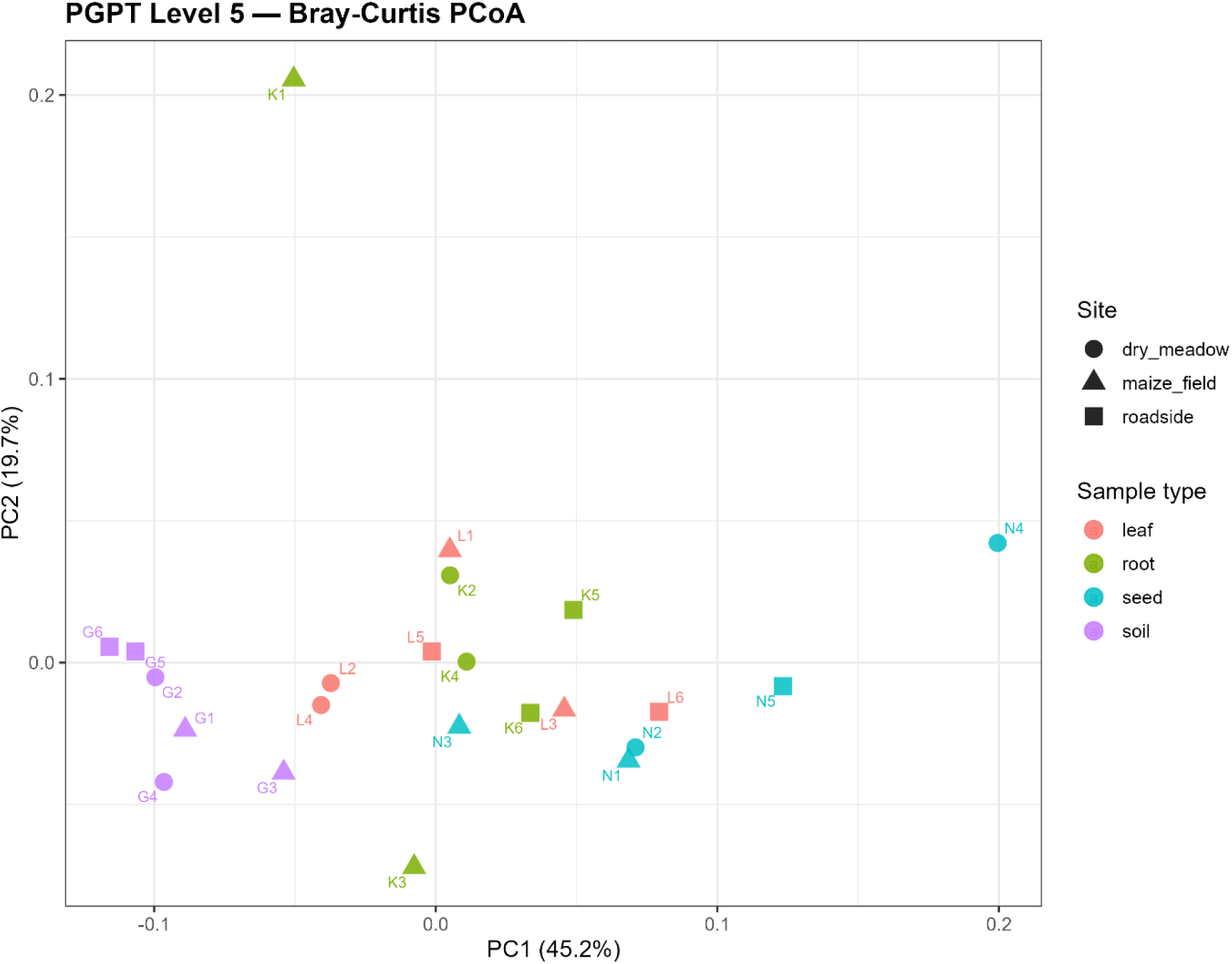
Principal coordinates analysis (PCoA) ordinations of predicted functional community composition – a PGPT Level 5 trait profile. The ordination is based on Bray-Curtis dissimilarity on relative abundances. Points are coloured by sample type and shaped by site. Sample labels are shown for identification.

PERMANOVA confirmed significant compositional differences among sample types (p = 0.001, R^2^ = 0.425; **Table S11a**), with a notably larger effect size than observed for taxonomic Bray-Curtis. No significant effects of region or site were detected in either the full or soil-excluded datasets (all p > 0.29; **Table S11a**). PERMDISP showed no evidence of dispersion heterogeneity for any factor combination (all p > 0.58; **Table S11b**), supporting the interpretation that the PERMANOVA result reflects centroid separation. Pairwise PERMANOVA with Benjamini-Hochberg correction (**Table S11c**) revealed a clear pattern: all contrasts involving soil were significant (leaf vs soil p = 0.004, R^2^ = 0.485; root vs soil p = 0.004, R^2^ = 0.353; seed vs soil p = 0.004, R^2^ = 0.533), as were seed contrasts with vegetative tissues (seed vs leaf p = 0.035; seed vs root p = 0.021). However, the **leaf-versus-root contrast was not significant** (p = 0.199, R^2^ = 0.121).

#### 3.2.2. Functional Core Microbiome and Indicator Feature Analysis

Functional core analysis at PGPT Level 5 identified large, stable cores across all compartments (**Figure S8a**). Core sizes ranged from 545–648 features at 50% prevalence, 485–597 at 80%, and 364–571 at 100%, with leaf retaining the largest core at all thresholds and seed showing the steepest decline at the strictest thresholds. The pan-tissue core included 578, 519, and 364 features at 50%, 80%, and 100% prevalence, respectively. Partitioning by soil core membership (**Figure S8b**) showed that the majority of pan-tissue features were shared with the soil core at all thresholds (513/578 at 50%, 450/519 at 80%, 343/364 at 100%), wheras the plant-specific subset consisted of 65, 69, and 21 features at 50%, 80%, and 100% prevalence, respectively. Venn analysis confirmed that the pan-all (all four compartments) intersection dominated the functional core (72% of all core features at 80% prevalence; 58% at 100%), while the pan-tissue plant-specific fraction accounted for 11% and 4%, respectively (Figure S8c–d). All remaining compartment combinations represented less than 13% of the functional (80% or 100%) core each. The complete abundance table of KEGG orthologs and their PGPT hierarchy is provided in **Table S12**.

Indicator species analysis at PGPT Level 5 identified 113 significant predicted indicators out of 964 features tested (BH-adjusted p < 0.05; **Table S13**), across eight compartment association patterns (**Table 5; Table S14–S15**). The dominant pattern was the **leaf+root+seed (plant-specific pan-tissue)** group, comprising 55 Lv5 features (32 Lv4 categories). These functions were primarily associated with plant-derived substrate utilization (predominantly carbohydrate transport), quorum sensing and biofilm formation, bacterial secretion systems (Sec, Type II and Type VI), cell envelope remodelling, stress tolerance, competitive fitness (toxin-antitoxin systems and antimicrobial resistance), insecticidal toxin production, phosphate solubilization, iron acquisition (siderophores and hemophores), heavy-metal resistance, and xenobiotic degradation.

**Table 5.** Functional indicator features identified by IndVal.g analysis at PGPT Level 5, aggregated to Level 4 categories. The number of contributing Level 5 indicator features is shown in parentheses after each Level 4 category. BH-adjusted p < 0.05, 999 permutations.

| Assigned group | Indicator features (Lv4, with n Lv5 in parentheses) |
| --- | --- |
| leaf+root+seed | plant-derived carbohydrate transport (13), QS autoinducer perception (4), toxin-antitoxin system (4), EPS amylovoran metabolism (2), insecticidal compounds (2), iron acquisition-siderophores (2), plant-derived aromatic/phenolic compound utilization (2), plant-derived sugar acid utilization (2), bactericidal compounds (1), mobile elements (1), multidrug resistance (1), resistance to antimicrobial/toxic compounds (1), glycerophospholipid remodelling (1), surface glycosylation patterns (1), peptidoglycan remodelling (1), Sec-secretion pathway (1), Type II secretion systems (1), Type VI secretion systems (1), host invasion factors (1), cobalt resistance (1), nickel resistance (1), tungstate resistance (1), high temperature tolerance (1), iron acquisition-hemophores (1), motility-pilus/fimbriae system (1), P-solubilisation-organic acid metabolism (1), plant-debranching spermidine/putrescine metabolism (1), plant-derived amino/nucleotide sugar utilization (1), plant-derived hexoses utilization (1), plant-derived organic acid transport (1), plant membrane polysaccharide degrading functions (1), xenobiotic benzoate/derivate degradation (1). **Subtotal: 32 Lv4 categories (55 Lv5 features)** |
| leaf+root | toxin-antitoxin system (3), multidrug resistance (2), plant-derived carbohydrate transport (2), adaptation to PIS gallate/tannine degradation (1), glycerophospholipid remodelling (1), peptidoglycan remodelling (1), iron acquisition-siderophores (1), low temperature tolerance (1), plant-signalling indole volatile metabolism (1), SAR-salicylic acid metabolism (1), LPS/lipid/IVA metabolism (1), xenobiotic benzoate/derivate degradation (1). **Subtotal: 12 Lv4 categories (16 Lv5 features)** |
| leaf+root+soil | xenobiotic benzoate/derivate degradation (4), bactericidal compounds (2), plant-derived amino/nucleotide sugar utilization (2), LPS/lipid/IVA metabolism (2), adaptation to PIS carotenoid derivate resistance (1), CRISPR-Cas system (1), glycerophospholipid remodelling (1), peptidoglycan remodelling (1), QS autoinducer perception (1), CO<sub>2</sub> fixation-RuBisCo (1), iron acquisition-other transport (1), N-acquisition-ethanolamine usage (1), neutralizing acidic stress (1), plant-derived amino acid utilization (1), plant-derived carbohydrate transport (1), plant-derived flavonoid utilization (1), plant-derived terpene utilization (1), lipid metabolism/oxidoreductase (1), xenobiotic toluene/derivate degradation (1). **Subtotal: 19 Lv4 categories (25 Lv5 features)** |
| root | bactericidal compounds (3), plant-derived hexoses utilization (2), plant-derived carbohydrate transport (2), QS autoinducer perception (1), xenobiotic styrene/derivate degradation (1), xenobiotic toluene/derivate degradation (1). **Subtotal: 6 Lv4 categories (10 Lv5 features)** |
| root+soil | bactericidal compounds (1), arsenic resistance (1), iron acquisition-siderophores (1). **Subtotal: 3 Lv4 categories (3 Lv5 features)** |
| leaf+seed | Toxin-antitoxin system (1). **Subtotal: 1 Lv4 category (1 Lv5 feature)** |
| leaf | xenobiotic benzoate/derivate degradation (1). **Subtotal: 1 Lv4 category (1 Lv5 feature)** |
| soil | xenobiotic dioxin/derivate degradation (1), xenobiotic hydrocarbons/oil degradation (1). **Subtotal: 2 Lv4 categories (2 Lv5 features)** |

The leaf+root+soil prediction pattern comprised 25 Lv5 features (19 Lv4 categories) dominated by xenobiotic degradation, bactericidal compound metabolism, carbohydrate utilization, adaptation to the plant immune system, CRISPR-Cas functions, CO_2_ fixation, nitrogen acquisition, and cell-envelope remodelling. The leaf+root pattern included 16 Lv5 features (12 Lv4 categories), notably including the only indicator related to plant immune response stimulation (salicylic acid transport), together with toxin-antitoxin systems, multidrug resistance, siderophore biosynthesis, carbohydrate transport, cell-envelope remodelling, indole volatile signalling, and stress adaptation. Root-exclusive indicators contained 10 Lv5 features (6 Lv4 categories), dominated by antibiotic biosynthesis, carbohydrate utilization and transport, quorum sensing, and aromatic compound degradation. The remaining patterns contributed relatively few indicators: root+soil comprised tylosin metabolism, arsenic resistance, and myxochelin biosynthesis; leaf+seed contained a single toxin-antitoxin system; leaf-only a single styrene/phenylethanol homeostasis function; and soil-only two xenobiotic degradation pathways. **No seed-exclusive functional indicators** were identified.

#### 3.2.3. Differential abundance

Differential abundance analysis using ANCOM-BC2 at PGPT Level 5 identified 613 of 964 features tested (63.6%) as significantly differentially abundant in the omnibus test (BH-adjusted p < 0.05; **Table S16a-b**). Of these, 362 unique features were confirmed in at least one pairwise comparison, yielding 752 significant pairwise contrast–feature combinations (**Table S16a,c**).

Aggregation to PGPT Lv2 categories (**Figure 7; Table S17**) confirmed that all eight Lv2 categories were engaged in at least one pairwise contrast. Soil-tissue contrasts engaged six to eight Lv2 categories, and tissue-tissue contrasts engaged six to seven. These comparison types were asymmetrical. Soil-versus-leaf was entirely leaf-enriched across all eight differentiated categories, including Plant Immune Response Stimulation, which was unique to this comparison. Similarly, all six significant categories in the soil-versus-root comparison were root-enriched, whereas the soil-versus-seed comparison showed six seed-enriched categories and Putative Functions as the only soil-dominant category.

**Figure 7.**
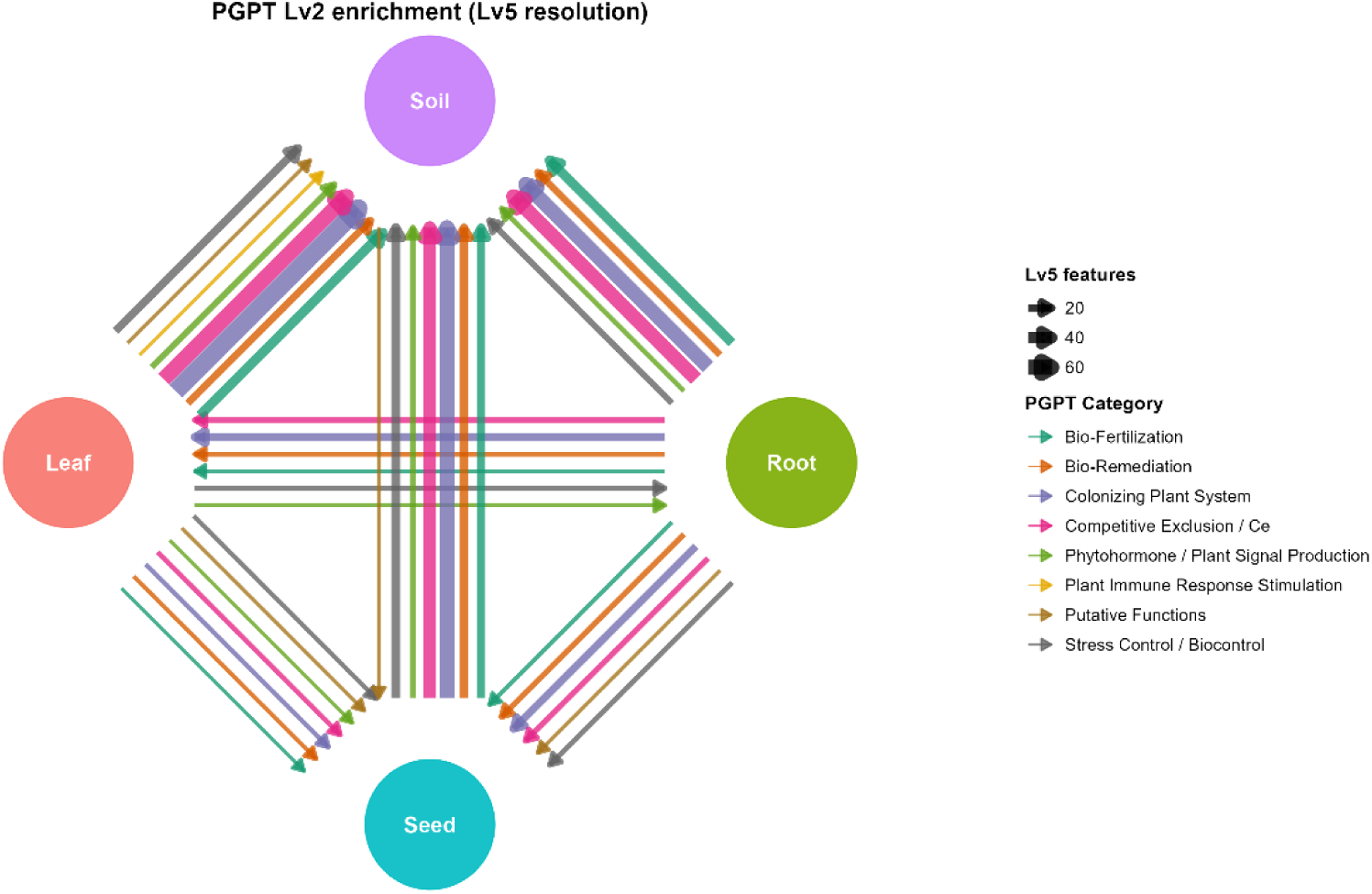
Network visualization of pairwise differential abundance in PGPT Lv2 categories between sample types. Nodes represent the four sample types (soil, leaf, root, seed). Edges represent significant differential abundance between compartment pairs (ANCOM-BC2, BH-adjusted p < 0.05) aggregated to PGPT Lv2 categories. Edge direction (arrowhead) points from the compartment with higher mean abundance ("dominant") to the compartment with lower abundance for each Lv2 category. Edge thickness is proportional to the number of constituent Lv4 features differentiating each pair within each Lv2 category. Edge color encodes the PGPT Lv2 category. PGPT Lv2 categories with tied directions in a given pair (equal numbers of Lv4 features higher in each compartment) are not represented as edges; full pairwise differential abundance results, including tied categories and underlying Lv4 features, are reported in Tables S16–S17.

At PGPT Level 5, soil-tissue contrasts yielded high numbers of differentially abundant features: soil-versus-leaf (230 features; 6:224), soil-versus-seed (191; 54:137), and soil-versus-root (187; 22:165). Tissue-tissue contrasts were substantially smaller in scale, including 59 features for seed-versus-root (14:45); 58 for root-versus-leaf (36:22), and only 27 for seed-versus-leaf, all of which were leaf-enriched.

Among tissue-tissue contrasts, seed-versus-leaf showed the strongest asymmetry with all seven Lv2 categories enriched in leaves. Seed-versus-root likewise showed root dominance in six of seven categories (one tied), whereas the root-versus-leaf comparison displayed a mixed pattern: root dominated in Bio-Fertilization, Bio-Remediation, Colonizing Plant System, and only slightly in Competitive Exclusion, while leaf dominated in Phytohormone/Plant Signal Production and Stress Control/Biocontrol. Predicted functional potential pointed to an order increasing from **soil < seed < leaf ≈ root**.

### 3.3. Integration of taxonomic and functional community structure

Concordance between taxonomic and functional community structure was assessed using Mantel tests and Procrustes superimposition on Bray-Curtis distance matrices computed on the K1-excluded primary dataset (n = 22). The Mantel test revealed a **strong and significant correlation** between taxonomic and PGPT Level 5 functional distance matrices (r = 0.770, p = 0.001; 999 permutations). Procrustes superimposition of the corresponding PCoA ordinations confirmed this concordance (Procrustes r = 0.620, p = 0.001; **Figure S9**).

## 4. Discussion

### 4.1. Tissue compartment as the primary determinant of the endobacteriome structure

Across the analyses, tissue compartment emerged as the sole factor differentiating *A. artemisiifolia*, endobacteriome, while neither geographic region nor habitat type showed such effect. This pattern was supported by alpha and beta diversity, indicator species analysis, core microbiome analysis, and predicted functional profiling. Although a significant regional effect was detected by Weighted UniFrac (beta diversity), it disappeared after excluding soil samples, was not supported by pairwise comparisons, and was not confirmed by any other analytical approach. Likewise, no evidence for habitat-specific structuring was detected.

Tissue compartment thus represents the dominant ecological filter shaping bacterial endophyte assembly in *A. artemisiifolia* across the sampled range, although leaf and root endophytes were not functionally distinguishable from each other. For that reason, subsequent analyses focused on compartment-specific core microbiomes, indicator taxa, and differentially abundant taxa and functions. For comparison, in the invasive *Asteraceae Senecio vulgaris* L., both tissue compartment and sampling location influenced endophytic community composition, although a stable bacterial core persisted across geographically separated populations (Cheng et al. 2019). Our results indicate that compartment identity outweighs geographic or habitat effects. Although our sampling design did not allow formal testing of region or habitat within each compartment separately, there is an absence of regular spatial patterns across multiple analyses, including indicator species analysis. Nevertheless, the *A. artemisiifolia* endobacteriome is characterized by a conserved, pan-tissue core shared across sites spanning two countries.

### 4.2. The shared plant-tissue endobacteriome

#### 4.2.1. Taxonomic pattern

Despite compartment-level differentiation, the three plant tissues shared a conserved taxonomic and functional profile that collectively distinguished them from soil. The most prominent pattern was a class-level pattern: ***Gammaproteobacteria*** dominated all endophytic compartments, whereas ***Alphaproteobacteria*** predominated in soil. This agrees with previous studies showing *Gammaproteobacteria* to dominate plant endospheres, while *Alphaproteobacteria* are more characteristic of rhizosphere communities (Gottel et al. 2011; Hardoim et al. 2015). Likewise, the dominance of *Pseudomonadota, Actinomycetota*, and *Bacillota* in our dataset matches the phyla described for the vast majority of bacterial endophytes (Hardoim et al. 2015).

In the differential abundance analysis, ***Pseudomonas (E)*** and ***Stenotrophomonas*** were enriched in plant tissues relative to soil across all soil-tissue comparisons, whereas ***Acinetobacter****, **Pseudomonas (B)**,* and ***Neorhizobium*** were enriched in at least two compartments. These findings indicate that the observed class-level inversion reflects changes in relative dominance, not the exclusion of *Alphaproteobacteria* from the endosphere. The enrichment of *Acinetobacter* should, however, be interpreted cautiously (see **Methods S5**) because the genus includes both recognised endophytes and common laboratory contaminants (Salter et al. 2014; Eisenhofer et al. 2019), a concern partially supported by its minimal presence in soil samples in this dataset. Quantitative microbiome profiling further demonstrated that, despite their low relative abundance in soil, *Gammaproteobacteria* reached absolute abundances comparable to or exceeding those in leaves, though large within-compartment variance precluded statistical confirmation of these differences.

The **pan-tissue core** at 80% prevalence comprised 13 genera, divided into two ecologically distinct groups. Four – ***Pseudomonas (E), Sphingomonas, Methylobacterium***, and ***Devosia*** – were also soil core members, and represented ubiquitous environmental bacteria recruited into and sustained across all compartments. The remaining nine genera (***Pantoea, Paenibacillus (J), Telluria, Neorhizobium, Pseudomonas (B), Acinetobacter, Curtobacterium, Ralstonia***, and ***Kineococcus***) constituted a plant-specific core absent from the soil core. At the strictest prevalence threshold (100%), five of these genera (***Pantoea, Paenibacillus (J), Telluria, Neorhizobium, Pseudomonas (B***)) remained plant-specific (additionally, **Methylobacterium)**, indicating that the most persistent endophytes are selectively maintained within plant tissues, and may partly be inherited via seeds (see <u>Wu et al. (2026)</u>).

Several members of the plant-specific core are well-established endophytes or plant growth-promoting bacteria (Hardoim et al. 2015; Rani et al. 2022). Particularly noteworthy is ***Pantoea***, whose members span a continuum from phytopathogens to plant growth-promoting and bio-control bacteria, with ecological outcomes depending on strain identity, host physiology, and microbial community context (Doni et al. 2021). A similar continuum characterizes ***Pseudomonas***, which includes numerous biocontrol strains alongside important plant and opportunistic human pathogens (De Bentzmann and Plésiat 2011; Brader et al. 2017). ***Paenibacillus*** is another well-established endophytic genus possessing multiple plant growth-promoting traits, including nitrogen fixation, phosphate solubilization, siderophore production, and phytohormone synthesis (Rani et al. 2022). The occurrence of ***Ralstonia*** within the pan-tissue core is noteworthy because the genus is primarily known for destructive plant pathogens and opportunistic human pathogens (Genin and Denny 2012; Ryan and Adley 2014), with no beneficial plant-associated role reported for this genus. Its ecological significance therefore remains uncertain and will require species-level identification beyond the taxonomic resolution of the V3-V4 region (Johnson et al. 2019). ***Curtobacterium, Telluria*** (reclassified from *Massilia*)**, *Kineococcus, Neorhizobium*** (delineated from *Rhizobium*)**, *Sphingomonas, Methylobacterium*** and ***Devosia*** have likewise been reported as endophytic bacteria, which supports the biological relevance of the identified core microbiome.

In the only so-far published high-throughput sequencing analysis of *A. artemisiifolia*’s endobacteriome, Wu et al. <u>(2026)</u> identified taxa shared by seeds and seedlings. *Pseudomonas* and *Sphingomonas* were among them, as in our study. Moreover, *Pseudomonas* was vertically transmitted in all five Chinese geographic populations. However, the second universally vertically transmitted genus, *Bacillus*, is almost absent in our samples, suggesting that even the taxa vertically transmitted in one region across all populations studied do not necessarily belong to the plant core in other invasion ranges.

Overall, the composition of the pan-tissue core supports the concept that the endosphere is not inhabited exclusively by mutualists, but by microorganisms spanning a continuum from beneficial symbionts to latent pathogens, with ecological outcomes determined by microbial genotype, host physiology, population density, quorum sensing, and environmental context (Hardoim et al. 2015; Grzyb and Szulc 2024). The leaf+root+seed indicator pattern contained five genera (***Pseudomonas (B), Paenibacillus (J), Pantoea, Neorhizobium****, and **Kineococcus***), four of which belonged to the **plant-specific 100% core**. Across differential abundance results, 15 of 16 significant comparisons contrasted soil against one or more plant tissues, with only a single tissue-tissue contrast detected (Stenotrophomonas, root > leaf). The pan-tissue core, the leaf+root+seed **indicator pattern**, and **differential abundance results are concordant**, pointing to a conserved bacterial consortium.

The existence of a shared pan-tissue core is consistent with the root-mediated colonization model in which bacteria establish endophytic populations in roots before spreading systemically to aerial tissues (Lengrand et al. 2024). Consequently, overlap among endophytic compartments exceeds that between endophytes and soil communities despite soil representing the principal source of colonizing bacteria.

#### 4.2.2. Functional perspective

The dominant leaf+root+seed indicator pattern, comprising 55 PGPT Level 5 and 32 Level 4 categories, defined the functional pattern distinguishing the endobacteriome from soil. This pattern is consistent with the prevailing model of endophytic colonization, in which bacteria are recruited primarily through roots, attach to the root surface, enter internal tissues, and subsequently spread systemically to aerial organs and localize within particular tissues (Compant et al. 2021; Rani et al. 2022; Lengrand et al. 2024). Root exudates provide diverse substrates that shape microbial recruitment, whereas organ-specific chemical environments and alternative transmission routes, including stomatal, floral, aerial, and insect-mediated entry, may contribute to compartment-specific differentiation (Frank et al. 2017; Compant et al. 2021). Thus, root-mediated systemic colonization may explain the shared pan-tissue functional set, while alternative entry routes and tissue-specific conditions may generate the differences discussed below.

**Colonizing Plant System** category functions supported this interpretation. **Plant-derived substrate utilization** included carbohydrate transport and the metabolism of aromatic compounds, sugar acids, hexoses, and organic acids, in line with the composition of root exudates and other tissue-derived substrates. Pectate lyase represented **plant cell-wall degradation**, whereas the type IVa pilus represented **surface attachment** and **motility**. Experimental studies have shown that disruption of cell wall-degrading enzymes or chemotaxis can reduce endophytic colonization efficiency (Rani et al. 2022), supporting the biological relevance of these predictions.

Indicator and differential abundance analyses provided complementary views of functional differentiation. IndVal.g identifies features that are both specific to and consistently present within a compartment (De Cáceres and Legendre 2009), whereas differential abundance detects quantitative shifts regardless of prevalence. Several colonization-related genes (e.g. *fliC, mglB, pilX)* had much higher predicted abundance in plant tissues relative to soil (**Table S12**) but did not qualify as Level 5 indicators because each Level 5 category aggregates multiple genes, potentially diluting gene-specific signals. The much larger number of significant features detected by differential abundance analysis than by indicator analysis reflects this difference in analytical logic. Aggregation to higher PGPT levels was therefore necessary for biological interpretation, particularly because PICRUSt2-based assignments indicate predicted genomic potential, not the confirmed pathway activity. The identification of amylovoran-type EPS biosynthesis despite the absence of *Erwinia* in the taxonomic dataset illustrates this limitation: predicted functions may derive from homologous gene clusters in related taxa.

Functions related to microbial competition and persistence formed another part of the plant-associated profile. **Quorum sensing** and **biofilm formation** were represented by autoinducer perception, consistent with their established roles in density-dependent regulation and host colonization (Hartmann et al. 2024). **Toxin-antitoxin systems** are diverse gene modules involved in plasmid stability, phage protection, stress tolerance, and biofilm formation; despite their ubiquity in plant-associated bacterial genomes, close to nothing is known regarding their relevance in the phytobiome (Shidore & Triplett, 2017).

**Bacterial secretion systems** were also detected, with the Sec pathway and Type II and Type VI systems identified as pan-tissue indicators. Secretion profiles have been perceived as markers of endophytic (T1SS, T2SS) or pathogenic (T3SS, T4SS) lifestyles (Khare et al. 2018; Kumar and Nautiyal 2023); however, exceptions exist in both directions: traditionally endophyte-associated systems can be essential for pathogenicity, while traditionally pathogen-associated systems can promote beneficial colonization (Genin and Denny 2012; Khare et al. 2018; Hu et al. 2021). The biological effect depends largely on the secreted effectors rather than the apparatus itself (Brader et al. 2017). In our differential abundance results, T1SS, T2SS, T3SS, T5SS, and T6SS were enriched in leaves and roots relative to soil (**Table S16**), whereas only T6SS was enriched in seeds. Together with the membrane-damaging proteins identified as pan-tissue indicators, these patterns reflect a fluid boundary between mutualistic and antagonistic behaviour in the endosphere.

Predicted **Stress Control/Biocontrol** functions included the FIT system, and the Tc toxin component TccC. These functions are associated with insecticidal activity and may contribute to protection against herbivores (Chen et al. 2014; Arrebola et al. 2022). **Bio-Fertilization** indicators included phosphate solubilization, hemophore-mediated iron acquisition, and siderophore biosynthesis. **Endophyte-mediated phosphate solubilization** may offer advantages over rhizobacterial mechanisms: Varga et al. (2020) demonstrated that phosphate taken up by poplar can form insoluble compounds within plant tissues, and endophytes being within the tissue are able to re-release phosphate in situ. Siderophores function both in plant iron nutrition (although the precise molecular mechanism by which bacterial siderophores deliver iron to the plant host remains experimentally unconfirmed) and in restricting pathogen access to iron (Rani et al. 2022; Grzyb and Szulc 2024). Finally, Bio-Remediation indicators included resistance to cobalt, nickel, and tungstate and the degradation of benzoate-related xenobiotics, in line with the broader role of endophytes in bioremediation and with the acquisition of such traits through horizontal gene transfer (Compant et al. 2021; Rani et al. 2022).

The discussed functions suggest mechanisms through which the endophytic microbiome may contribute to plant performance, although they represent predicted genomic potential and require experimental validation.

### 4.3. Seed endobacteriome

The seed-associated endophytic microbiome remains **one of the least characterized** compartments of the plant holobiont, with available studies heavily biased towards crop species. To our knowledge, the *A. artemisiifolia* seed endobacteriome has been characterized one study of five Chinese populations. Seeds are unique among plant compartments because they integrate both horizontal and vertical microbial transmission. While horizontal acquisition occurs from the surrounding environment, vertically transmitted microorganisms connect the microbiomes of successive plant generations (Abdelfattah et al. 2023; Qi et al. 2026). Among horizontal vectors, anthropochory is considered a leading cause of rapid spreading for *A. artemisiifolia* (Knolmajer et al., 2024) and may represent a significant yet entirely unexplored source of microbial contact. Because mature seeds were collected directly from plants, post-dispersal environmental colonization was excluded from this study. In annual invasive species such as *A. artemisiifolia*, the inherited microbiome may influence germination, seedling establishment, nutrient acquisition, stress tolerance, and pathogen resistance. The potential influence of the seed microbiome on invasion success remains largely unexplored and needs further attention.

Seed endophytic communities showed **the lowest alpha diversity** of all compartments across all diversity metrics. This agrees with previous studies showing that seeds represent a major bottleneck in microbiome transmission, harbouring up to 100-fold fewer taxa than vegetative tissues (Newcombe et al. 2018; Abdelfattah et al. 2023). Despite this reduced diversity, seed communities varied widely between samples. N4 was almost entirely dominated by *Pantoea*, whereas N3 showed the greatest taxonomic evenness and contained the highest number of genera exceeding 1% relative abundance. Similar variability has been reported for seed microbiomes of numerous plant species and may reflect variety described for stable core and transient microbial fractions (Acuña et al. 2023; Davies et al. 2024; Qi et al. 2026). Variability can occur even between individual seeds of the same plant, though Bintarti et al. (2022) found the highest microbiome variability at the plant level rather than the singular seed level, suggesting that pooling seeds from the same plant is appropriate for experimental designs.

*Pseudomonas (E)* and *Pantoea* occurred in all seed samples, while *Paenibacillus (J)* represented a frequent secondary taxon. Their occurrence within both the pan-tissue core and the pan-tissue indicator assemblage indicates that these genera are stable members of the broader *A. artemisiifolia* endobacteriome rather than seed-specific colonizers. The near-monodominance of *Pantoea* in N4 resembles the reports describing this genus as one of the most prevalent vertically transmitted seed endophytes across diverse host species (Simonin et al. 2022; Sanz-Puente et al. 2025). In the study by Wu et al. <u>(2026)</u>, *Pantoea*, was a seed core genus and ranged from 0.2% to 77% across ragweed’s Chinese populations (although it was not vertically transmitted). Their seed endophytic core included *Pseudomonas* (as in our analysis), but also *Pedobacter* (in 3 of our 5 seed samples), *Bacillus* (in 1 of 5), *Oceanobacillus,* and *Achromobacter* (in none of our samples). As seed communities differed even among the five Chinese populations (Wu et al. 2026), differences between European and Chinese populations are expected.

Differential abundance analysis showed that seeds differed significantly from soil but not from leaves or roots. Genera enriched in seeds relative to soil included ***Curtobacterium*, *Agrobacterium*, *Neorhizobium*, *Pseudomonas (E)*, *Pseudomonas (B)***, and ***Stenotrophomonas***. More broadly, the major seed-associated genera identified here – including *Pantoea*, *Pseudomonas*, *Sphingomonas*, *Paenibacillus*, *Methylobacterium*, *Curtobacterium*, *Ralstonia*, *Neorhizobium*, and *Devosia* – have repeatedly been reported from seed microbiomes of diverse plant species (Bziuk et al. 2021; Simonin et al. 2022; Zhang et al. 2022; Fadiji et al. 2025; Qi et al. 2026). Thus, the ragweed seed microbiome appears to consist largely of widespread plant-associated endophytes, and not necessarily taxa unique to this invasive host.

The seed core comprised only 14 genera at 80% prevalence, the smallest core among all compartments. Four genera were shared with soil, nine belonged to the plant-specific pan-tissue core, and only ***Sphingomonas (R)*** was exclusive to seeds. Conversely, the five genera forming the leaf+root+soil indicator (*Reyranella, Sphingomicrobium, Arthrobacter, Bradyrhizobium*, *Chryseolinea*) pattern were largely absent from seeds, although N3 retained several of them at considerable abundance. The predicted functional profile matched the taxonomic results. Functional cores were dominated by pan-all and plant-specific pan-tissue categories, whereas no seed-exclusive functional indicator was identified. The only additional seed-associated pattern was the leaf+seed combination, represented by a single toxin-antitoxin system.

Differential abundance analysis further showed that leaves and roots possessed significantly higher predicted functional abundances than seeds across all PGPT Level 2 categories that differed, whereas seeds remained enriched relative to soil in all differing categories except Putative Functions. Seeds therefore retained a characteristic **endophytic functional profile distinct from soil,** but quantitatively reduced relative to vegetative tissues, as in the functional gradient **soil < seed < leaf ≈ root**. The taxonomic and functional data, indicates that the *A. artemisiifolia* seed endobacteriome represents a filtered, **bottlenecked** subset of the broader plant-associated microbiome. Features the microbiome requires to colonize seeds, including plant cell-wall modification, motility, substrate utilization, and quorum sensing, correspond directly to the pantissue functional profile. Whether this profile primarily reflects passive filtering during seed maturation or active maternal selection remains an open question.

### 4.4. Root and leaf endobacteriomes – beyond seeds

Beyond the shared pan-tissue core and pan-tissue indicator profile discussed in section 4.2, roots and leaves exhibited distinct compartment-specific features. Root samples were collected from the middle section of the taproot at the onset of flowering and therefore represent established endophytic communities after host-mediated selection. These spatial and temporal constraints should be considered when interpreting root-specific patterns.

Alpha diversity did not differ significantly between roots and leaves, as all significant pairwise contrasts involved seeds. In contrast, Bray-Curtis beta diversity analysis showed significant taxonomic differentiation between the two vegetative compartments, whereas their predicted functional profiles did not differ significantly. Roots and leaves thus harboured taxonomically distinct but potentially functionally redundant bacterial communities.

#### 4.4.1. Taxonomy – roots

Root samples exhibited the highest between-sample heterogeneity of any compartment. Although *Gammaproteobacteria* generally dominated, **no two root samples shared the same hierarchy at lower taxonomic levels**. *Stenotrophomonas*, *Pseudomonas (E)*, *Streptomyces*, *Acinetobacter*, and *Pantoea* accounted for much of the community, but their relative positions varied markedly. ***Stenotrophomonas*** dominated three of five samples, whereas *Pseudomonas (E)*, repeatedly dominant or co-dominant in leaves and seeds, never ranked first in roots. This variability suggests a strong influence of sample-specific local conditions on root communities.

***Paraburkholderia*** was stably present in roots. The genus comprises primarily environmental and plant-associated species (no described pathogens so far), but as phylogenetic position alone does not predict pathogenic potential, strain-level characterization remains necessary (Eberl and Vandamme 2016). Its best-studied member, *P. phytofirmans* PsJN, is a model beneficial endophyte capable of colonizing diverse hosts, promoting growth, and enhancing tolerance to biotic and abiotic stress (Esmaeel et al. 2018).

At 80% prevalence, roots contained 24 exclusive core genera, the second-largest compartment-specific core after leaves. They shared 12 core genera exclusively with leaves, none exclusively with seeds, and only three with soil. This indicates that the root core was not just a passive reflection of the surrounding soil microbiota. Four genera, ***Ferrovibrio*, *Actinoplanes*, *Kribbella***, and ***Agromyces***, were identified as root-specific indicators, three of which belonged to *Actinomycetota*. Endophytic *Actinomycetes* are widely recognized for plant growth promotion, antifungal activity, and production of bioactive compounds (Shan et al. 2018). *Actinoplanes* includes strains with demonstrated biocontrol activity (El-Tarabily 2003), whereas *Kribbella* and *Agromyces* have been detected in healthy or stressed root endospheres (Liu et al. 2019; Li et al. 2023). *Ferrovibrio*, although rarely reported as an endophyte, has also been detected in rice roots and increased under abiotic stress (Chalouhi et al. 2025). The association of *Actinomycetes* with vegetative tissues was further supported by *Brachybacterium* as a leaf+root indicator and *Kineococcus* as a member of the plant-specific pan-tissue core. *Streptomyces*, despite dominating K3, was not a significant indicator because of its irregular occurrence. ***Rhizobium (E)*** was associated with the root+soil pattern, consistent with a niche at the root–soil interface. This GTDB lineage includes plant-associated taxa such as *R. setariae*, which produces indole-3-acetic acid (Kang and Seo 2022).

Differential abundance analysis identified ***Acinetobacter, Stenotrophomonas,*** and ***Pseudomonas (E)*** as enriched in roots relative to soil. The only tissue vs. tissue contrast in the entire dataset – ***Stenotrophomonas*** (root > leaf) – involved this compartment. **The number and taxonomic range of differentially abundant genera increased with distance from the root entry point**: only three genera, all *Gammaproteobacteria*, distinguished roots from soil, whereas leaves and seeds each differed from soil by six genera spanning additional classes. This pattern may reflect p**rogressive host-mediated selection** along the colonization route.

Two earlier high-throughput studies analyzed root-associated bacteria in China (Li et al. 2025; Wu et al. 2026). Li et al. <u>(2025)</u>, who did not distinguish epiphytes from endophytes, reported substantial variation among populations, whereas in Wu et al. <u>(2026)</u>, endophytes from roots of plants grown in a common garden were similar across populations. Our field-collected root endophytes were highly variable, which may suggest that differing local conditions, and uniform ones in the common garden, are an important factor.

Several dominant genera reported by Li et al. <u>(2025)</u> overlapped with our core or indicator taxa, including *Pseudomonas*, *Sphingomonas*, *Pantoea*, *Neorhizobium*, and *Actinoplanes*. Root endophytes studied by Wu et al. <u>(2026)</u> also contained *Pseudomonas*, rhizobia, and *Actinoplanes*. However, they were dominated by unclassified *Alcaligenaceae*, with no corresponding match in our data. Other dominant taxa reported by Li et al. <u>(2025)</u>, including *Candidatus Phytoplasma*, *Halomonas*, and *Flavobacterium*, as well as *Roseateles* reported by Wu et al. <u>(2026)</u>, were absent from our roots. These differences may reflect geography, host population, primer selection, as well as the inclusion of epiphytes (Li et al. 2025) or common-garden conditions (Wu et al. 2026). The occurrence of the same dominant genera across European and Chinese populations suggests a partially conserved root-associated bacterial core associated with this invasive species.

#### 4.4.2. Taxonomy – leaves

The leaf endobacteriome develops within the highly variable phyllosphere environment, characterized by fluctuations in moisture, nutrients, temperature, and UV exposure. Internal tissues offer greater protection than leaf surfaces but expose bacteria to closer host immune surveillance. Leaf-associated microorganisms may originate from soil, seeds, air, or rain, while *Sphingomonas*, *Methylobacterium*, and *Pseudomonas* are commonly dominant in eudicot phyllosphere communities (Grzyb and Szulc 2024).

Leaf communities were more compositionally stable than root or seed communities. Four of six samples followed the general endophytic pattern of *Gammaproteobacteria* dominance. In L2 and L4, however, *Alphaproteobacteria* exceeded *Gammaproteobacteria* – the only instances across plant compartments – accompanied by increased contributions of *Sphingomonadales* and greater genus-level evenness. Both samples originated from meadows in different regions, although the present design does not allow this pattern to be attributed conclusively to habitat. At the genus level, ***Pseudomonas (E)*** was abundant in all leaf samples, whereas *Stenotrophomonas* remained scarce. Other prominent genera largely overlapped with the pan-tissue core, including ***Sphingomonas*, *Acinetobacter*, *Pantoea*, *Paenibacillus* (J)**, and ***Methylobacterium***. ***Sphingomicrobium***, an indicator of the leaf+root+soil pattern, was also repeatedly present. ***Rossellomorea*** (delineated from *Bacillus*) occurred as a minor component in five of six leaf samples. Although little is known about its leaf-endophytic occurrence, representatives have been isolated from several plant endospheres and shown to possess ACC deaminase activity, growth-promoting properties, or heavy-metal tolerance (Navarro-Torre et al. 2021; Khamwan et al. 2022). *Stutzerimonas* exceeded 1% abundance only in L2 but has previously been reported among leaf endophytes with antibacterial properties (Priyanto et al. 2024).

Leaves contained the largest exclusive core at 80% prevalence, but no genus was a significant leaf-only indicator. The strongest leaf-associated signal instead involved shared patterns, particularly leaf+soil, which comprised ten genera and represented the largest indicator group in the dataset. These mostly **uncultured or poorly characterized taxa** formed a stable but low-abundance part of leaf communities (no leaf+soil functional indicator was detected) while remaining absent from roots and seeds. The leaf+soil pattern suggests recruitment routes that may bypass stable root colonization. Bacteria may reach leaves through rain splash, soil-derived bioaerosols, dust, or direct entry through stomata, and wounds (Frank et al. 2017; Lengrand et al. 2024). Such pathways have been demonstrated for both pathogenic and beneficial endophytes, while host regulation of stomatal opening may contribute to microbial selection during entry. The dominance of uncultured lineages within the leaf+soil indicator group limits interpretation but illustrates an unexplored diversity within the *A. artemisiifolia* leaf endobacteriome.

Differential abundance analysis identified ***Neorhizobium, Paenibacillus (J), Acinetobacter, Stenotrophomonas, Pseudomonas (E)***, and ***Pseudomonas (B)*** as enriched in leaves relative to soil. The broader taxonomic range of leaf-enriched genera than of root-enriched genera is fitting to a progressive community differentiation from soil during colonization.

Two shotgun-metagenomics studies of *A. artemisiifolia*, did not distinguish epiphytes from endophytes. Bieker et al. (2020), detected species of *Pseudomonas*, *Pantoea*, and *Methylobacterium* in modern or herbarium specimens – all genera present in our pan-tissue core. A subsequent pathogen-focused survey (Bieker et al. 2022) reported phytopathogens belonging to *Agrobacterium, Pseudomonas, Pantoea, Curtobacterium, Ralstonia, Clavibacter* and *Pectobacterium*, all also detected in our leaf samples. Two genera reported in 2022 were absent from our leaf endobacteriome but present in other compartments: *Xanthomonas* occurred only in seeds, and *Herbaspirillum* only in roots and one seed sample. Other phytopathogens reported by Bieker et al. were not detected in any compartment. These discrepancies likely reflect the distinction between total leaf-associated microbiota and the endophytic fraction, differences in geography and host population, and preservation-related biases in herbarium specimens. The Bieker et al. (2022) pathogen list also included recognized plant growth promoters and nitrogen-fixing symbionts. There is a need for species-level identification before assigning ecological roles. In the common garden experiment of Wu et al. <u>(2026)</u>, leaf endophytes were dominated by *Sphingomonas*, *Methylobacterium* (both present in our pan-tissue core), and *Aureimonas*, which in our samples were present only in two root and two seed samples, but not leaves. *Pseudomonas, Curtobacterium*, and rhizobia (present in our core, rhizobia by *Neorhizobium*), were also a visible fraction. This also may illustrate the potential role of geography or conditions (field vs. cultivation).

#### 4.4.3. Roots and leaves – functional perspective and functional redundancy

The predicted functional profiles of roots and leaves did not differ significantly in beta diversity, despite significant taxonomic differentiation between these compartments. This pattern aligns with **partial functional redundancy**, whereby some phylogenetically distinct taxa perform the same focal functions (Louca et al. 2018). Functional redundancy is an inevitable property of open microbial communities, being more pronounced for broad than for narrow functional categories (Louca et al. 2018; Chen et al. 2022a). For comparison, using a different approach, Bai et al. (2015) reported extensive functional overlap between leaf- and root-derived bacteria in *Arabidopsis thaliana*, but alongside taxonomic overlap in the cultured bacteria, not with the taxonomic divergence observed here.

Nevertheless, differential abundance and indicator analyses revealed functional specialization. Only 58 PGPT Level 5 features differentiated roots from leaves, compared with 187 and 230 features distinguishing roots and leaves from soil, respectively. This indicates that the functional contrast between vegetative tissues was substantially weaker than their differentiation from soil. Root-enriched functions predominated in **Colonizing Plant System**, **Bio-Remediation**, and **Bio-Fertilization**, whereas leaf-enriched functions were associated mainly with **Phytohormone/Plant Signal Production** and **Stress Control/Biocontrol; Competitive Exclusion** was almost balanced. Indicator analysis complemented these abundance-based differences by identifying functions consistently associated with particular compartment combinations.

Beyond the shared pan-tissue functional pattern, vegetative tissues showed an expanded set related to plant-associated lifestyles. Additional indicators included utilization of plant-derived substrates (carbohydrates, amino acids, flavonoids, and terpenes), degradation or transport of plant defence compounds, and lipid/LPS metabolism associated with bacterial attachment. **Flavonoids** are particularly important because they act as microbial substrates, antimicrobial compounds, and signalling molecules involved in microbial recruitment and rhizobial *nod*-gene activation (Compant et al. 2021; Wang et al. 2022). Carbohydrate transport and xenobiotic degradation indicators showed progressive expansion from shared multi-compartment to root-exclusive patterns.

Competitive and persistence-related functions were likewise expanded, including CRISPR–Cas systems, quorum sensing, toxin-antitoxin systems, multidrug efflux pumps, and indole metabolism. Root-associated indicator patterns further included biosynthesis of antimicrobial compounds, additional siderophore pathways, xenobiotic degradation, and stress-response mechanisms, matching the more heterogeneous and chemically complex below-ground environment. **Salicylic acid** transport was the only indicator assigned to Plant Immune Response Stimulation, supporting the key role of salicylic acid signalling in both plant defence and beneficial plant– endophyte interactions (Benjamin et al. 2022).

The order **soil < seed < leaf ≈ root** remained relatively stable. Compared with soil, both vegetative compartments showed enrichment across nearly all functional categories, whereas seeds displayed lower predicted functional abundance than leaves and roots. The pan-tissue set therefore prepares for colonization, persistence, competition, and nutrient acquisition, while vegetative tissues add functions related to specialized substrate utilization, antimicrobial activity, stress tolerance, and bioremediation.

**Roots occupied the central position in the functional network**, participating in nearly all multi-compartment indicator patterns and displaying the largest compartment-specific functional profile. Leaves, in contrast, showed almost no exclusive functional indicators. The strong leaf+soil association observed in the taxonomic analysis was largely absent from the functional indicator profile, suggesting that taxa shared between leaves and soil do not contribute unique functional capacities beyond those already common to the plant endobacteriome.

### 4.5. Total bacterial load

Quantitative microbiome profiling by qPCR revealed a non-significant gradient of total bacterial 16S rRNA gene copies across compartments: soil > roots (7.47 × 10^7^ – 8.89 × 10^8^) > leaves (1.31 × 10^7^ – 5.03 × 10^8^) > seeds (1.67 × 10^7^ – 3.94 × 10^8^). Although these differences were not statistically significant, the estimates address an important gap in endophyte ecology, as quantitative measurements of total endophytic bacterial load remain relatively uncommon. Culture-based counts have been reported across compartments, but **gene-copy-based estimates remain scarce**. The seed estimates obtained here fall within the range previously reported for vegetable seeds (Acuña et al. 2023).

These estimates should be interpreted with caution because the analyzed compartments differ fundamentally in tissue structure and cell density. Moreover, root samples represented the middle section of the taproot rather than the actively colonized root apex, universal 16S rRNA gene primers may co-amplify plant organellar DNA despite correction using sequencing data, and limited primer coverage for several soil-associated phyla probably resulted in underestimation of soil bacterial abundance. Despite these limitations, the results provide the first quantitative baseline for bacterial colonization levels in the *A. artemisiifolia* endobacteriome, facilitating future comparisons across populations, host species, and invasion contexts.

### 4.6. Concordance and methodological limitations

Functional profiles were inferred using PICRUSt2, which predicts gene content from phylogenetic placement of ASVs rather than directly measuring genes (Douglas et al. 2020). Although shotgun metagenomics provides direct functional information, its application to endosphere samples remains challenging because host DNA typically dominates sequencing output (even over 98% of total reads) (Verhoeven et al. 2025). Amplicon sequencing remains the practical approach for endophyte profiling, though co-amplification of host plastid and mitochondrial sequences requires mitigation. Given these constraints, PICRUSt2 represents the most accessible tool for estimating functional potential from amplicon data. The recently updated PIC-RUSt2-SC reference database, based on GTDB, further improves prediction accuracy and aligns with the taxonomic framework used in this study (Wright and Langille 2025).

Concordance between taxonomic and functional community structure was strong and statistically significant. Part of this correlation is expected, as PICRUSt2 infers gene content from the ASVs themselves: samples with similar ASV composition will also have similar predicted profiles. The opposite, however, does not need to be true, and the relationship was imperfect, in line with the potential partial functional redundancy observed between root and leaf endobacteriomes. What is interesting, phylogeny-weighted metrics (Weighted UniFrac and U^A^) did not separate compartments, while Bray-Curtis did.

Several methodological limitations should be considered. Predicted functions reflect genomic potential (not the confirmed activity), and taxonomic resolution was limited by V3–V4 16S rRNA gene sequencing (Johnson et al. 2019). Comparisons with other studies were further constrained by differences in 16S rRNA gene region, databases, and sequence units. For instance, Wu et al. <u>(2026)</u>, use 97% OTUs whereas we opt for ASVs, offering finer resolution and being reproducible across studies (Callahan et al. 2017). Primer bias likely underestimated some soil-associated taxa, while contaminant filtering (a partly subjective process) and exclusion of root sample K1 (**Methods S5**) inevitably involved analytical decisions. The sample size (5–6 samples per compartment) was sufficient for between-compartment comparisons but did not allow robust testing of within-compartment functional redundancy or geographic effects. Quantitative microbiome profiling was additionally constrained by high within-compartment variability, preventing reliable comparisons of absolute bacterial abundances.

Finally, the study represents a single developmental stage (onset of flowering for leaves and roots) and one root region (middle taproot), targeting already established endophytes. Although sampling across six sites in two countries demonstrated that compartment effects outweighed spatial variation, broader geographic and seasonal sampling will be needed to evaluate the generality of the conserved endobacterial core identified here.

### 4.7. Invasion biology context

The microbiome has been increasingly recognized as a missing element in our understanding of plant invasions (Gioria et al. 2023). Endophytes may be implicated in several hypotheses including endophyte-mediated enemy release, enhanced mutualism, disease-mediated invasion (spillover and spillback), plant-soil feedback, and invasional meltdown (Reinhart and Callaway 2006; Simberloff 2006; Strauss et al. 2012; Bard et al. 2024). Despite growing interest in bacterial endophytes possessing plant growth-promoting and biocontrol properties (Rani et al. 2022), their role in biological invasions remains largely unexplored.

The present study does not test these hypotheses directly. Such analyses would require comparisons between native and invasive populations and, ultimately, species- and strain-level identification combined with culture-based validation, as genera detected here include pathogens, commensals, and mutualists (Hardoim et al. 2015; Grzyb and Szulc 2024). Nevertheless, two hypotheses are particularly relevant to our findings. The conserved pan-tissue functional profile, including colonization, biofertilization, stress tolerance, siderophore production, and competitive exclusion, matches traits expected under the enhanced mutualism hypothesis, although it is equally compatible with commensal interactions. In addition, consistent with pathogen spillback, several genera previously reported as phytopathogens in *A. artemisiifolia* (Bieker et al. 2022), including *Pseudomonas*, *Pantoea*, *Curtobacterium*, and *Ralstonia*, formed part of the conserved endobacterial core identified here. The presence of these genera as stable core members across all plant tissues, in the absence of disease symptoms, shows that the genera that include known phytopathogens are maintained within the *A. artemisiifolia* endobacteriome.

Each of these hypotheses, however, must begin somewhere. Our results provide a starting point: the *A. artemisiifolia* endobacteriome across two countries, three geographic regions, and contrasting habitats harbours a conserved taxonomic and predicted functional core across all plant compartments, complemented by compartment-specific indicator taxa and functional profiles. The pan-tissue profile suggests a functionally integrated endophytic community maintained throughout the plant.

The seed endobacteriome deserves particular attention in the context of invasion biology. As an annual species, *A. artemisiifolia* relies entirely on seed production and dispersal for population persistence and range expansion. It makes seeds the vehicle for not only the plant genome but also its associated microbiome. During the transport stage of invasion, endophyte diversity is expected to be low, as most taxa are not transmitted vertically (Bard et al. 2024). Our data are consistent with this prediction: seeds exhibited the lowest alpha diversity, the smallest core, and no seed-exclusive functional indicators, with the predicted functional profile defined almost entirely by the pan-tissue profile. For an annual invasive species where each generation begins and ends with seed, this microbial inoculum, potentially exhibiting intergenerational stability (Qi et al., 2026), may represent an overlooked yet important element of invasion success.

Comparisons with bulk soil showed that **the endobacteriome is not a random subset of surrounding microbial communities.** Across taxonomic, indicator, differential abundance, and predicted functional analyses, **endophytic communities were distinct from soil**, indicating strong host filtering during colonization or alternative pathways. This study provides the first field-based multi-compartment characterization of the *A. artemisiifolia* endobacteriome, in the European invasive range, and contributes to laying a foundation for future studies on the role of endophytes in plant invasion.

## 5. Conclusions

This study constitutes the first high-throughput sequencing characterization of the bacterial endobacteriome of *Ambrosia artemisiifolia* across leaves, roots, and seeds within its European invasive range, complemented by quantitative PCR and PICRUSt2-based functional prediction. Plant compartment was the dominant factor structuring endobacterial communities, whereas no consistent effect of geographic region or habitat type was detected. The combination of core microbiome, indicator species, and differential abundance analyses identified a generally consistent pan-tissue endobacteriome composed of both genera shared with soil and a plant-specific fraction, with *Gammaproteobacteria* dominating *Alphaproteobacteria* in plant tissues in contrast to surrounding soil.

Concordance between taxonomic and functional community structure was statistically significant. The predicted functional profile indicated a conserved set of functions associated with plant colonization, competitive interactions, stress tolerance, and plant-beneficial traits. Roots and leaves did not differ significantly in predicted functional beta diversity, despite significant taxonomic differentiation, corresponding to partial functional redundancy. Roots were the most taxonomically heterogeneous compartment, with root-exclusive indicators dominated by *Actinomycetes* and the largest compartment-exclusive functional pattern. Leaves were the most compositionally stable compartment, with the largest exclusive core but no exclusive taxonomic indicator; a distinct leaf+soil indicator pattern – the largest in the dataset – suggested alternative recruitment pathways not reflected in the functional analysis. In contrast, seeds harboured the least diverse and most functionally reduced endobacteriome, representing a filtered subset of the broader plant-associated microbiome, agreeing with a transmission bottleneck. Variability among seed samples was pronounced, ranging from near-monodominance to substantially higher evenness. As an annual species dependent on seed-mediated dispersal, *A. artemisiifolia* most probably transfers its endophytic inoculum via its seeds. Therefore, characterizing this inoculum across broader populations should be treated as an immediate priority.

The conserved core included genera encompassing mutualists, commensals, and known phytopathogens, highlighting that genus-level amplicon sequencing cannot resolve ecological function. These results establish a quantitative multi-compartment baseline for the *A. artemisiifolia* endobacteriome and provide a basis for testing microbiome-mediated hypotheses of plant invasion. From our perspective, this includes enhanced mutualism, enemy release, and pathogen spillback, through comparative studies of native and invasive populations, temporal sampling across developmental stages, and fungal microbiome profiling.

## 6. Declarations

### Ethics approval and consent to participate

Not applicable

### Consent for publication

Not applicable

### Availability of data and materials

All analyzed sequences are deposited in SRA, under project No. PRJNA1454844.

### Authors’ contributions (CRediT authorship contribution statement)

Conceptualization – T.G., Data curation – T.G., Formal analysis – T.G., Funding acquisition – T.G., J.S., Investigation – T.G., M.H., B.T-G., Methodology – T.G., Project administration – T.G., J.S., Resources – J.S., Software – T.G., Supervision – J.S., Validation – T.G., Visualization – T.G., Writing – original draft – T.G., J.S., Writing – review and editing – J.S., T.G., B.T-G., M.H.

## Acknowledgements

This article was completed while the first author was a Doctoral Candidate at the Interdisciplinary Doctoral School at Lodz University of Technology, Poland.

## Supplementary materials

Supplementary Figures: S1-S9

Supplementary Tables: S1-S17

Supplementary Methods: S1-S5

Supplementary File S1

