## Supplementary Figures for "There are cores, indicators, and differences – a high-throughput and quantitative PCR study of the *Ambrosia artemisiifolia* L. leaf, root, and seed endophytic bacterial microbiomes within the European invasive range"

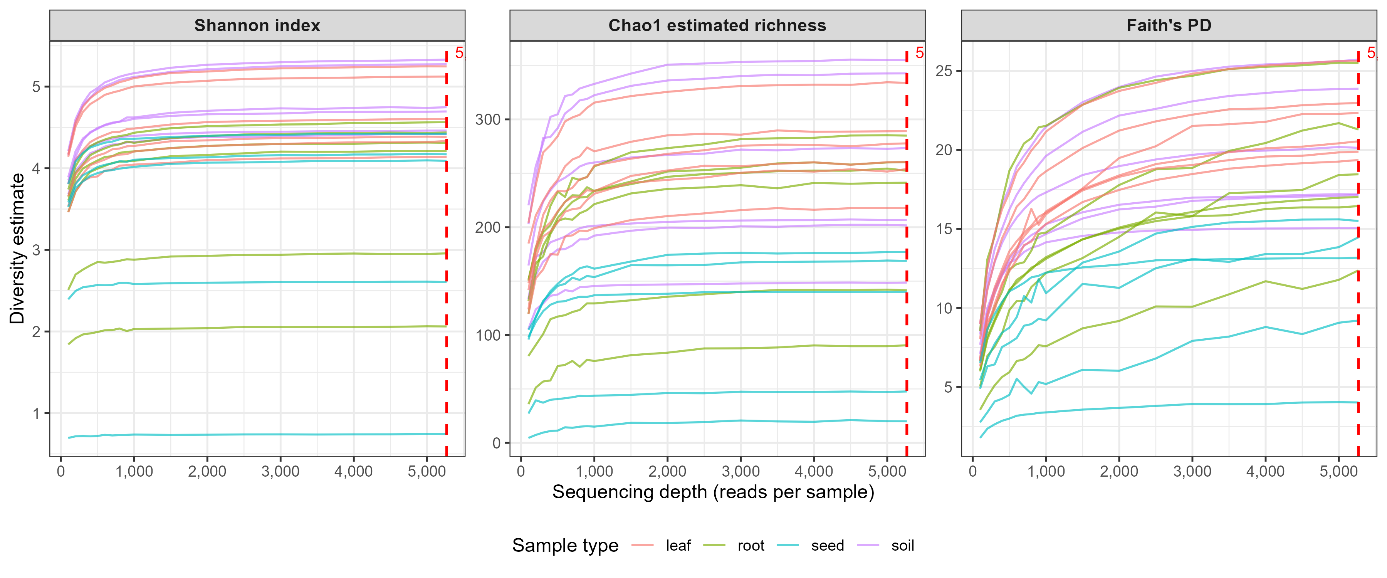

**Figure S1.** Rarefaction curves for three alpha diversity metrics (Shannon index, Chao1 estimated richness, and Faith's phylogenetic diversity) computed for all 23 samples across sequencing depths from 100 to 5,268 reads. Each curve represents a single sample (colored by sample type: leaf, root, seed, soil) and shows the mean diversity estimate across 50 independent rarefaction iterations per depth. The dashed vertical line marks the rarefaction depth of 5,268 reads used in the primary alpha diversity analysis (corresponding to the smallest post-filtering library). Curves reach a plateau well below this depth in all sample types, which supports the chosen rarefaction depth for unbiased alpha diversity comparison.

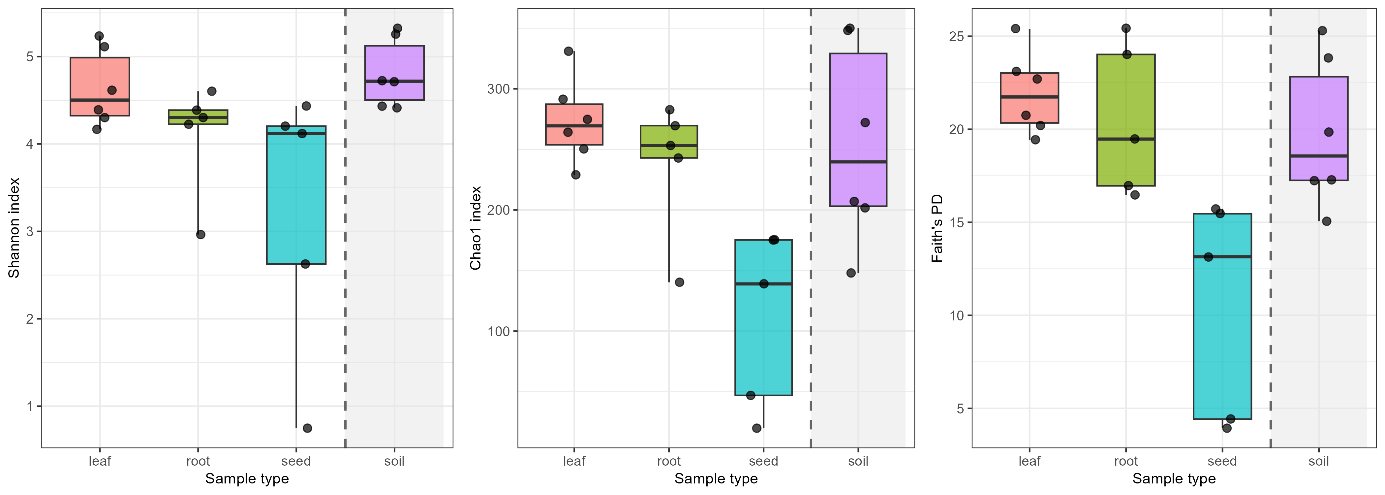

**Figure S2**. Alpha diversity of bacterial communities across sample types (dataset excluding K1; n = 5 for root and seed, n = 6 for leaf and soil). Boxplots with individual data points for (a) Shan-non index, (b) Chao1 estimated richness, and (c) Faith's phylogenetic diversity (PD).

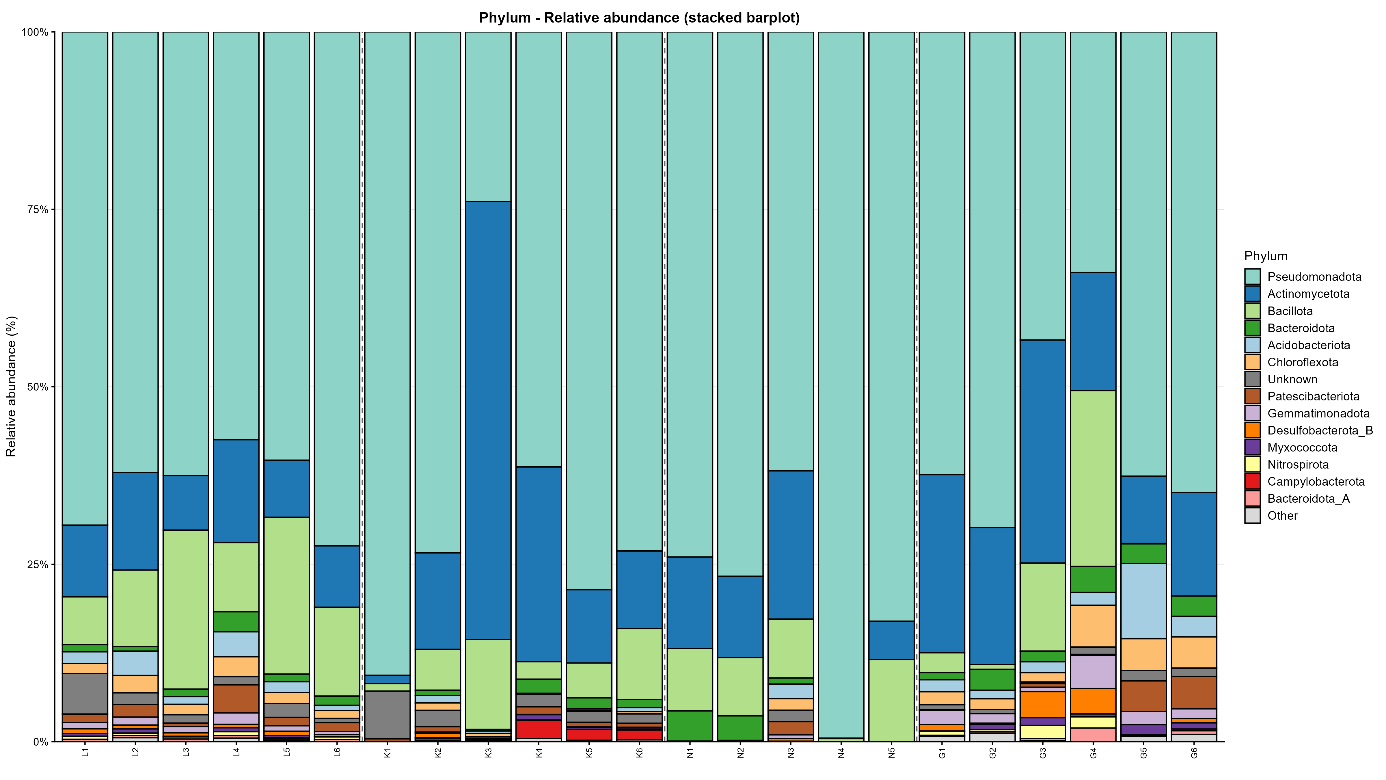

**Figure S3.** Relative abundance of bacterial phyla across sample types. Stacked barplot showing per-sample compositional profiles at the phylum level. Taxa with mean relative abundance ≥0.5% in at least one sample type are displayed; remaining taxa are collapsed into "Other". Dashed vertical lines separate tissue compartments (leaf, root, seed, soil). Samples are ordered by tissue type and sample number.

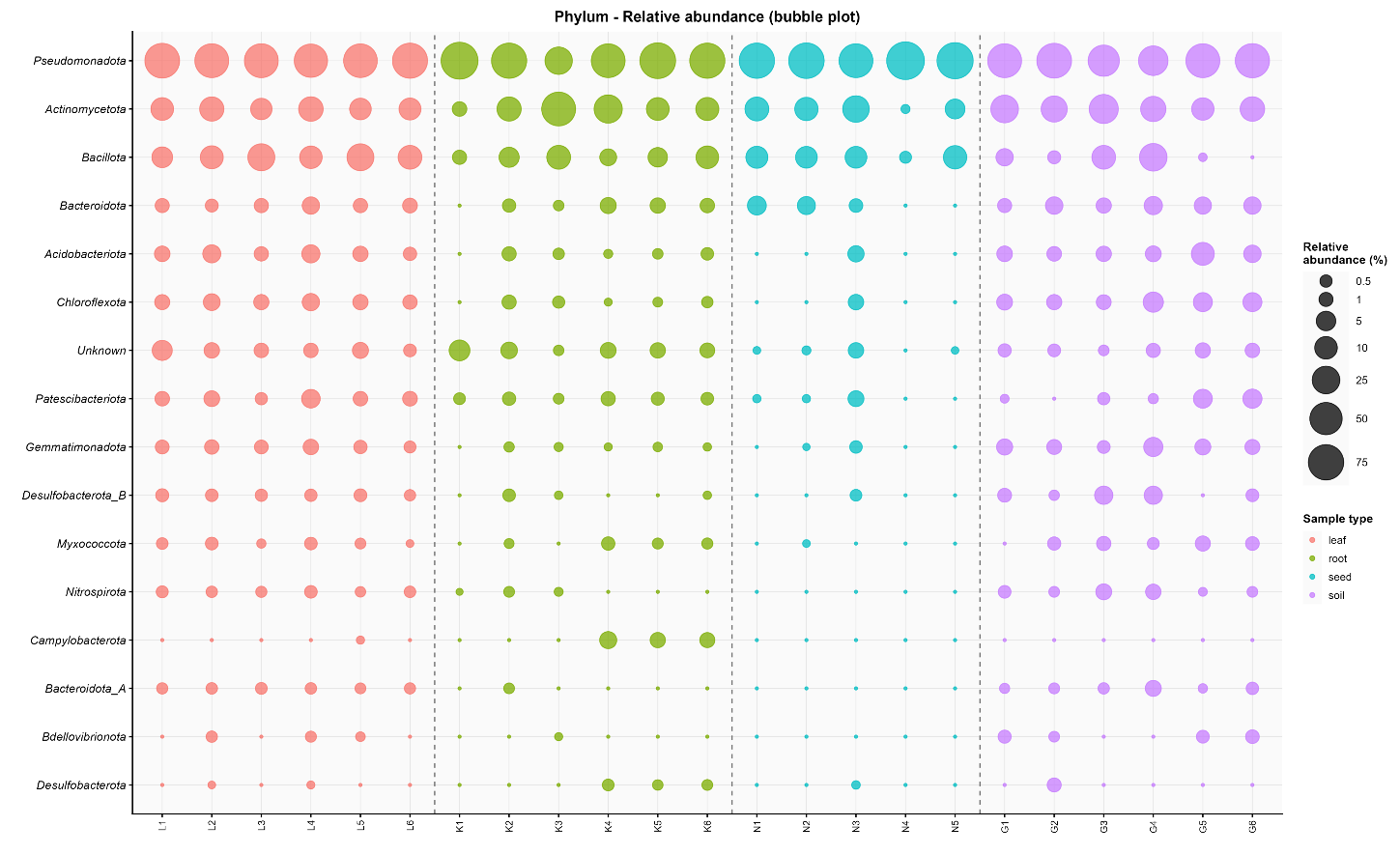

(a)

(b)

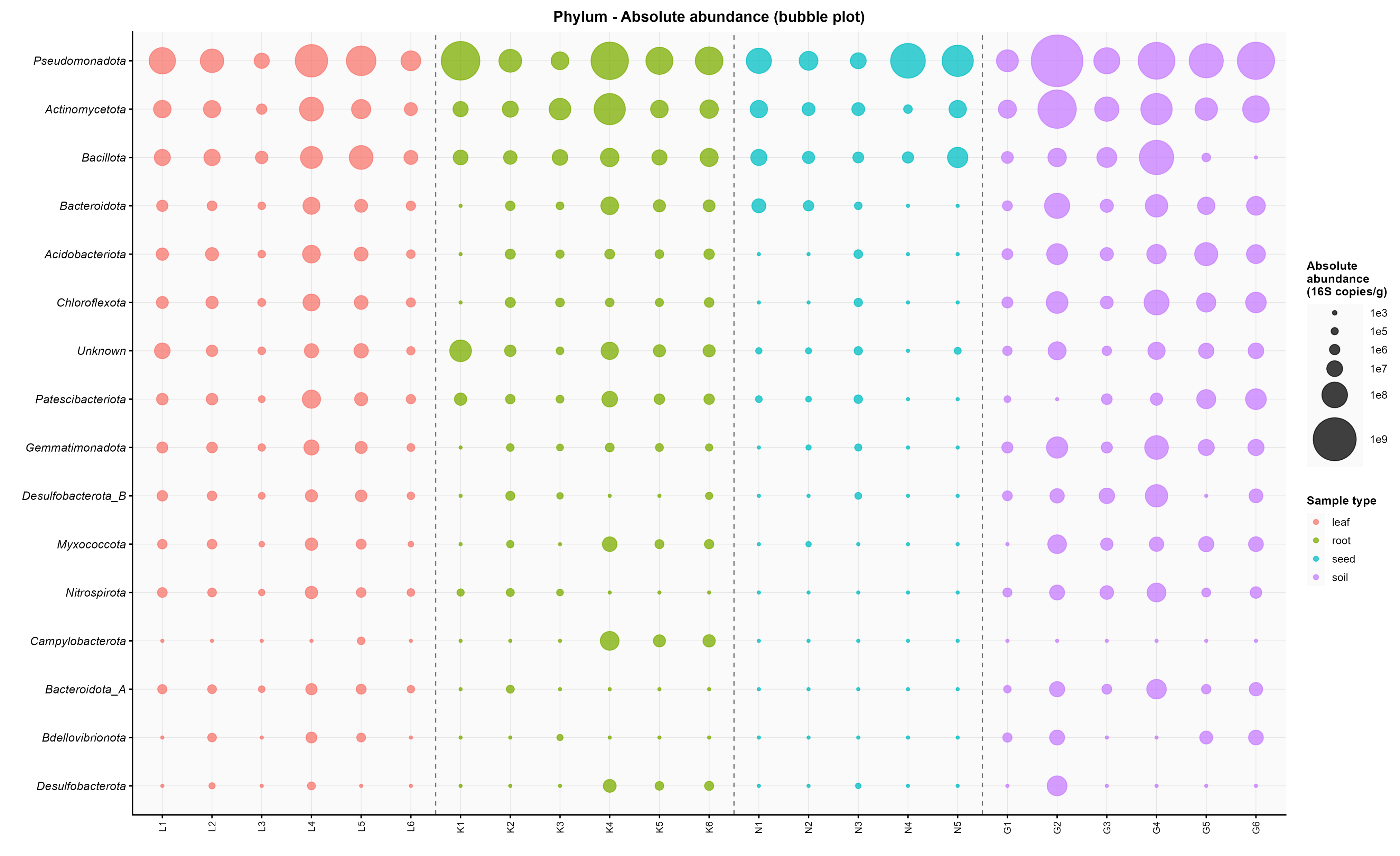

**Figure S4.** Phylum-level bubble plots comparing relative and absolute bacterial abundances. (a) Relative abundance (%; square-root-scaled bubble size) and (b) absolute abundance (16S rRNA gene copies g^-1^ dry weight; square-root-scaled bubble size) for a union of the top 15 phyla by mean relative abundance and the top 15 by total absolute abundance. Phyla on the y-axis are ordered by descending mean relative abundance. Bubble colour indicates sample type. Dashed vertical lines separate tissue compartments.

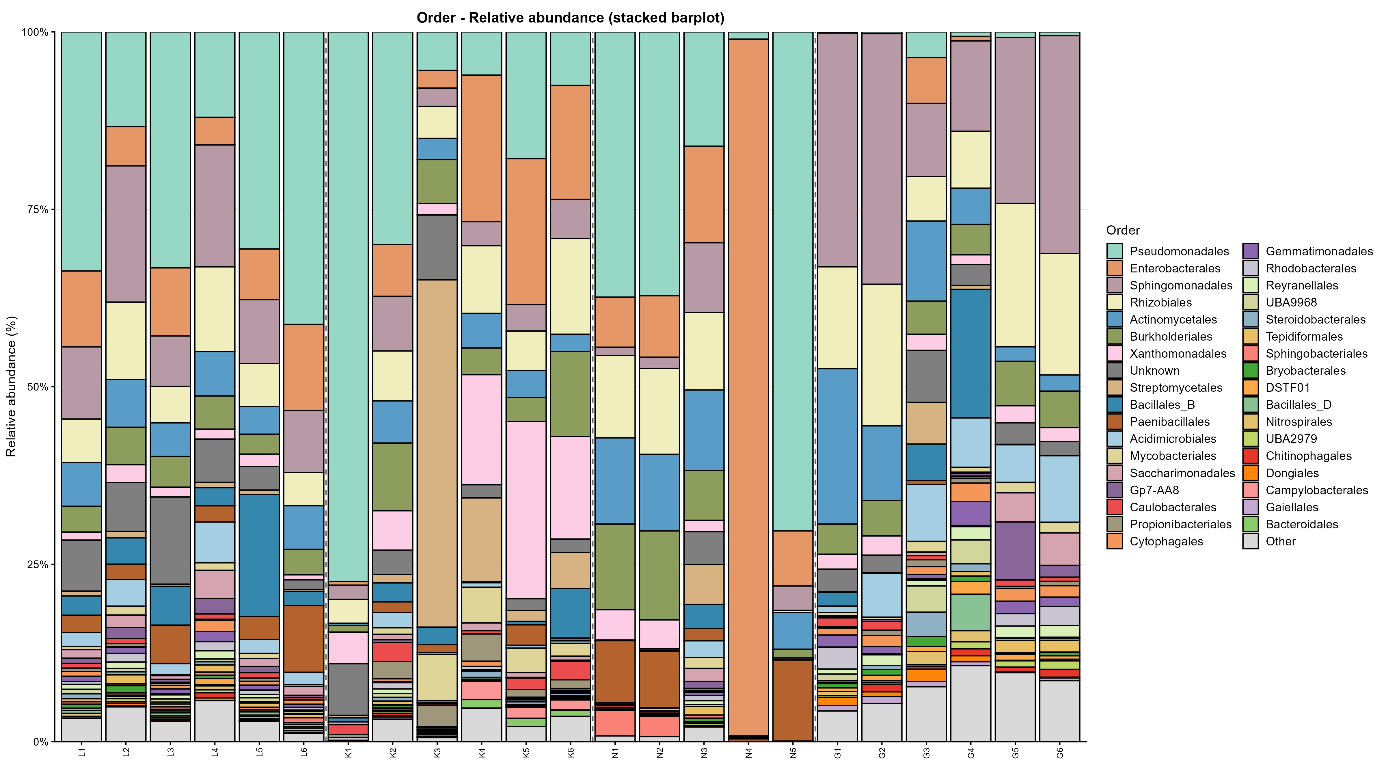

**Figure S5.** Relative abundance of bacterial orders across sample types. Stacked barplot showing per-sample compositional profiles at the order level. Taxa with mean relative abundance ≥0.5% in at least one sample type are displayed; remaining taxa are collapsed into "Other". Dashed vertical lines separate tissue compartments (leaf, root, seed, soil). Samples are ordered by tissue type and sample number.

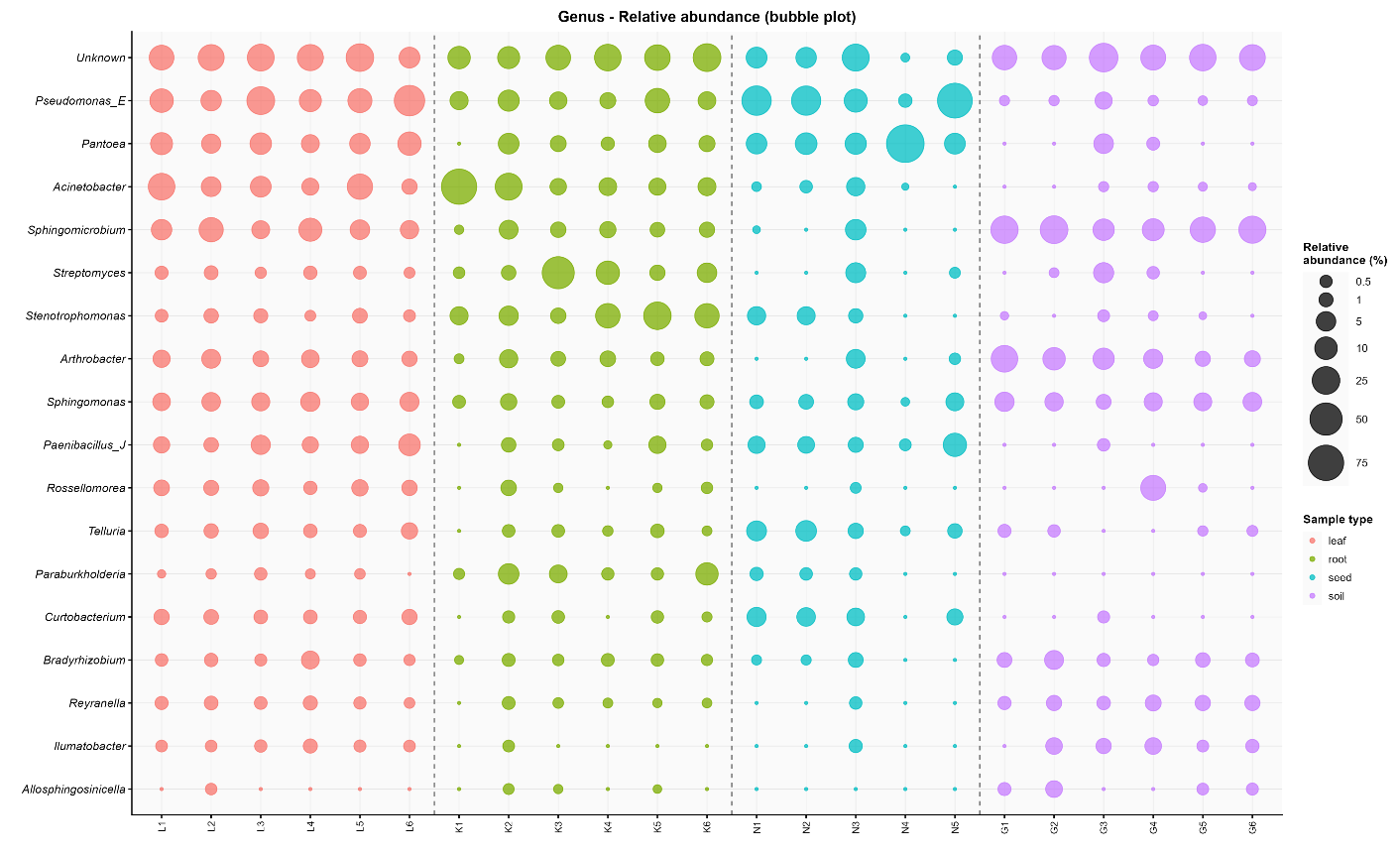

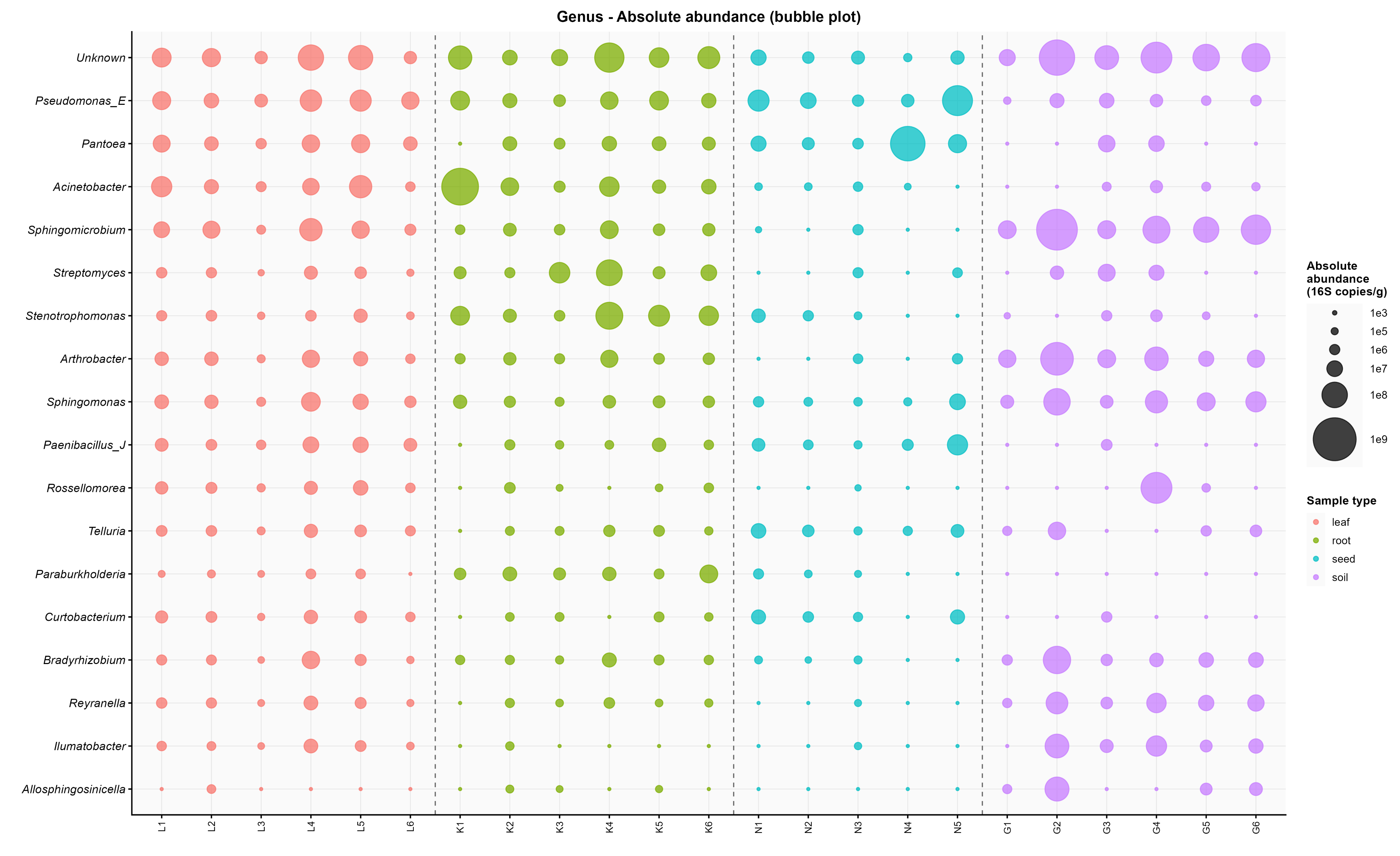

(a)

(b)

**Figure S6.** Genus-level bubble plots comparing relative and absolute bacterial abundances. (a) Relative abundance (%; square-root-scaled bubble size) and (b) absolute abundance (16S rRNA gene copies g^-1^ dry weight; square-root-scaled bubble size) for a union of the top 15 genera by mean relative abundance and the top 15 genera by total absolute abundance. Genera on the y-axis are ordered by descending mean relative abundance. Bubble colour indicates sample type. Dashed vertical lines separate tissue compartments.

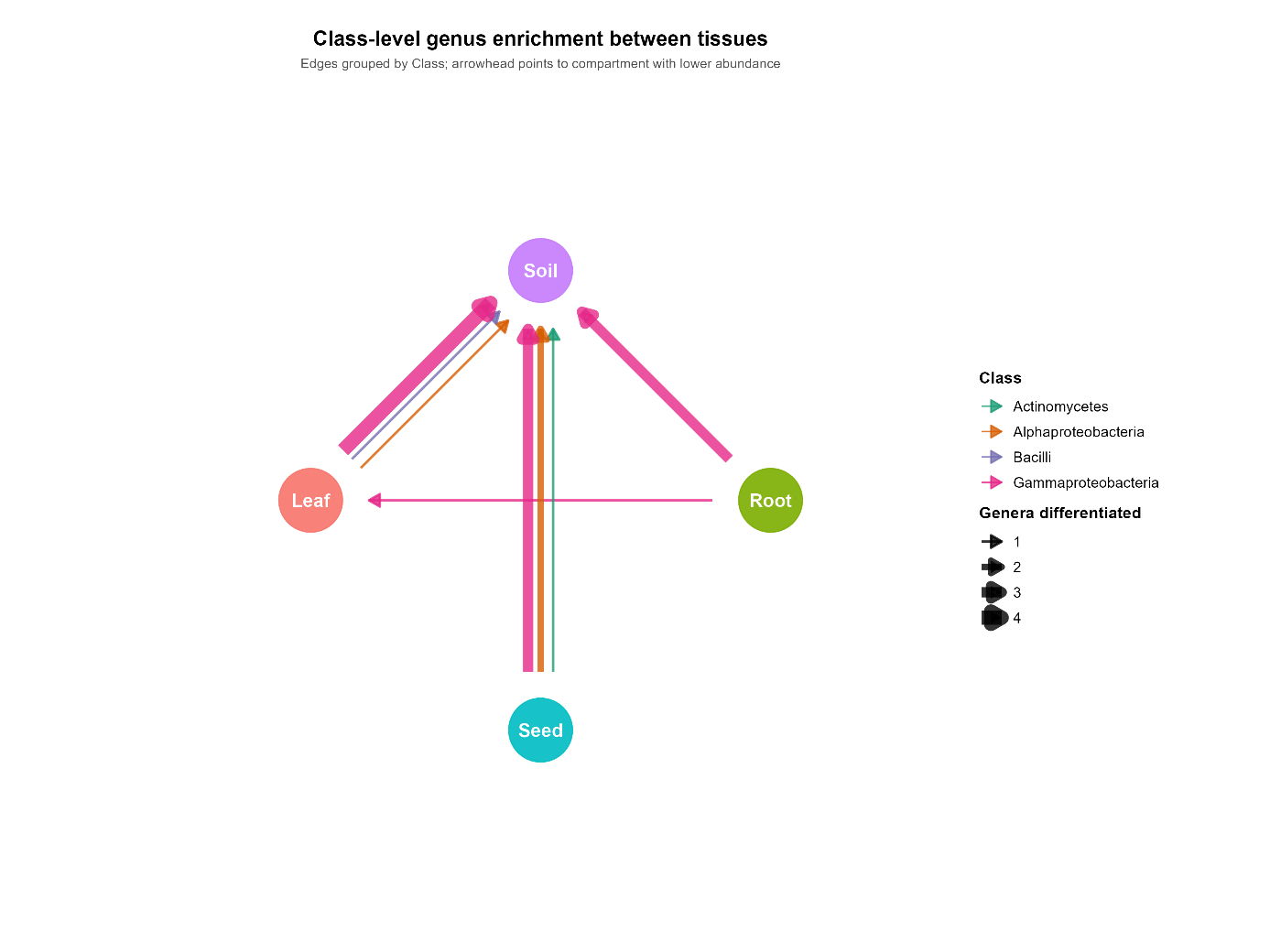

**Figure S7.** Network visualization of pairwise differential abundance among genera between sample types, aggregated to taxonomic Class level (primary dataset excluding K1). Nodes represent the four sample types (soil, leaf, root, seed). Edges represent significant differential abundance between compartment pairs (ANCOM-BC2, BH-adjusted p < 0.05) at the genus level, grouped by their parent Class. Edge direction (arrowhead) points from the compartment with higher mean abundance to the compartment with lower abundance for each Class. Edge thickness is proportional to the number of constituent genera differentiating each pair within each Class. Edge color encodes the parent Class. Class entries with tied directions in a given pair (equal numbers of genera higher in each compartment) are not represented as edges; full pairwise differential abundance results, including tied directions and per-genus statistics, are reported in Table S10.

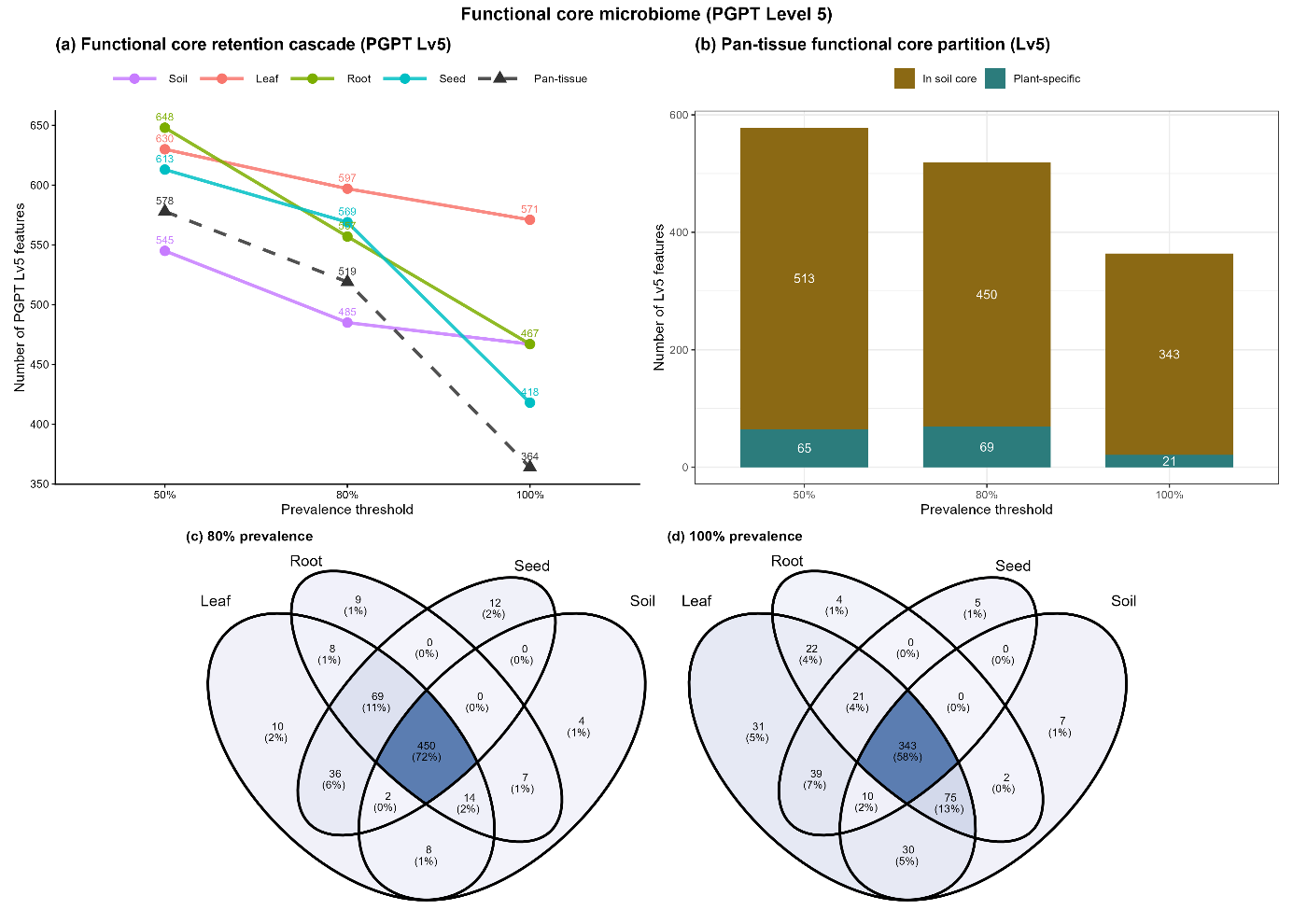

**Figure S8.** Functional core microbiome at PGPT Level 5. (a) Retention cascade across prevalence thresholds (50%, 80%, 100%) for each compartment (soil, leaf, root, seed) and the pan-tissue intersection (leaf+root+seed). Numerical values are shown next to each point. Note: y-axis range starts at 350 to enhance visibility of differences across compartments and thresholds (b) Pan-tissue partition showing the proportion of pan-tissue functional core features shared with the soil core at the same prevalence threshold ("Shared with soil core") versus features exclusive to plant tissues ("Plant-specific"). (c) Four-set Venn diagram of compartment cores at the 80% prevalence threshold. (d) Four-set Venn diagram of compartment cores at the 100% prevalence threshold. All analyses based on the primary dataset excluding K1 (n = 22).

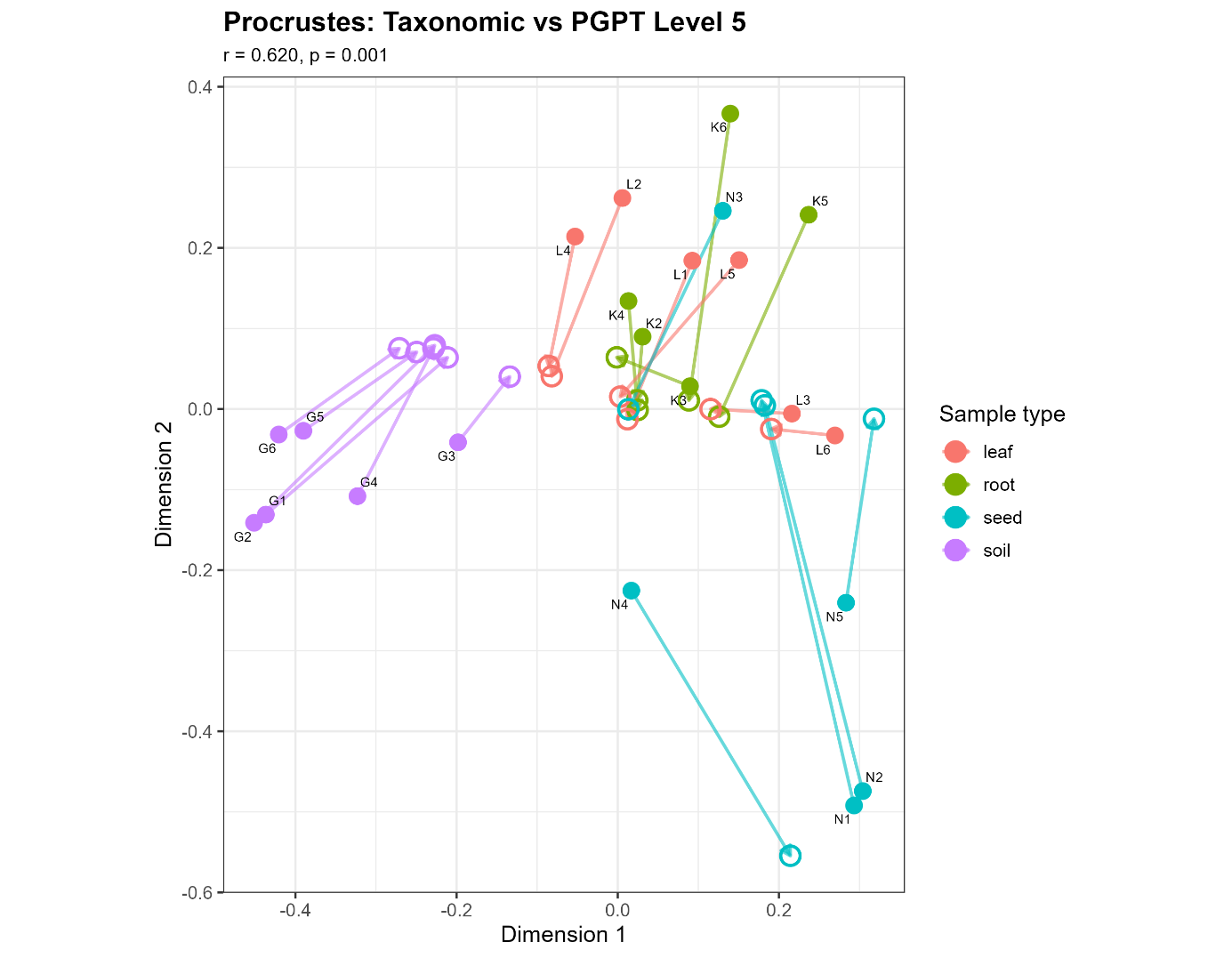

**Figure S9.** Procrustes superimposition of taxonomic and PGPT Level 5 functional Bray–Curtis PCoA ordinations (k = 2), primary dataset excluding K1 (n = 22). Filled symbols represent taxonomic ordination positions; open symbols connected by arrows indicate the corresponding PGPT Level 5 functional positions after symmetric Procrustes rotation and scaling. Symbol shape/colour denotes sample type (leaf, root, seed, soil).
