## Supplementary Methods for "There are cores, indicators, and differences – a high-throughput and quantitative PCR study of the *Ambrosia artemisiifolia* L. leaf, root, and seed endophytic bacterial microbiomes within the European invasive range"

### Methods S1 Sampling

#### S1.1 Phase 1 sampling (early flowering)

Phase 1 sampling was conducted in early August 2025. At each site, 7–10 mature *A. artemisiifolia* individuals at the early flowering stage were randomly selected to capture within-site heterogeneity. The following materials were collected per individual:

- **Leaves**: the second, third, and fourth leaves from the apex were collected using clean pruning shears disinfected with 70% ethanol between individuals.
- **Taproot**: a 10 cm middle section of the taproot was excised using disinfected pruning shears.
- **Bulk soil**: collected at a distance from the plant in plant-free spaces, 0-10 cm depth. Bulk soil served as an environmental reference and was stored without surface sterilization.

Samples were immediately placed in plastic or paper bags and transported at 4 °C between cooling blocks to the nearest location where surface sterilization procedures were possible. All working surfaces and tools were disinfected with 70% ethanol prior to and between samples.

#### S1.2 Phase 2 sampling (seed maturity)

Phase 2 sampling was conducted in October – early November 2025, depending on local seed ripeness. Mature seeds (dry, brown achenes) were collected from reproductively mature plants at five of the six sites. Seed collection was not possible at Kuźnica Warężyńska, where plants had been mowed by road maintenance authorities prior to seed maturation. Seeds were transported to the laboratory in dry conditions.

#### S1.3 Surface sterilization protocols

Surface sterilization followed Sahu et al. (2022) with tissue-specific adjustments:

**Leaves:**

1. Immersion in 70% ethanol for 30 s
2. Three rinses in sterile distilled water
3. Immersion in 2% NaClO solution for 2 min
4. Five rinses in sterile distilled water

**Roots:**

1. Immersion in 70% ethanol for 30 s
2. Three rinses in sterile distilled water
3. Immersion in 3% NaClO solution for 3 min
4. Five rinses in sterile distilled water

**Seeds:**

1. Immersion in 70% ethanol for 30 s
2. Three rinses in sterile distilled water
3. Immersion in 5% NaClO solution for 8 min
4. Five rinses in sterile distilled water

#### S1.4 Sample preservation and storage

Surface-sterilized tissues were immediately placed in DESS preservative (20% DMSO, 0.25 M EDTA, saturated NaCl) following Carvalhais et al. (2022). Bulk soil was placed in sterile containers without sterilization. All samples were transported to the laboratory and stored at −80 °C until DNA extraction.

### Methods S2 DNA isolation

#### S2.1 Sample preparation

Tissue samples (leaves, roots, seeds) preserved in DESS solution and stored at -80 °C were processed in three independent extraction replicates per biological sample. For each replicate, a sub-sample of preserved tissue (0.4-0.5 g fresh weight) was retrieved from the parent sample and washed sequentially in sterile molecular-grade water (Eurx) to remove protectant residues, then centrifuged (7,000 × g, 5 min). Supernatants were discarded and tissues were transferred to a mortar and ground with a pestle. A separate sub-sample of each parent biological sample was used for dry mass determination (oven-dried at 105 °C to constant weight).

#### S2.2 Mechanical homogenization

Ground tissues were transferred to Eppendorf tubes containing tissue lysis buffer (composition in S2.3) and tissue-specific glass beads:

| **Tissue** | **Bead diameters** | **Mill cycles** | **Cycle duration** | **Speed** |
| --- | --- | --- | --- | --- |
| Leaves | 2.5 + 3.5 mm | 3 cycles | 10 s each | 3.7 m/s |
| Roots | 1.5 + 3.5 + 5 mm | 4 cycles | 15 s each | 4.4 m/s |
| Seeds | 2.5 + 3.5 mm | 4 cycles | 15 s each | 4.4 m/s |

Homogenization was performed using a Mini-Bead Mill (VWR).

#### S2.3 Tissue DNA extraction (modified from Maropola et al., 2015)

Tissue lysis buffer composition:

- 25 mM Tris-HCl pH 8.0
- 10 mM EDTA pH 8.0
- 1% w/v CTAB
- 50 mM sucrose
- 25 mg·mL^-1^ lysozyme

Extraction workflow:

1. **Enzymatic lysis**: addition of proteinase K to 1 mg·mL^-1^ final concentration; incubation at 37°C for 1 h.
2. **Detergent lysis**: addition of SDS to 1% final concentration; incubation at 65 °C for 30 min.
3. **First organic extraction**: addition of equal volume of phenol; mixing and centrifugation; aqueous phase recovery.
4. **Second organic extraction**: addition of equal volume of chloroform:isoamyl alcohol (24:1); mixing and centrifugation; aqueous phase recovery.
5. **Precipitation**: addition of isopropanol; overnight incubation at -20 °C; centrifugation.
6. **Wash**: 70% ethanol; centrifugation.
7. **Resuspension**: TE buffer (pH 8.0, preheated to 65 °C).

#### S2.4 Soil DNA extraction (adjusted Protocol III from Dwivedi & Yadav, 2023)

Approximately 250 mg of soil per replicate was processed. Soil lysis buffer composition:

- 100 mM Tris-HCl pH 8.0
- 100 mM EDTA pH 8.0
- 100 mM CaCl_2_
- 1.5 M NaCl
- 1% CTAB
- 100 mM potassium phosphate buffer pH 8.0
- 10 mg·mL^-1^ lysozyme
- 10 mg·mL^-1^ proteinase K

Extraction workflow:

1. **Enzymatic lysis**: incubation at 37 °C for 1 h.
2. **Detergent lysis**: addition of SDS to 5% final concentration; incubation at 60 °C for 1 h.
3. **First organic extraction**: chloroform:isoamyl alcohol (24:1); aqueous phase recovery.
4. **Selective precipitation**: PEG-6000 to 30% final concentration with 0.1 M sodium acetate pH 5.2; overnight precipitation.
5. **Second organic extraction**: phenol:chloroform:isoamyl alcohol (25:24:1).
6. **Final precipitation**: isopropanol; overnight incubation at -20 °C.
7. **Sequential washes**: 5 M NaCl, then 70% ethanol.
8. **Resuspension**: TE buffer (pH 8.0, preheated to 65 °C).

#### S2.5 Composite DNA preparation and quantification

For each biological sample, the three independent DNA extracts were pooled into one tube per sample. DNA concentration was measured using a Qubit fluorometer with the dsDNA HS assay. The pooled DNA was used for qPCR and library preparation.

### Methods S3 PCR amplification

#### S3.1 Reaction composition

PCR amplification of the bacterial 16S rRNA gene V3–V4 region was performed using primers 322F (5'-ACGGHCCARACTCCTACGGGA-3') and 796R (5'-CTACCMGGGTATCTAATCCKG-3') (Chen et al. 2022).

Reaction composition (25 µL total volume):

- Template DNA: 1 µL (diluted 1:10 with sterile molecular-grade water if PCR inhibition was detected)
- ColorTaq polymerase (EurX): 1.5 U
- 1× EurX PCR buffer
- MgCl_2_: 2 mM final concentration
- BSA: 0.4 µg/µL
- Each primer: 0.2 µM
- dNTPs: 0.2 mM each

#### S3.2 Thermocycler program

| **Step** | **Temperature** | **Duration** | **Cycles** |
| --- | --- | --- | --- |
| Initial denaturation | 95°C | 3 min | 1 |
| Denaturation | 95°C | 30 s | 32 |
| Annealing | 51°C | 30 s |  |
| Extension | 72°C | 30 s |  |
| Final extension | 72°C | 7 min | 1 |

#### S3.3 Verification, purification, and sequencing

PCR products were verified by 2% TBE agarose gel electrophoresis. Products were purified using Agencourt AMPure XP magnetic beads following the manufacturer's protocol and resuspended in Elution Buffer. Library construction, quality assessment using an Agilent 2100 Bioanalyzer, and sequencing were performed by BGI Genomics using the DNBSEQ-G400 platform with paired-end 300 bp reads.

### Methods S4 qPCR – detailed protocol

#### S4.1 Reaction composition

Quantitative PCR was performed using qPCR HS Mix SYBR (including RUN-HS Taq Polymerase; A&A Biotechnology) at 1× final concentration with the following modifications to the manufacturer's protocol:

- MgCl_2_ adjusted to 2.6 mM (vs default formulation)
- BSA added to 0.8 µg/µL final concentration

Reaction composition (25 µL total volume):

- Template DNA: 1 µL (diluted as necessary to reduce PCR inhibition)
- 1× qPCR HS Mix SYBR
- MgCl_2_: 2.6 mM final
- BSA: 0.8 µg/µL
- Each primer (322F, 796R): 0.2 µM

Each sample was measured in triplicate.

#### S4.2 Standard curve generation

Standard curves were generated using serial dilutions of purified Bacillus cereus zielonkawy genomic DNA (NCBI RefSeq GCF_043099295.1).

#### S4.3 Thermocycler program

| **Step** | **Temperature** | **Duration** | **Cycles** |
| --- | --- | --- | --- |
| Initial denaturation | 95°C | 5 min | 1 |
| Denaturation | 95°C | 15 s | 40 |
| Annealing | 49°C | 45 s |  |
| Extension | 72°C | 1 min |  |

Plate reading was performed at the end of each cycle.

#### S4.4 Melting curve analysis

Melting curve analysis was performed from 65 °C to 95 °C with 0.3 °C increments and 5 s hold time per step, to confirm PCR product specificity.

#### S4.5 Performance characteristics

qPCR efficiency ranged from 74-76% across runs, which is within the expected range for the long ~470 bp amplicon and consistent with published values for amplicons >400 bp (Zemb et al. 2020; Van Holm et al. 2021).

#### S4.6 Data analysis

qPCR results were analyzed using Bio-Rad CFX Maestro software, exported to Microsoft Excel, and subsequently imported into R. Standard curve equations were applied to calculate the number of 16S rRNA gene copies per reaction, then normalized to per gram of dry weight using the dry mass measurements **(Methods S2.1)** and corrected for the contaminant and organellar fraction (**Section 2.6.3**). qPCR-derived gene copy numbers were merged with the phyloseq metadata to enable absolute abundance calculations.

### Methods S5 Exclusion of root sample K1

#### S5.1 Rationale

In K1, *Acinetobacter* made up 73.8% of reads, more than any single genus in any other root or leaf sample; only *Pantoea* in seed sample N4 was higher (98.1%). Unlike Pantoea – a well-established seed endophyte (Nelson 2018) whose monodominance in seeds is consistent with documented seed microbiome variability ranging from near-monodominance to communities of thousands of taxa (Simonin et al. 2022; Abdelfattah et al. 2023) – Acinetobacter has been repeatedly identified as a reagent and laboratory contaminant across multiple independent studies (Salter et al. 2014; Eisenhofer et al. 2019). Although Acinetobacter is also a recognised component of the plant microbiome (Bai et al. 2015), consistent with its detection at moderate levels in other leaf and root samples in this dataset, its dominance in K1 was without precedent in the remaining 22 samples. For comparison, the second-highest single-genus dominance in roots was Streptomyces at 48.9% in K3 – a taxon not listed among common laboratory contaminants (Salter et al. 2014; Eisenhofer et al. 2019) and a well-characterised plant root endophyte (Viaene et al. 2016; Worsley et al. 2020), also present at 5.9% in the corresponding soil sample G3. Given this combination of evidence – extreme dominance, known kitome association, and absence of a comparable pattern in any other sample – K1 was excluded from primary analyses as a conservative measure.

#### S5.2 Treatment across analytical layers

| **Analysis** | **Primary dataset** | **K1-inclusive sensitivity** |
| --- | --- | --- |
| Bacterial load (qPCR) | Included | na |
| Alpha diversity | Excluded | + |
| Taxonomic composition visualisation | Included | na |
| PCoA ordination (taxonomic) | Included | na |
| PCoA ordination (functional) | Included | na |
| Beta diversity (PERMANOVA/PERMDISP) | Excluded | + |
| Core microbiome (taxonomic) | Excluded | – |
| Indicator species (taxonomic) | Excluded | + |
| Differential abundance (taxonomic) | Excluded | – |
| Functional beta diversity | Excluded | – |
| Functional core | Excluded | – |
| Functional indicator | Excluded | – |
| Functional differential abundance | Excluded | – |
| Concordance | Excluded | – |

PCoA ordinations and taxonomic composition visualisations include K1 to illustrate its position within the broader sample distribution, but K1 was excluded from all accompanying statistical tests. All remaining analyses were performed on the K1-excluded primary dataset (n = 22). When applicable, parallel K1-inclusive runs are provided in the Supplementary Materials to assess the sensitivity of results to this exclusion.
