## Supplementary figures and images for "There are cores, indicators, and differences – a high-throughput and quantitative PCR study of the *Ambrosia artemisiifolia* L. leaf, root, and seed endophytic bacterial microbiomes within the European invasive range"

### 1_relative_bar_Class.png

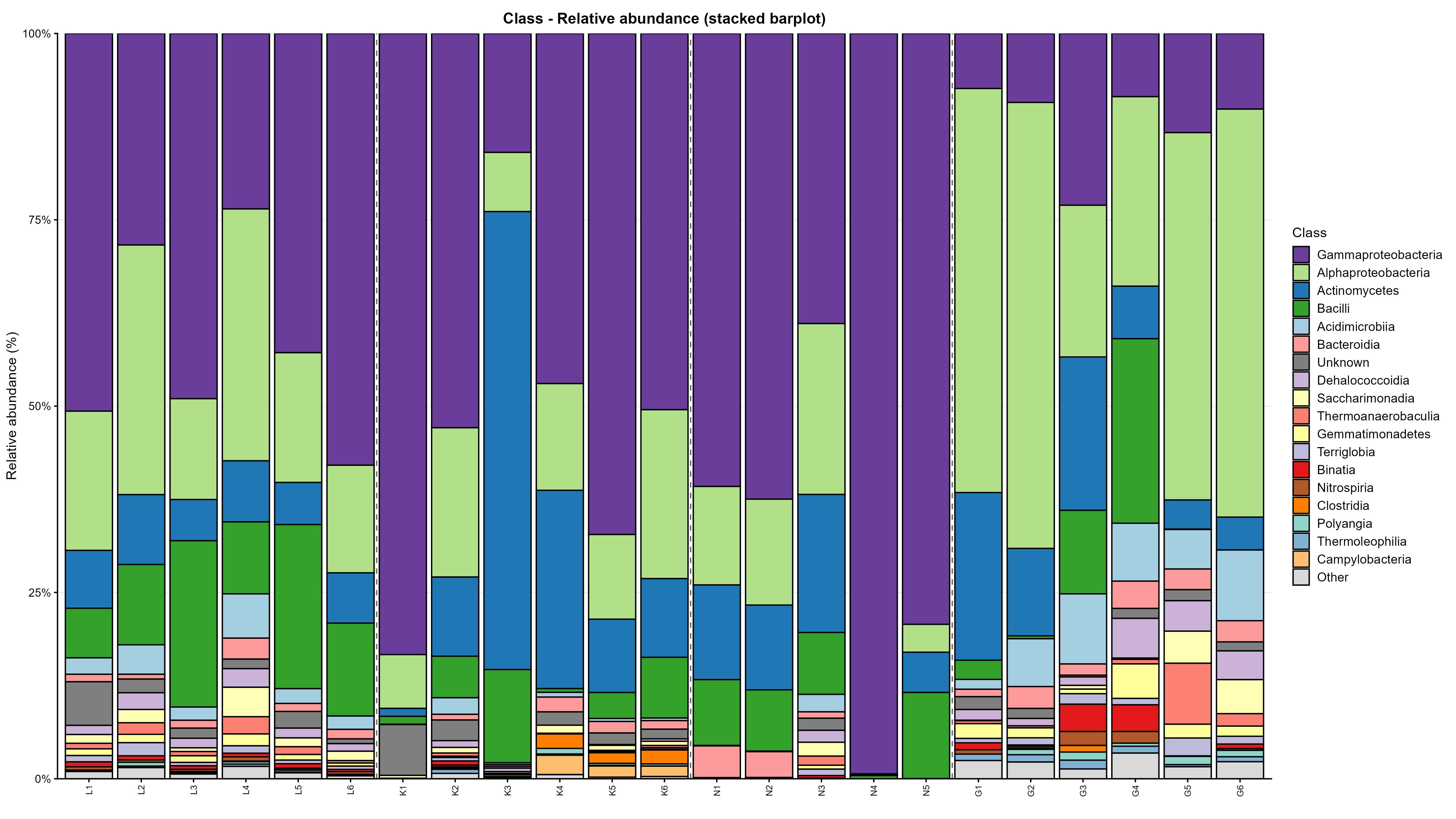

### 1_relative_bar_Family.png

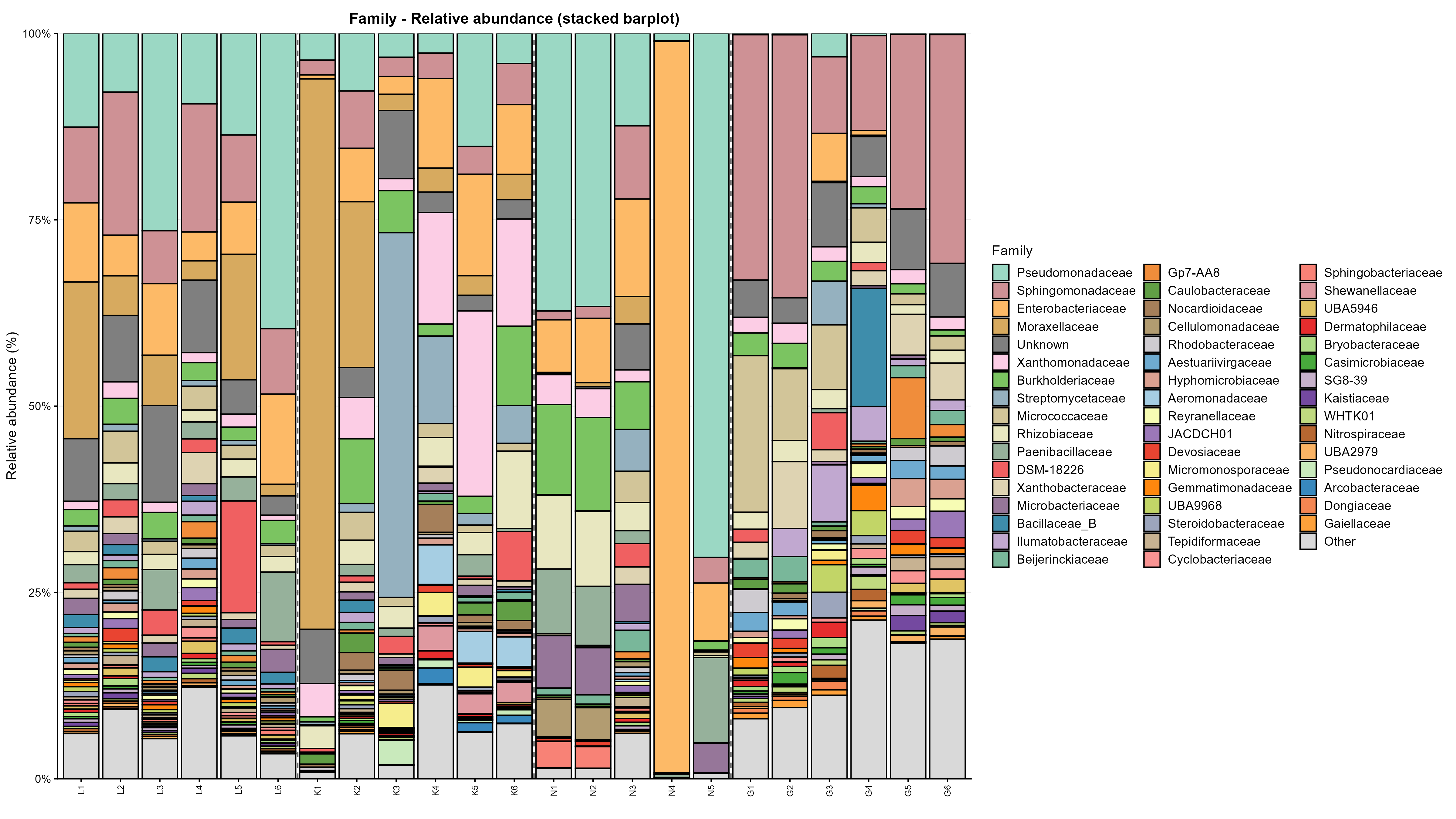

### 1_relative_bar_Genus.png

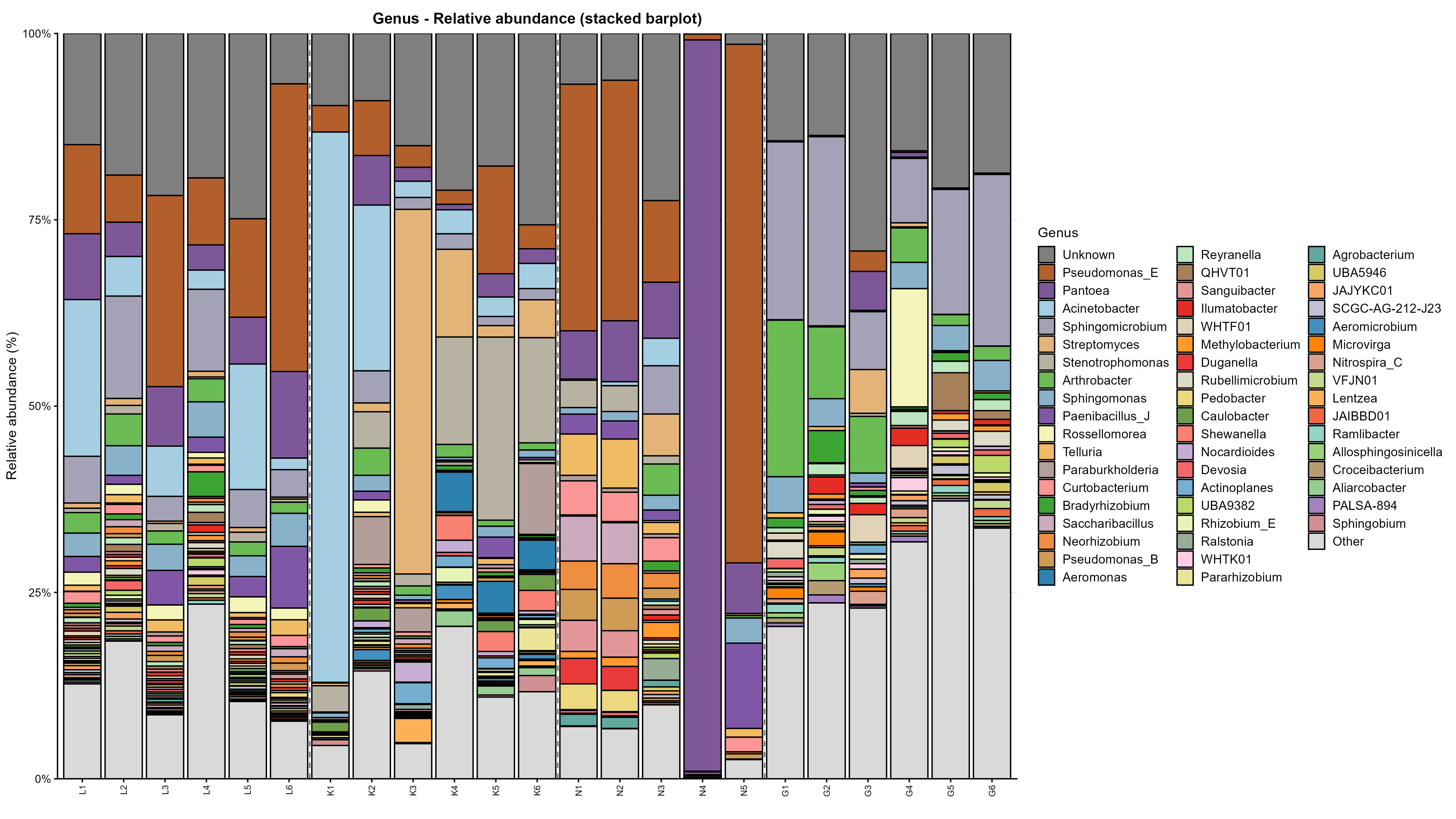

### 1_relative_bar_Order.png

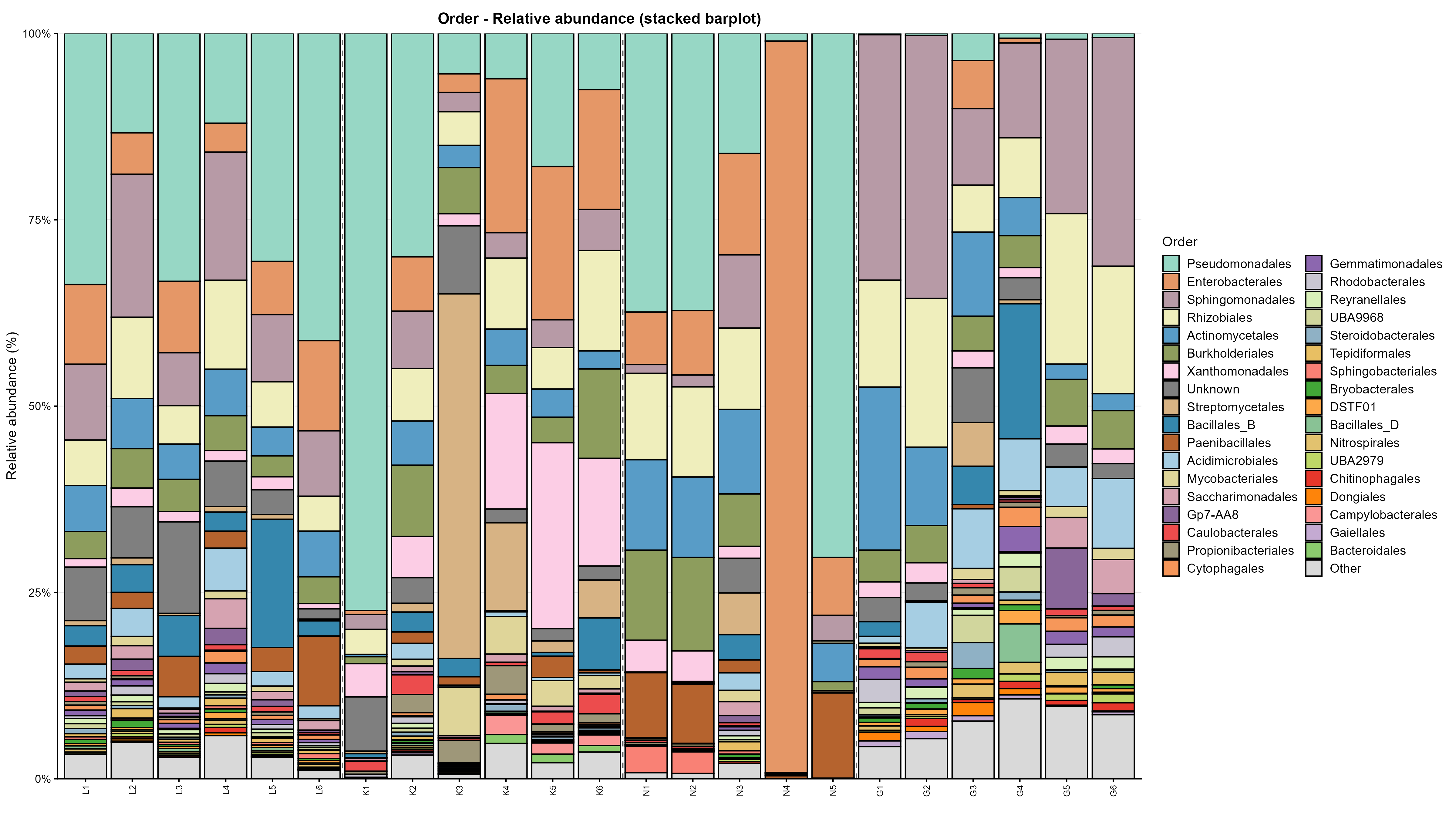

### 1_relative_bar_Phylum.png

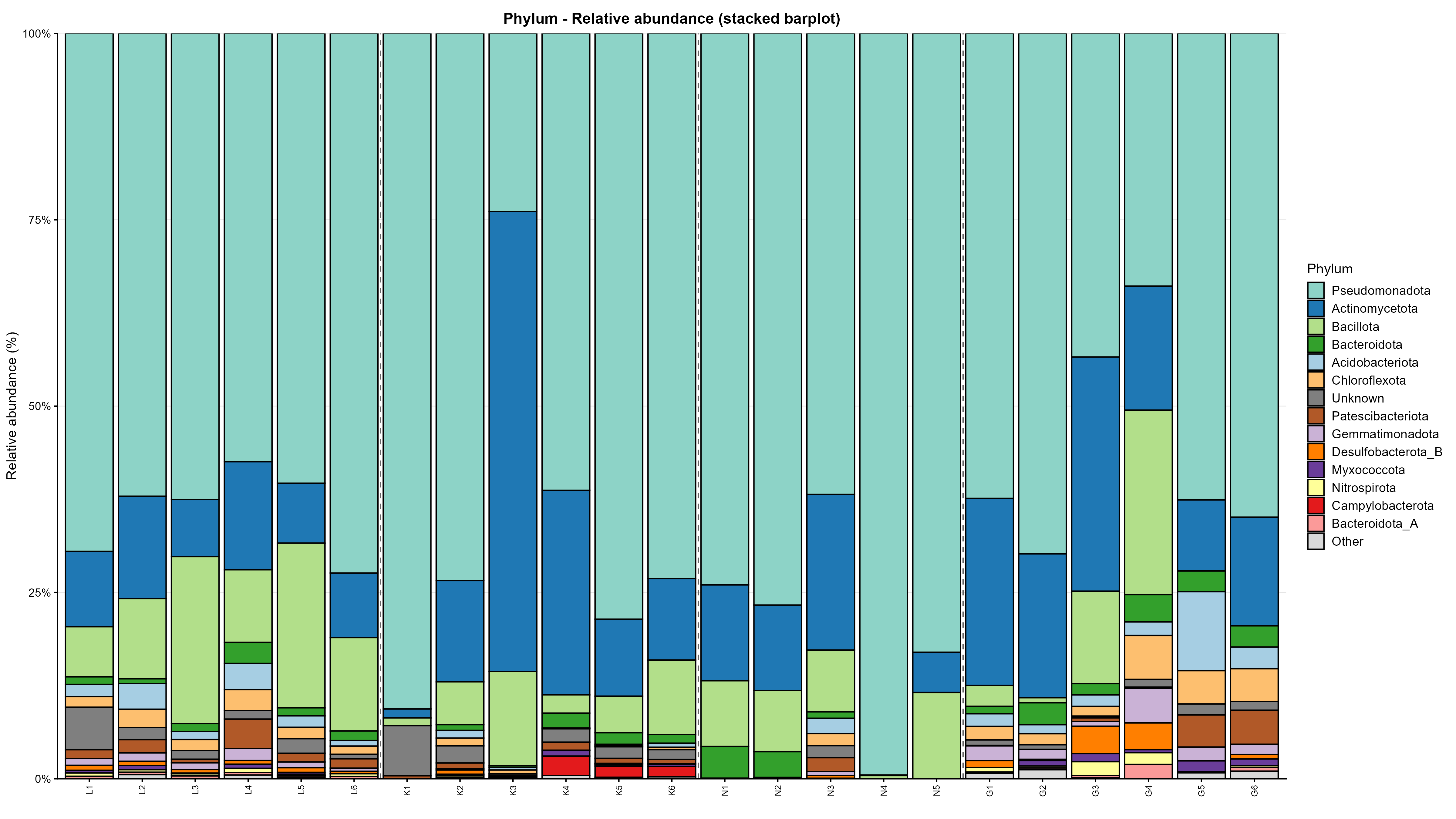

### 2_relative_bubble_Class.png

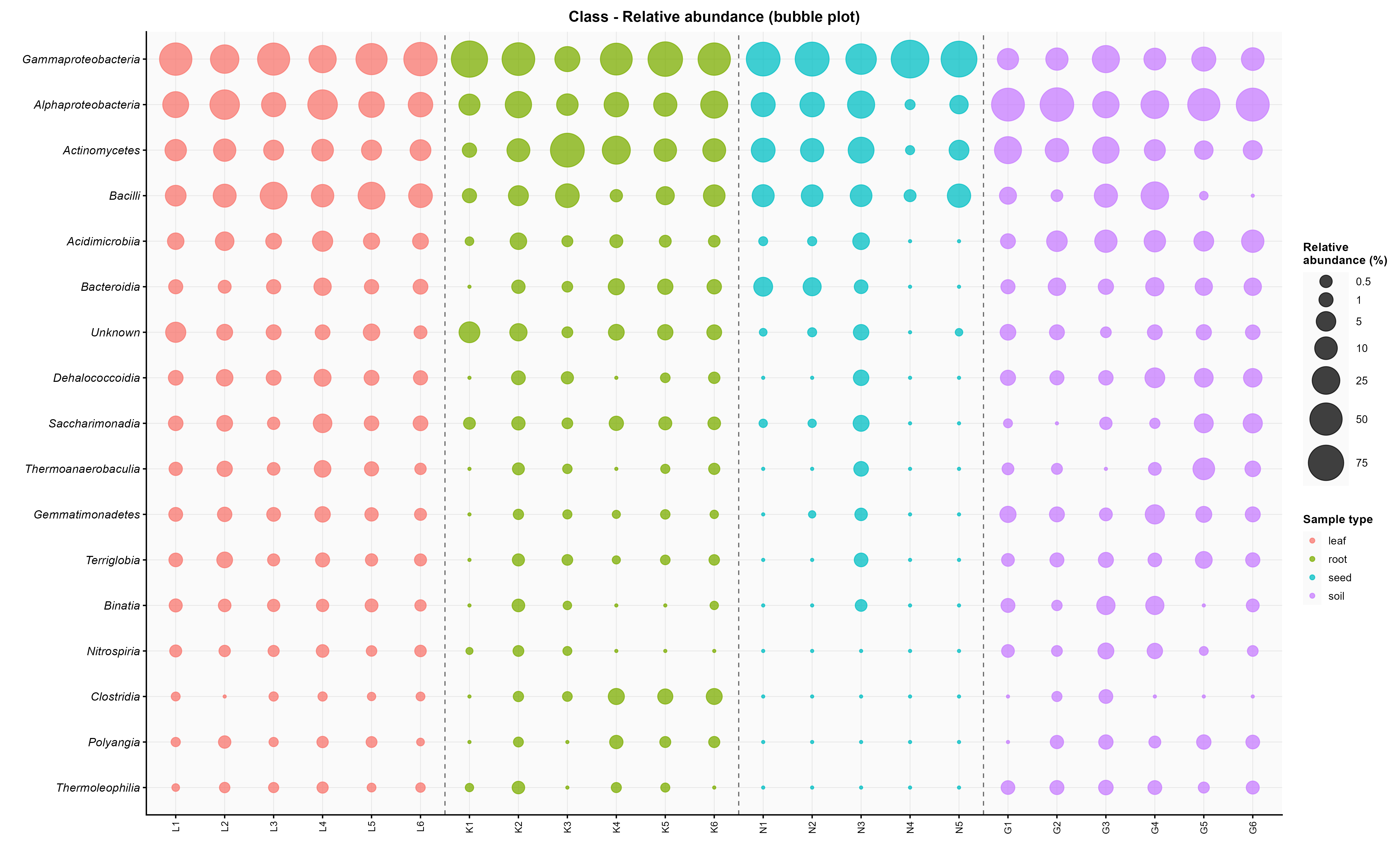

### 2_relative_bubble_Family.png

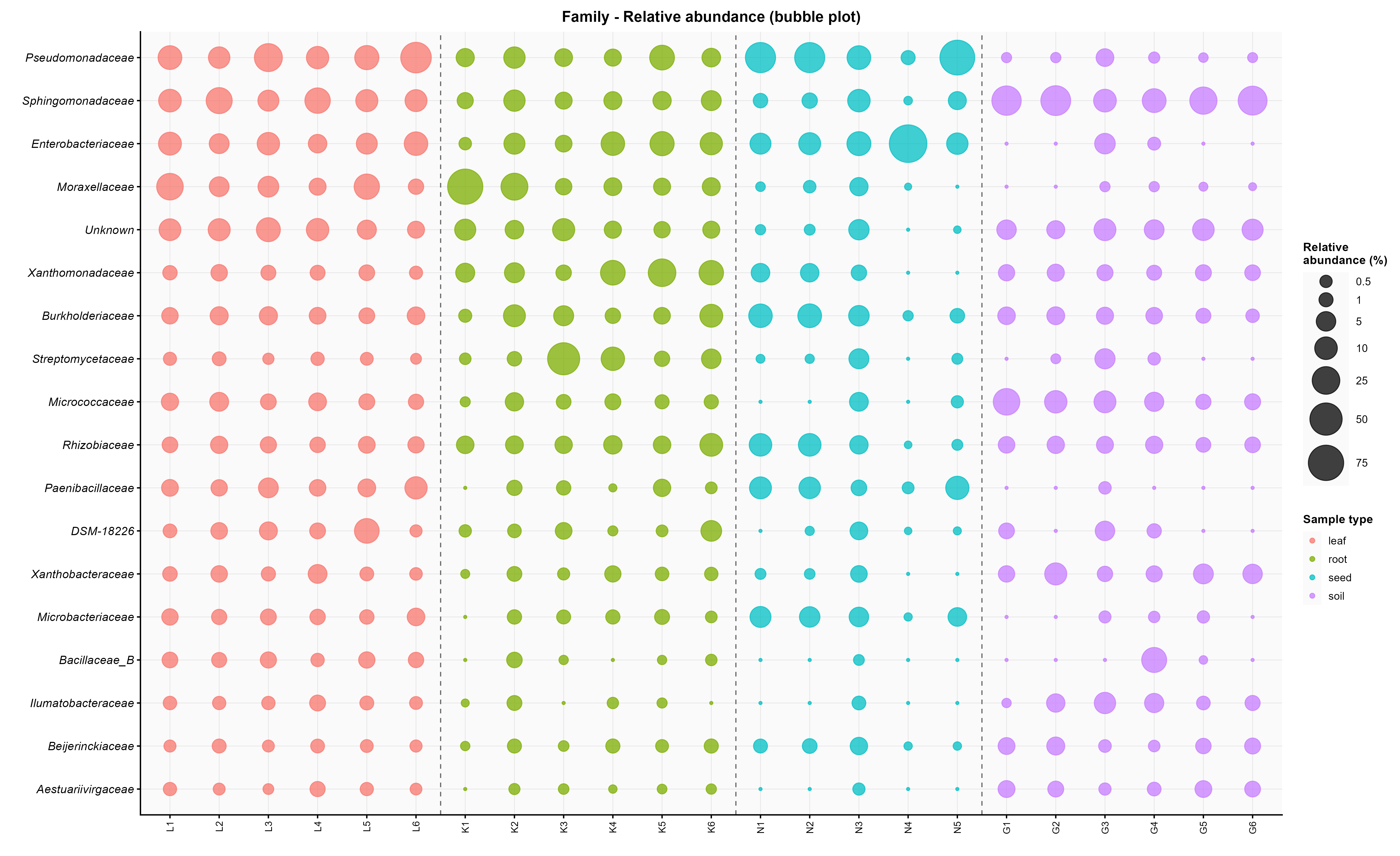

### 2_relative_bubble_Genus.png

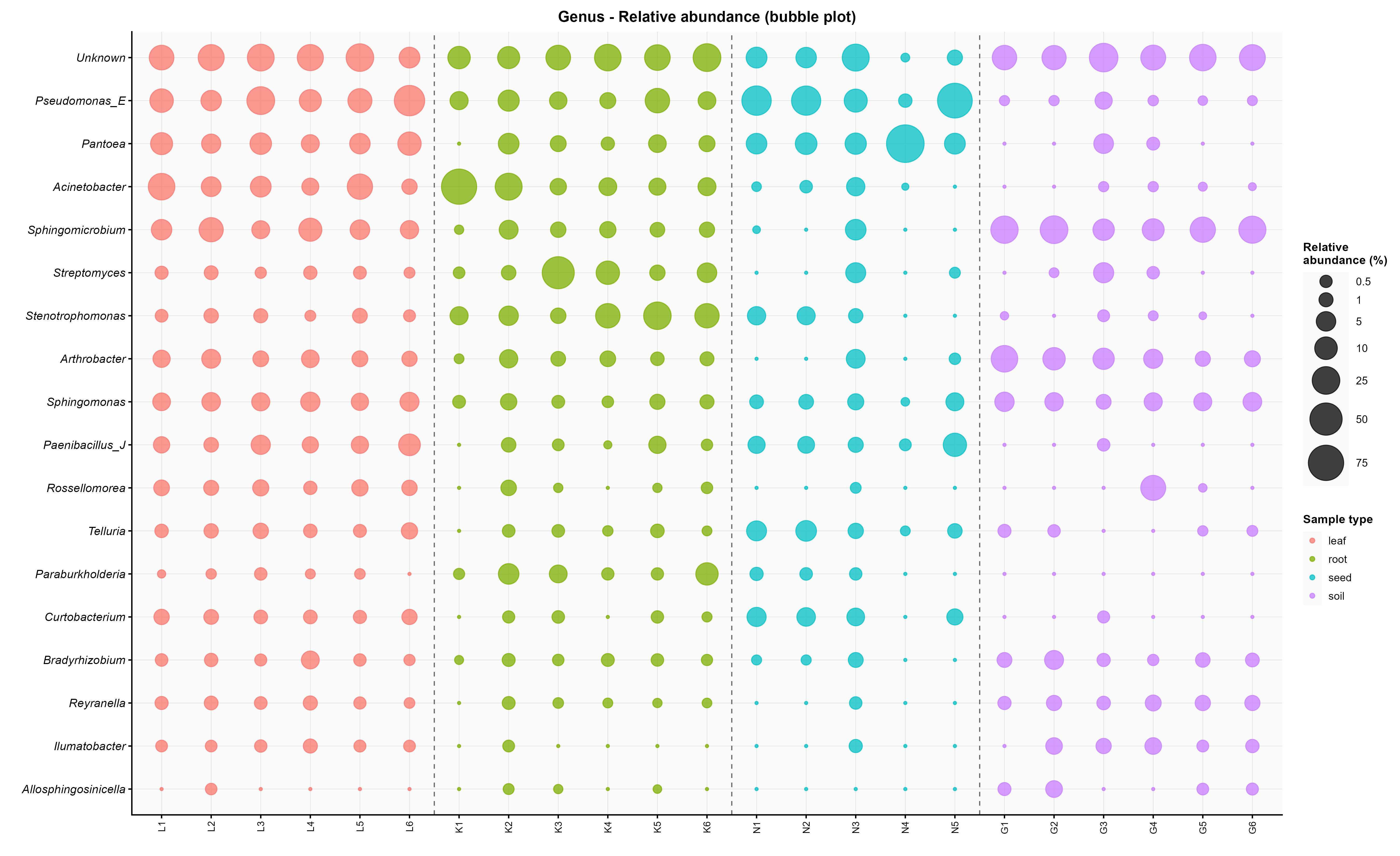

### 2_relative_bubble_Order.png

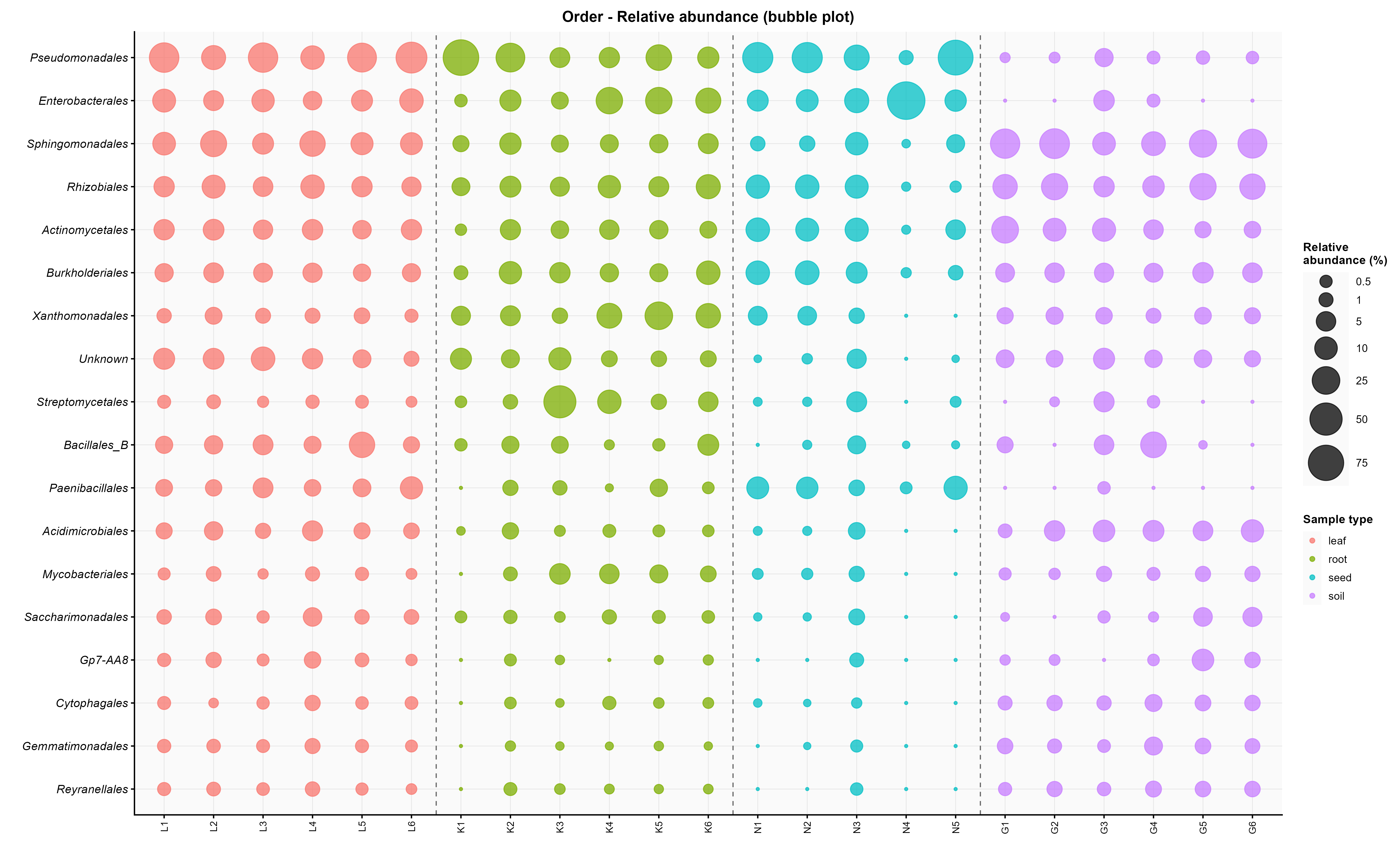

### 2_relative_bubble_Phylum.png

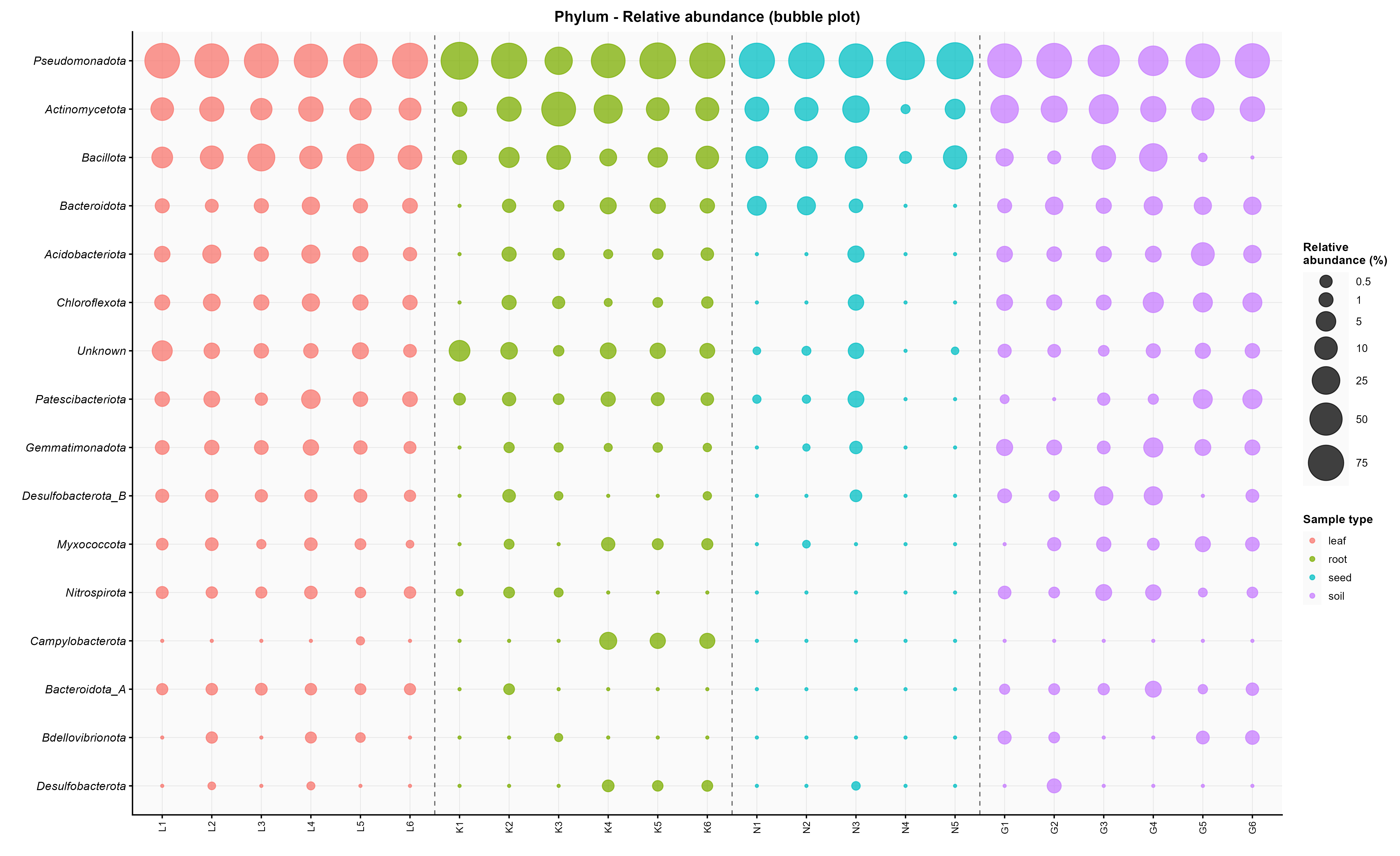

### 3_absolute_bubble_Class.png

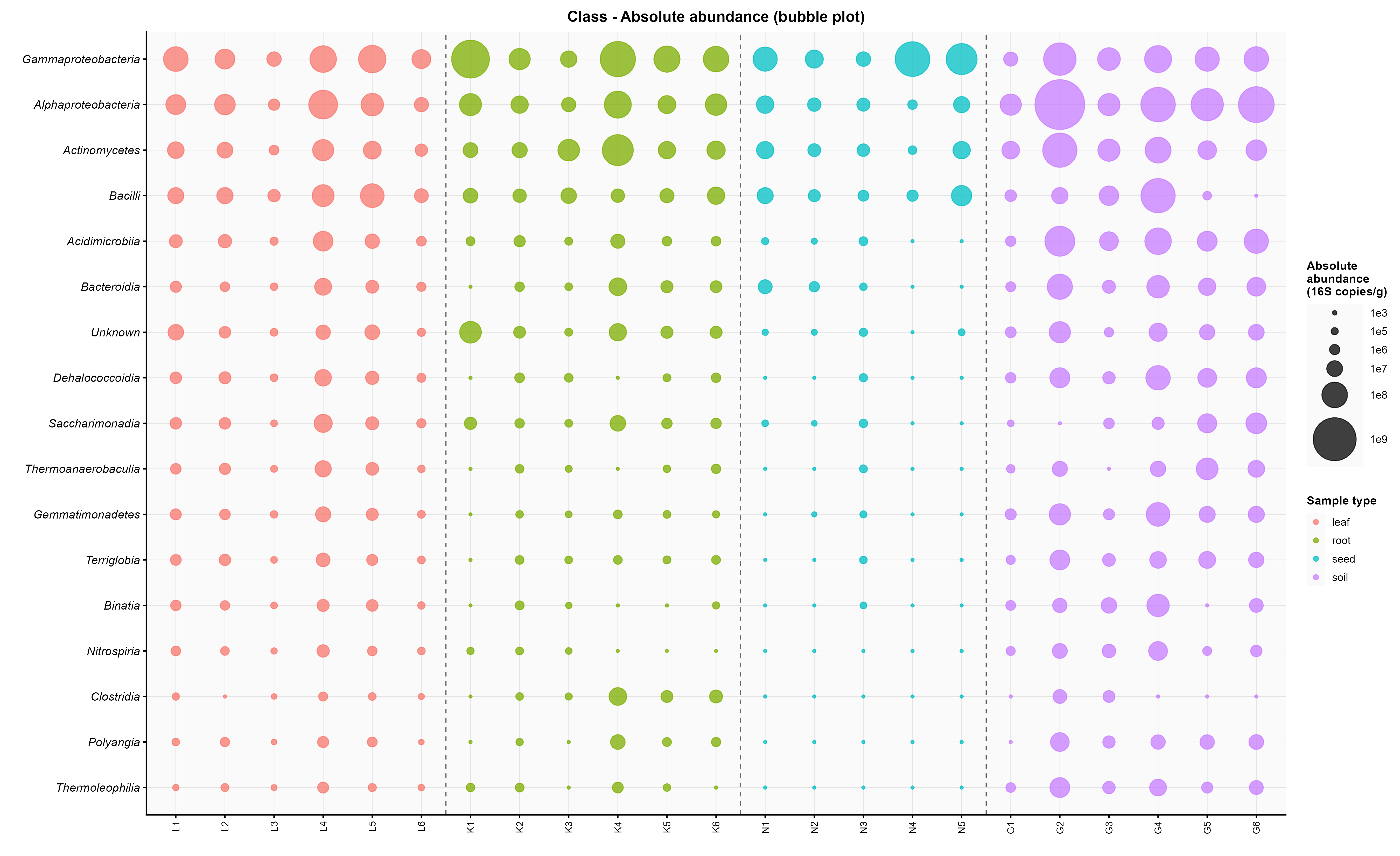

### 3_absolute_bubble_Family.png

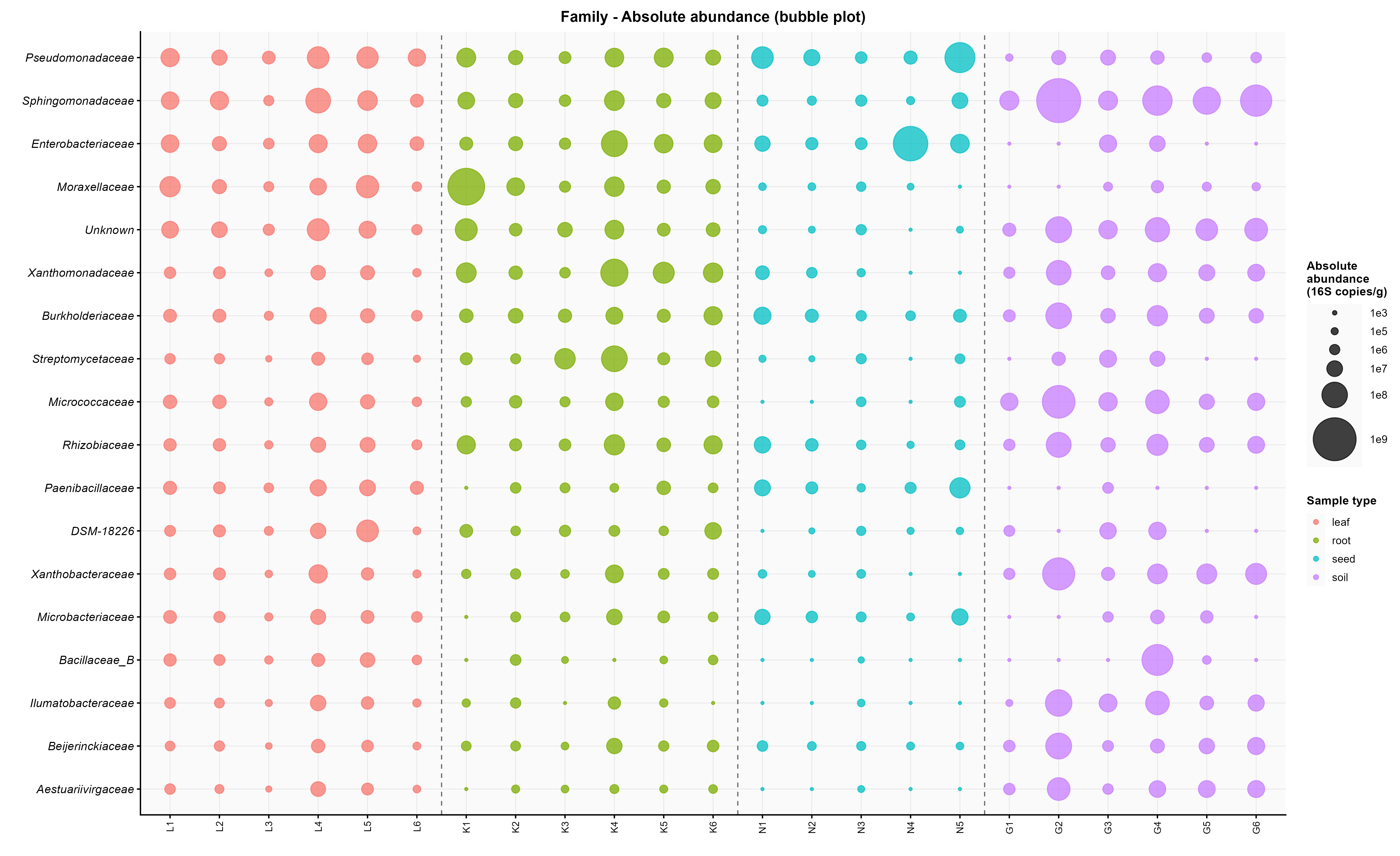

### 3_absolute_bubble_Genus.png

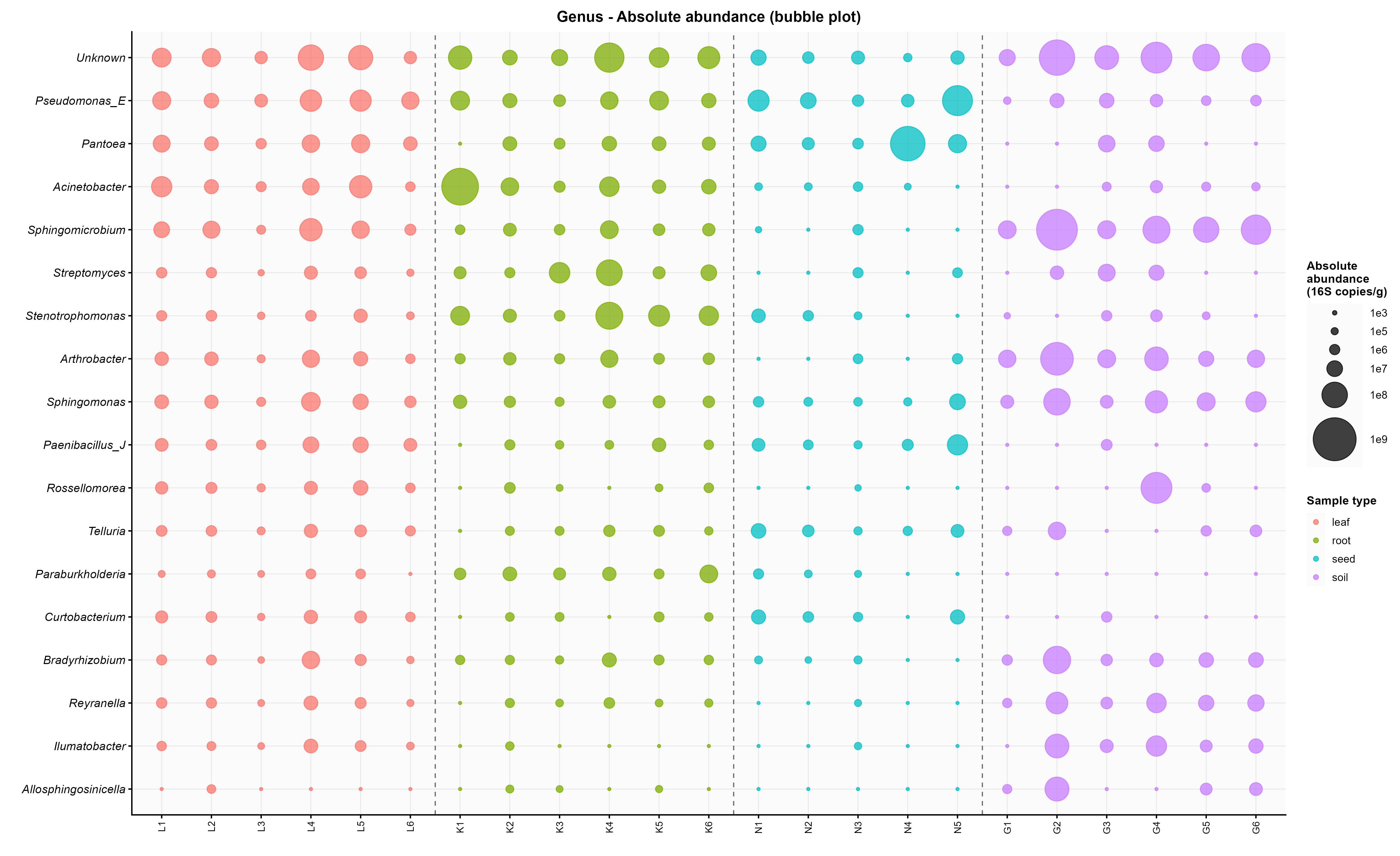

### 3_absolute_bubble_Order.png

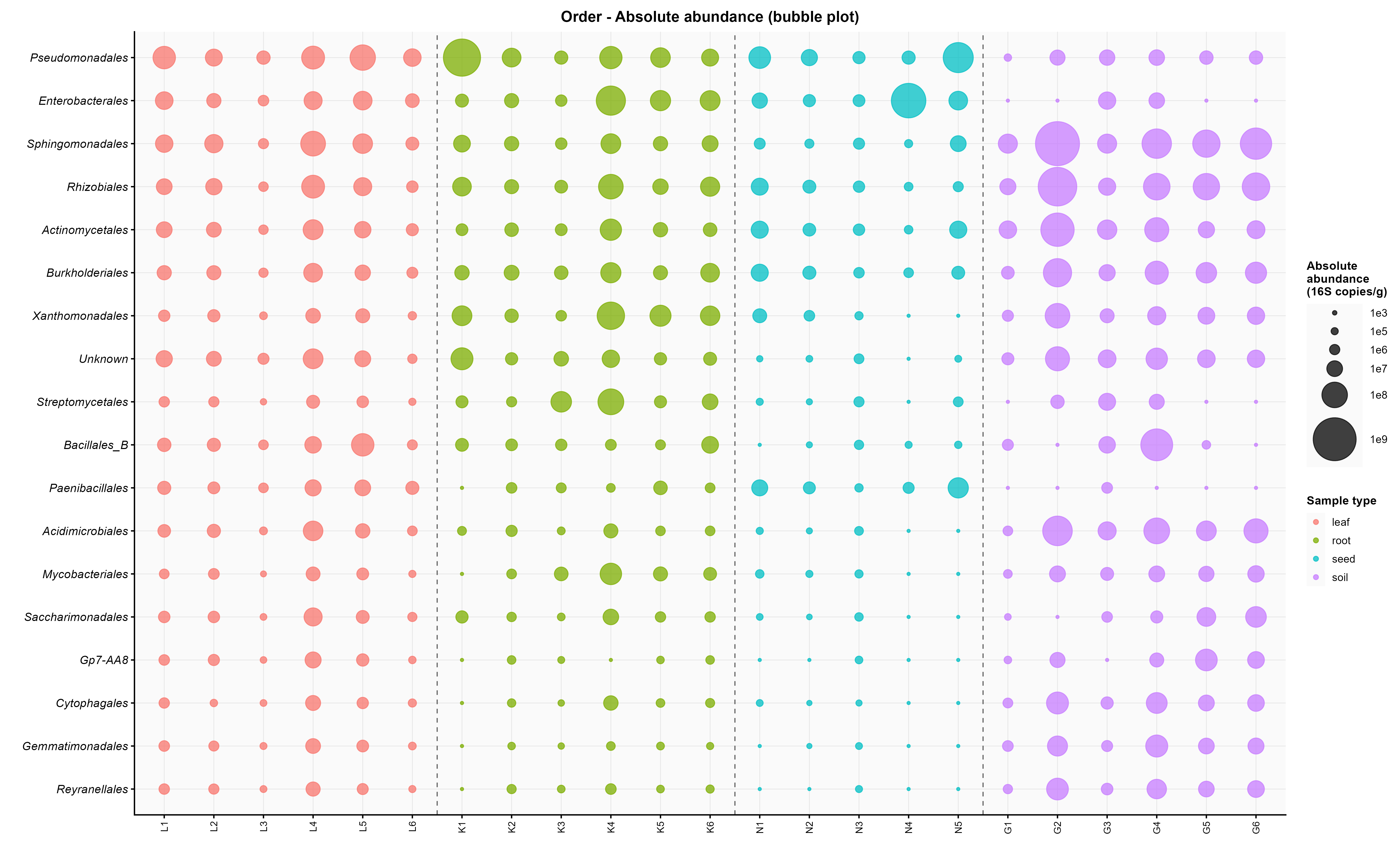

### 3_absolute_bubble_Phylum.png

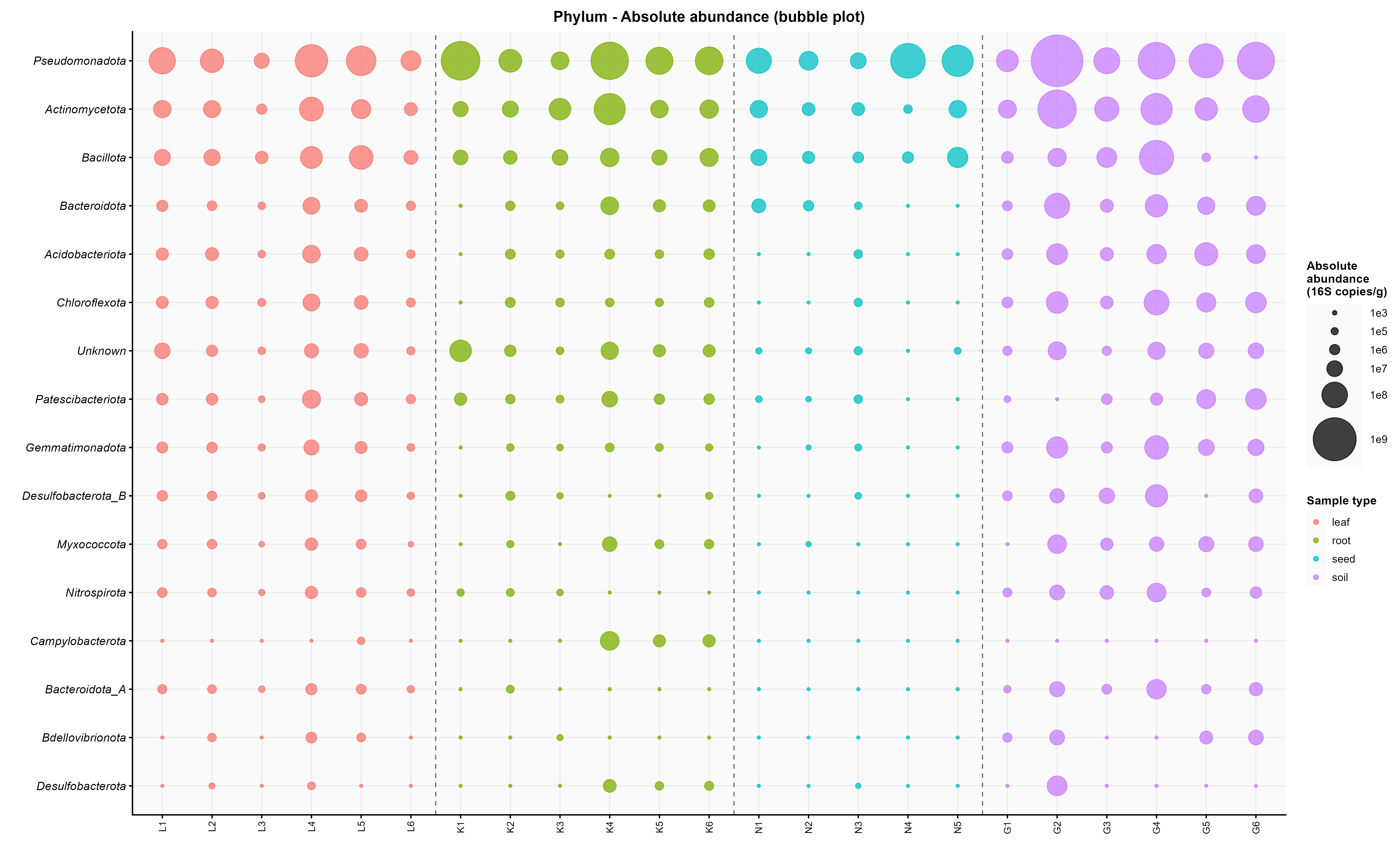
